# A bacterial effector counteracts host autophagy by promoting degradation of an autophagy component

**DOI:** 10.1101/2021.03.17.435853

**Authors:** Jia Xuan Leong, Margot Raffeiner, Daniela Spinti, Gautier Langin, Mirita Franz-Wachtel, Andrew R. Guzman, Jung-Gun Kim, Pooja Pandey, Alyona E. Minina, Boris Macek, Anders Hafrén, Tolga O. Bozkurt, Mary Beth Mudgett, Frederik Börnke, Daniel Hofius, Suayib Üstün

## Abstract

Beyond its role in cellular homeostasis, autophagy plays anti- and pro-microbial roles in host-microbe interactions, both in animals and plants. One prominent role of anti-microbial autophagy is to degrade intracellular pathogens or microbial molecules, in a process termed xenophagy. Consequently, microbes evolved mechanisms to hijack or modulate autophagy to escape elimination. Although well-described in animals, the extent to which xenophagy contributes to plant-bacteria interactions remains unknown. Here, we provide evidence that *Xanthomonas campestris* pv. *vesicatoria (Xcv)* suppresses host autophagy by utilizing type-III effector XopL. XopL interacts with and degrades the autophagy component SH3P2 via its E3 ligase activity to promote infection. Intriguingly, XopL is targeted for degradation by defense-related selective autophagy mediated by NBR1/Joka2, revealing a complex antagonistic interplay between XopL and the host autophagy machinery. Our results implicate plant antimicrobial autophagy in depletion of a bacterial virulence factor and unravels an unprecedented pathogen strategy to counteract defense-related autophagy.

## Introduction

Eukaryotic cells react dynamically to external and internal stimuli by adjusting their proteome. This requires a stringent regulation of protein homeostasis which is achieved in large part by regulated protein degradation. Cellular degradation machineries including the proteasome and autophagy maintain protein homeostasis by recycling unwanted or dysfunctional proteins (Pohl & Dikic, 2019). While the proteasome degrades short-lived proteins or mis-folded proteins, autophagy can remove larger protein complexes, insoluble aggregates, entire organelles as well as pathogens. Under normal conditions, both degradation pathways are critical for cellular housekeeping functions, while under stress conditions they facilitate the reorganization of the proteome to adapt to a changing environment (Marshall & Vierstra, 2018).

Regulated proteolytic degradation by proteasome has been identified as an essential component of immunity influencing the outcome of host-microbe interactions across kingdoms (Adams & Spoel, 2018; Hu & Sun, 2016). In the recent years, autophagy has also emerged as a central player in immunity and disease in humans and plants (Germic *et al*, 2019; Leary *et al*, 2019; Levine *et al*, 2011; Üstün *et al*, 2017; Yang & Klionsky, 2020). In mammals, autophagy has various connections to several diseases, regulating cell death and innate immunity (Germic *et al*, 2019; Yang & Klionsky, 2020). Dual roles have also been ascribed to autophagy in host-bacteria interactions (Mostowy, 2013). While some bacterial pathogens recruit the autophagy machinery in order to create a replicative niche (pro-bacterial autophagy), anti-bacterial autophagy removes bacterial intruders to limit pathogen infection (Huang & Brumell, 2014). The elimination of bacteria is a selective autophagy response, termed xenophagy (Gomes & Dikic, 2014). In this process, bacterial pathogens such as *Salmonella* and *Shigella* are degraded by autophagy through a ubiquitin-dependent mechanism (Dupont *et al*, 2009; van Wijk *et al*, 2012). This demonstrates that autophagy is not only a largely unspecific (“bulk”) catabolic and recycling process, as increasing evidence now indicates that autophagy also acts as a selective mechanism to degrade protein aggregates, organelles and pathogens. Selectivity is mediated by autophagy receptors, of which p62 and NBR1 play key roles in controlling pathogenic infection in mammals (Gomes & Dikic, 2014). Both autophagy receptors can bind to ubiquitinated bacteria, and degrade them, through their ability to bind autophagosome-associated ATG8 proteins (Gomes & Dikic, 2014).

It is known that type-III effector (T3E) proteins of plant pathogenic bacteria are present in the host cell while bacteria reside in the extracellular space. These effectors are able to manipulate host defense responses for the benefit of the pathogen (Khan *et al*, 2018). Very recently, it has been shown that microbial effectors perturb or hijack degradation machineries to attenuate plant immune reactions (Banfield, 2015; Langin *et al*, 2020). For instance, *Pst* activates autophagy via the action of the T3E HopM1 to degrade the proteasome and suppress its function in a process termed proteaphagy (Üstün *et al*, 2018; Üstün *et al*, 2016). Although this process can be categorized as a pro-bacterial role of autophagy, NBR1-mediated anti-bacterial autophagy seems to restrict lesion formation and pathogenicity of *Pst* (Üstün & Hofius, 2018). The dual role of autophagy in plant-bacteria interactions is further confirmed by findings that certain effectors are also able to suppress autophagy responses, although the understanding of the exact molecular mechanisms are still very limited (Lal *et al*, 2020). In addition, plant NBR1, which is also referred as Joka2 in solanaceous species, is able to restrict growth and disease progression of the plant pathogenic oomycete *Phytophthora infestans* (Dagdas et al, 2016; Dagdas et al, 2018). Recently, plant NBR1-mediated xenophagy was described to remove intracellular viral proteins (Hafrén *et al*, 2017; Hafrén *et al*, 2018). These studies demonstrated that, similar to that in mammals, plant NBR1 participates in xenophagy by degrading viral proteins. Given the fact that plant pathogenic bacteria reside in the extracellular space and the presence of T3Es inside the host cell it is not known whether NBR1-mediated xenophagy might play a role in plant-microbe interactions by targeting intracellular T3Es.

Like *Pst*, *Xanthomonas campestris* pv*. vesicatoria (Xcv)* is another well-studied hemi-biotrophic bacterium, causing disease on tomato and pepper plants (Timilsina *et al*, 2020). Mounting evidence has been established that *Xcv* and its T3Es exploit plant ubiquitin- and ubiquitin-like pathways (Buttner, 2016; Üstün & Börnke, 2014). While the role of the proteasome system in *Xanthomonas* infections is well understood, little is known about how autophagy shapes the outcome of *Xanthomonas*-host interactions. Recent findings in the cassava-*Xanthomonas axonopodis* pv. *manihotis* (*Xam*) model suggest that autophagy has an anti-bacterial role (Yan et al, 2017; Zeng et al, 2018). However, our current understanding about how T3Es might modulate and regulate this response is very limited. Are there similar mechanisms operating in pro- and antibacterial roles across different pathogenic bacteria? Do plants utilize xenophagy as an anti-bacterial mechanism to degrade pathogenic components in plant-bacteria interactions?

To address these questions, we performed a mechanistic analysis of the interplay of plant defence-related autophagy and *Xcv* pathogenesis. Here, we provide evidence that NBR1/Joka2 degrades T3E XopL, which in turn is used by *<u>Xcv</u>* to block autophagy via its E3 ligase activity by degrading autophagy component SH3P2 in a proteasome-dependent manner. This prevents T3E XopL from being targeted for degradation by the selective autophagy receptor NBR1/Joka2. We show that specificity in autophagy pathways play a role during plant-pathogen interactions.

## Results

### Xanthomonas blocks autophagy in an effector-dependent manner to promote pathogenicity

Given the prominent role of autophagy in host-microbe interactions, we investigated autophagic response after *Xanthomonas* infection. To this end we used the model plant *Nicotiana benthamiana*, since methods such as Agrobacterium-mediated transient expression, virus-induced gene silencing (VIGS), and autophagy activity reporter assays are well-established and reproducible. Assays were conducted with a *Xcv* strain harboring a deletion in the T3E XopQ (*Xcv ΔxopQ)*, which is a host range determinant in *Nicotiana* species (Adlung *et al*, 2016), thus restoring the full virulence of *Xcv* in *Nicotiana benthamiana* in the absence of XopQ.

First, we monitored autophagosome formation using RFP-ATG8g which is a structural component of autophagosomes and is widely used to label these structures (19). We infected *N. benthamiana* plants transiently expressing RFP-ATG8g with *Xcv ΔxopQ* and monitored autophagosomal structures during Concanamycin A (ConA) treatment. ConA is an inhibitor of vacuolar acidification that blocks autophagic body degradation (Minina *et al*, 2018; Svenning *et al*, 2011). In the absence of ConA, *Xcv ΔxopQ* induced massive accumulation of autophagosome-like structures which could not be further enhanced by the presence of ConA (Fig. 1A). This suggests that *Xcv* blocks autophagic degradation *in planta*. To provide additional evidence that these structures are indeed autophagosomes, we imaged *N. benthamiana* plants which were silenced using VIGS for *ATG7*, a crucial component of the autophagy pathway, and transiently expressing GFP-ATG8e, and found that the structures that accumulated under Xcv infection after 6hpi or AZD8055 treatment, a compound known to induce autophagy, no longer accumulating (Fig. S1A). The induction of autophagosome formation and suppression of autophagic degradation prompted us to investigate host autophagy by immunoblotting for endogenous ATG8 and Joka2 in *N. benthamiana* (Dagdas *et al*, 2016; Svenning *et al*, 2011). We also used the previously described autophagy suppressor AIMp, a small peptide sequence derived from the *Phytophthora* PexRD54 effector (Pandey *et al*, 2021), as a positive control for autophagy suppression. Joka2 protein abundance increased during infection, to a small extent 1-day post-inoculation (dpi), and to a greater extent at 2 dpi, while ATG8 protein levels only slightly increased at 2 dpi (Fig. 1B). Protein accumulation seen during immunoblotting could be attributed to increased transcription and/or decreased degradation. Thus, to uncouple these effects we utilized a quantitative autophagy assay to measure autophagic degradation during infection. This assay is based on Agrobacterium-mediated transient expression of 35S promoter-driven *Renilla* luciferase (RLUC) fused to ATG8a (RLUC-ATG8a) or NBR1 (RLUC-NBR1), together with free *Firefly* luciferase (FLUC) which serves as an internal control for expression as it is not degraded with autophagy (Dauphinee *et al*, 2019; Üstün *et al*., 2018). The autophagy reporter assay revealed that *Xcv ΔxopQ* infection led to a significant increase of RLUC-ATG8a/FLUC and RLUC-NBR1/FLUC ratios, suggesting reduced autophagic turnover after 2dpi (Fig. 1C, Fig. S1B). Another indicator of impaired autophagy is the increased gene expression of the autophagic markers (Minina *et al*, 2018). Transcript levels of *Joka2*, *NbATG8-2.1/2* and *NbATG8-1.1/1.2* were significantly higher compared to mock infection at 2 dpi (Fig. 1D), with *NbATG8-2.1/2.2* showing an earlier increase than the two other genes and suggesting at a differential response of NbATG8 isoforms during *Xcv* infection. Taken together with previous results, accumulation of Joka2 protein levels at 1 dpi which was observed earlier than its induced gene expression at 2 dpi, as well as reduced autophagic turnover after 6 hpi using the autophagy reporter assay (Fig. S1B) strongly suggest that *Xcv* dampens autophagic flux. To assess the biological relevance of suppressed autophagic degradation during *Xcv* infection, we determined bacterial growth in *N. benthamiana roq1* plants which carry a mutation in *Recognition of XopQ 1 (Roq1)* that recognizes the effector XopQ to activate resistance to *Xcv* (Gantner *et al*, 2019; Schultink *et al*, 2017). In these plants we transiently expressed RFP-AIMp as an autophagy suppressor. At 6 dpi, *Xcv* growth was significantly elevated in *roq1* plants transiently expressing RFP-AIMp compared to empty vector (EV) control (Fig. 1E). The same trend was observed when *ATG7* was silenced using VIGS in *N. benthamiana,* as *ATG7* silencing rendered plants more susceptible to *Xcv ΔxopQ* at 6 dpi (Fig. S2).

**Figure 1:**
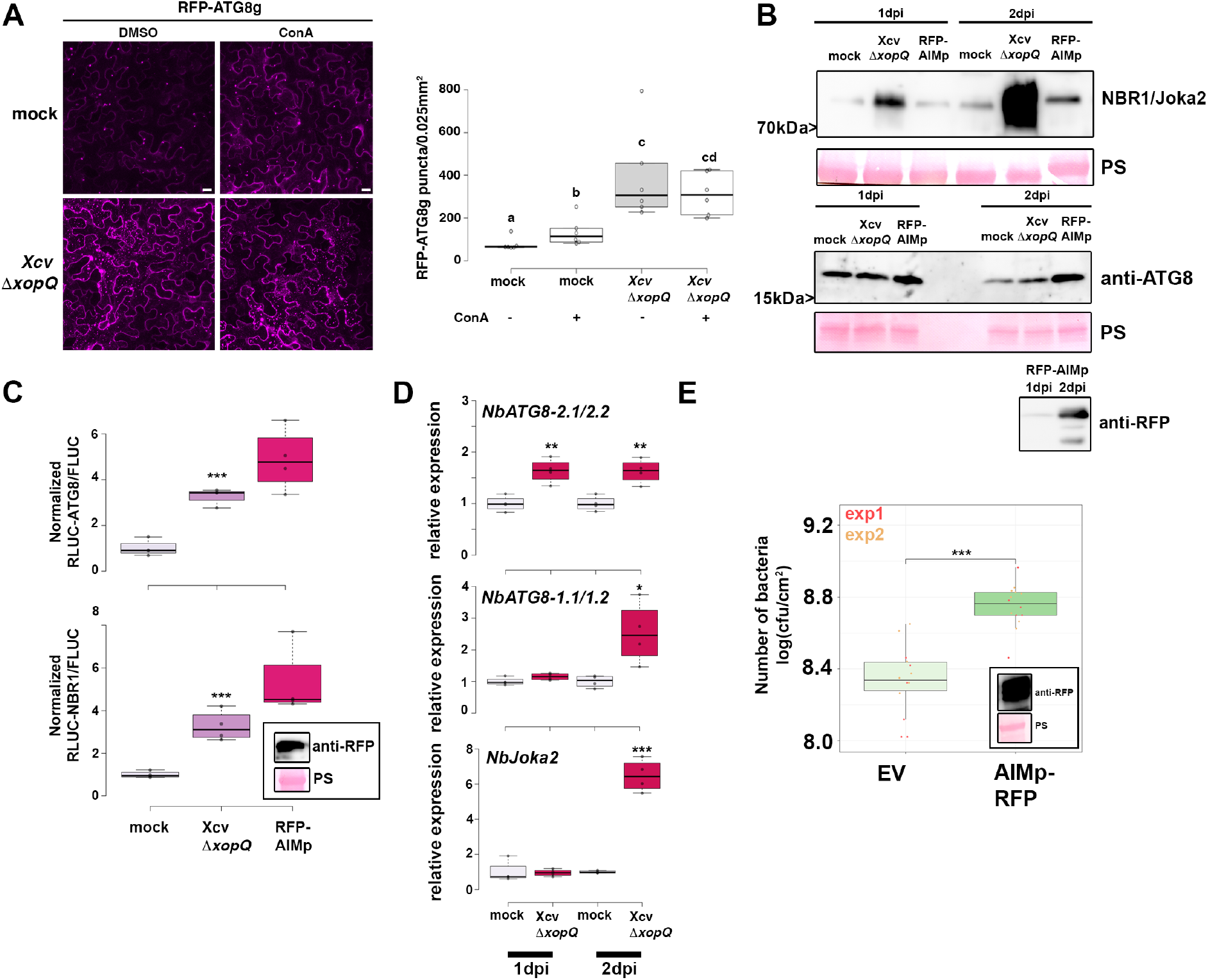
Xanthomonas blocks autophagy to enhance its pathogenicity. **(A)** RFP-ATG8g-labeled autophagosomes were quantified from plants infected with mock or Xcv *ΔxopQ* at 2 dpi in the presence or absence of ConA (bars = 20 μm). Puncta were calculated from z-stacks (15) of *n*=6 individuals using ImageJ. Data points are plotted as open circles. Different letters indicate statistically different groups (*P* < 0.05) as determined by one-way ANOVA. The experiment was repeated twice with similar results. **(B)** Immunoblot analysis of NBR1 and ATG8 protein levels in Xcv *ΔxopQ* or mock infected *N. benthamiana* plants at 1 and 2dpi. Agrobacterium-mediated transient expression of AIMp-RFP serves as a control for autophagy suppression. Ponceau Staining (PS) served as a loading control. The experiment was repeated three times with similar results. **(C)** RLUC-ATG8a or RLUC-NBR1 constructs were coexpressed with internal control FLUC in *N. benthamiana*. *Xcv ΔxopQ* was co-infiltrated with Agrobacteria containing the luciferase reporter constructs. Coexpression of RFP-AIMp serves as a control for autophagy inhibition. Expression of the latter was confirmed with western blot (inset). *Renilla* (RLUC) and *Firefly* (FLUC) luciferase activities were simultaneously measured in leaf extracts at 48 h post-infiltration using the dual-luciferase system (n=4). Statistical significance (\*\*\**P*<0.001) was revealed by Student’s *t*-test. The experiment was repeated more than 3 times with similar results. **(D)** RT-qPCR analysis of *NbATG8-1.1/1.2, NbATG8-2.1/2.2* and *NbJoka2* transcript levels upon challenge of *N. benthamiana* plants with Xcv *ΔxopQ* for 1 and 2 dpi compared to mock infected plants. Values represent expression relative to mock control of respective time point and were normalized to *Actin*. Statistical significance (\**P* < 0.05, \*\**P* < 0.01, *** *P* < 0.001) was revealed by Student’s *t*-test. **(E)** Bacterial density in leaves of *N. benthamiana roq1* infected with *Xcv* in the presence or absence autophagy suppressor AIMp-RFP. Leaves were syringe-infiltrated with OD_600_ = 0.0004, and colony-forming units were counted at 6 dpi. Compared to empty vector control (EV), AIMp expressing plants (n=6) harbour significantly more bacteria. Bacterial growth was repeated with the same result in 12 plants over two independent experiments. Red and yellow data points indicate independent repeats of the experiment. Statistical significance (\*\*\**P* < 0.001) was revealed by Student’s *t*-test. Expression of RFP-AIMp was verified at 6 dpi with an anti-RFP blot (inset).

Because T3Es were previously shown to modulate proteasome function and autophagy (Üstün et al, 2018; Üstün et al, 2016; Üstün *et al*, 2013), we analyzed host autophagy response to a nonpathogenic type-III secretion system (T3SS) mutant *Xcv ΔhrcN*, which is unable to drive secretion of T3Es (Lorenz & Buttner, 2009). In contrast to *Xcv ΔxopQ*, the T3SS-deficient mutant *Xcv ΔhrcN* did not alter the protein abundance of ATG8 and Joka2 (Fig. S3A), nor RLUC-ATG8a/FLUC or RLUC-NBR1/FLUC ratio (Fig. S3B) or transcript abundance of autophagy marker genes *NbJoka2* and *NbATG8-1.1/1.2* (Fig S3C). Together, these data support the model that *Xcv* blocks autophagy in a T3E-dependent manner to promote virulence.

### T3E XopL suppresses autophagy

To address which T3E(s) might manipulate autophagy, we screened for *Xcv* effectors XopJ, XopD and XopL which have known function in modulating proteolytic degradation pathways (Üstün & Börnke, 2014; Kim *et al*, 2013; Singer *et al*, 2013; Üstün *et al*, 2015; Üstün & Börnke, 2015) as well as XopS which has not been described to modulate degradation machineries. To this end, we used the quantitative dual-luciferase autophagy reporter assay. Transient expression of XopL, a previously characterized E3 ligase (Singer *et al*, 2013), and XopJ, an effector previously shown to inhibit host proteasome, led to a significant increase in RLUC-ATG8a/FLUC and RLUC-NBR1/FLUC ratio (Fig. 2A, S4A), which was consistent across multiple experiments. In contrast, transient expression of XopD and XopS had no evident effect on autophagic degradation (Fig. S4A). We chose to study XopL further, as XopJ was previously shown to inhibit host proteasome (Üstün *et al*, 2013), which may result in modulation of autophagy as shown by the effect of treatment with a proteasome inhibitor MG132 (Fig. S4B). Performing immunoblot analysis of ATG8 protein levels in *N. benthamiana* leaves, we found that transient expression of XopL resulted in an accumulation of NBR1 and ATG8 proteins at 2 dpi (Fig. 2B). While this was also consistent with elevated gene expression of ATG8, NBR1/Joka2 expression was only induced at 1 dpi but not 2 dpi upon XopL expression (Fig. S4C). Transient expression of the autophagy inhibitor AIMp showed similar expression trends (Fig. S4C). Treatment with ConA revealed that ATG8 levels could not be further enhanced when XopL was expressed (Fig. 2C), providing strong evidence that XopL inhibits autophagic turnover. We note that XopL accumulates under ConA treatment (Fig. 2C), which suggests that XopL is also subject to autophagic degradation.

**Figure 2:**
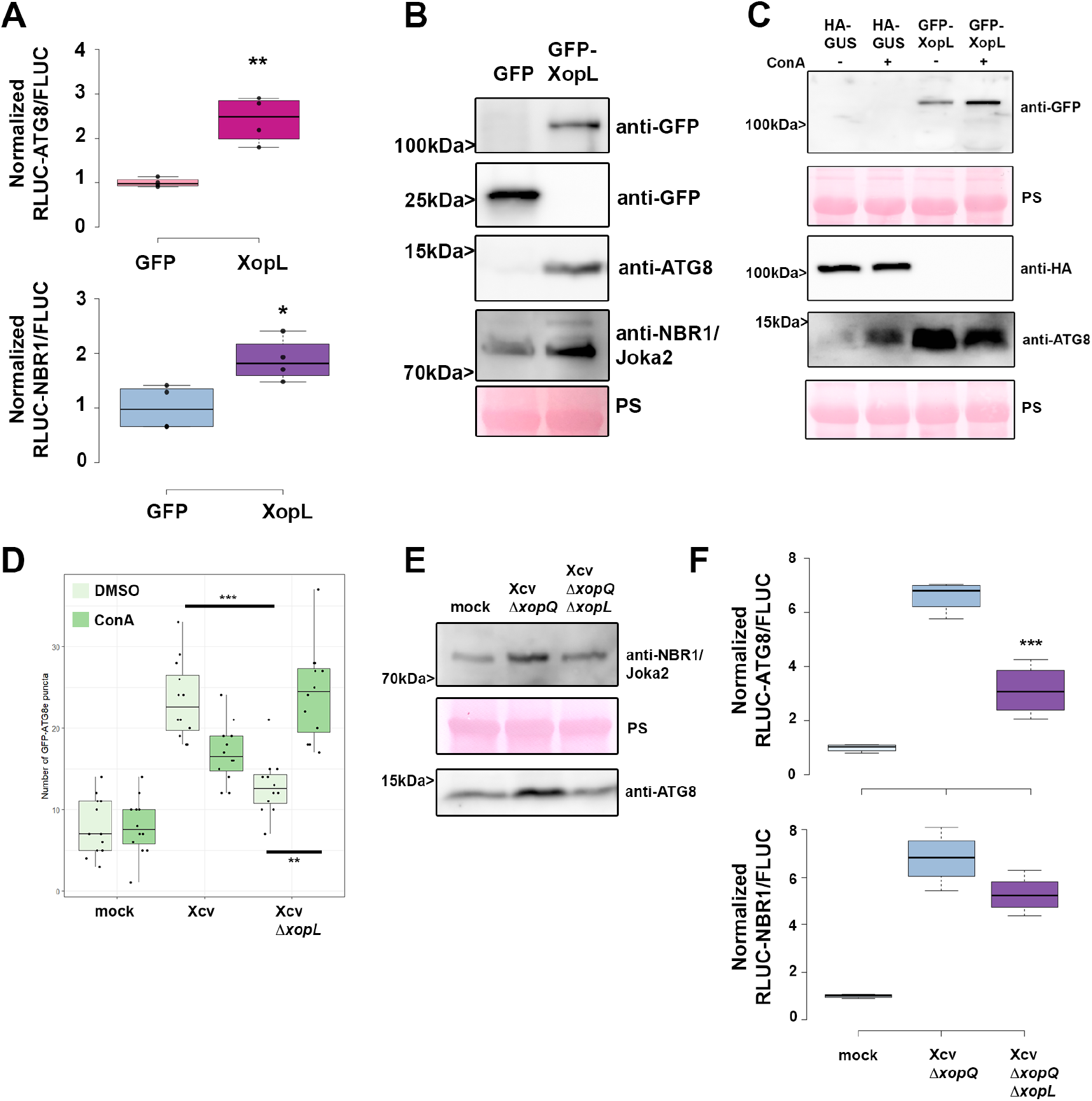
Xanthomonas T3E XopL is suppressing autophagy. **(A)** RLUC-ATG8a or RLUC-NBR1 constructs were coexpressed with internal control FLUC in *N. benthamiana*. XopL or GFP constructs were co-infiltrated. RLUC and FLUC signals were simultaneously measured in leaf extracts at 48 h post-infiltration using the dual-luciferase system. Values represent the ratio of RLUC-ATG8a and FLUC activities to the mean of control (n=4). Statistical significance (*P* < 0.01) was shown by Student’s *t*-test. The experiment was repeated more than 3 times by with similar results. **(B)** Immunoblot analysis of NBR1 and ATG8 protein levels in *N. benthamiana* plants transiently expressing GFP-XopL or GFP control at 2dpi verified with an anti-GFP antibody. Ponceau Staining (PS) served as a loading control. The experiment was repeated at least three times with similar results. **(C)** Immunoblot analysis of NBR1 and ATG8 protein levels in *N. benthamiana* plants transiently expressing XopL or GUS control at 2dpi after ConA or DMSO treatment. Expression of GFP-XopL was verified with an anti-GFP antibody, while expression of GUS-HA was confirmed with an anti-HA antibody. Ponceau Staining (PS) served as a loading control. The experiment was repeated twice with similar results. **(D)** GFP-ATG8g-labeled autophagosomes were quantified from plants infected with *Xcv ΔxopQ* or *Xcv ΔxopQ ΔxopL* at 2 dpi in the presence or absence of ConA. Puncta were calculated from z-stacks (15) of *n*=12 individuals using ImageJ. Statistical significance (** *P* < 0.01, *** *P* < 0.001) was determined by one way ANOVA. The experiment was repeated twice with similar results. **(E)** Immunoblot analysis of NBR1 and ATG8 protein levels in Xcv *ΔxopQ,* Xcv *ΔxopQ ΔxopL* or mock infected *N. benthamiana* plants at 2dpi. Ponceau Staining (PS) served as a loading control. The experiment was repeated twice with similar results. **(F)** RLUC-ATG8a or RLUC-NBR1 constructs were coexpressed with internal control FLUC in *N. benthamiana*. *Xcv ΔxopQ and Xcv ΔxopQ ΔxopL* were co-infiltrated with Agrobacteria containing the respective constructs. RLUC and FLUC activities were simultaneously measured in leaf extracts at 48 h post-infiltration using the dual-luciferase system. Statistical significance comparing *Xcv ΔxopQ and Xcv ΔxopQ ΔxopL* values (\*\*\**P* < 0.001) was revealed by Student’s *t*-test. The experiment was repeated 3 times with similar results.

To validate that XopL also acts as an autophagy suppressor during *Xcv* infection we constructed a *xopL* deletion mutant in *Xcv* WT and *Xcv ΔxopQ* backgrounds. *Xcv ΔxopL* displayed reduced growth and symptom development upon infection of tomato plants and the same, but to a lesser extent, was observed for *Xcv ΔxopL* in *N. benthamiana* (Fig S5A-E), demonstrating that XopL has a role during infection. Monitoring autophagosome-like structures in leaves transiently expressing GFP-ATG8e revealed that tissue infected with *Xcv ΔxopL* induced fewer GFP-ATG8e puncta than *Xcv* in the absence of ConA (Fig 2D, Fig S6A). Addition of ConA increased GFP-ATG8e puncta in leaves infected with *Xcv ΔxopL* but not in *Xcv*, indicating that XopL has a role in dampening autophagy during infection (Fig. 2D). By analyzing ATG8 and NBR1 protein levels, we also verified that XopL partially contributes to ATG8 and NBR1/Joka2 accumulation (Fig. 2E), supporting the notion that XopL suppresses autophagic degradation. We confirmed this when we monitored RFP-Joka2 in transiently expressing *N. benthamiana roq1* plants after Xcv infection. Infection with *Xcv* resulted in an induction of NBR1/Joka2 bodies in comparison to *Xcv ΔxopL* infected leaves (Fig. S6B). Utilizing the quantitative dual-luciferase autophagy assay, we show that *Xcv ΔxopQ ΔxopL* was unable to suppress autophagy to levels observed in tissues infected with *Xcv ΔxopQ* levels, both at 2dpi (Fig. 2F), indicating that XopL has a major impact on autophagy during infection. However, *Xcv ΔxopQ ΔxopL* still leads to a slight increase in both RLUC-ATG8a/FLUC and RLUC-NBR1/FLUC ratios, suggesting that *Xcv* possesses another T3E with a redundant function.

To analyze whether XopL has similar functions in other plant species, we generated transgenic *Arabidopsis thaliana* lines expressing GFP-XopL under the UBQ10 promoter. Similar to the results we obtained in *N. benthamiana*, GFP-XopL transgenic *A. thaliana* plants showed increased NBR1 protein abundance in the absence and presence of ConA treatment (Fig. S7A), suggesting a block of NBR1 turnover. The early senescence phenotype of transgenic lines expressing XopL (Fig. S7B) and elevated gene expression of *ATG8a* and *NBR1* is indicative of altered autophagy activity (Fig. S7D). Imaging with confocal microscopy revealed that GFP-XopL localizes to punctate structures in *A. thaliana* leaf epidermal cells (Fig. S7C).

### XopL interacts with and degrades the autophagy component SH3P2 contributing to Xcv virulence during infection

Previously, XopL was characterized as belonging to a novel class of E3 ligases and is capable of suppressing plant defense responses. There are no known plant targets of XopL (Singer *et al*, 2013), so we carried out a yeast-two hybrid (Y2H) screen using a cDNA library from tobacco (*Nicotiana tabacum*) to investigate whether XopL directly targets autophagy components to act as an autophagy suppressor. Our previous interactions studies indicate that the tobacco cDNA library is sufficient to identify host targets of *Xcv* T3Es that are conserved across different plant species, such as pepper, tomato and *A. thaliana* (Albers *et al*, 2019; Üstün *et al*., 2013; Üstün *et al*, 2014). One cDNA identified in the Y2H screening for XopL interacting proteins encoded a homologue of *A. thaliana* SH3P2, which has an amino acid identity of 74 % to the *N. tabacum* homologue (Fig S8A). Homologues are also present in *Nicotiana benthamiana* (NbSH3P2a and NbSH3P2b, 98% identity) and tomato (SlSH3P2, 96% to NtSH3P2) (Fig S8A, B). Direct interaction assays in yeast revealed that XopL is able to interact with SH3P2 from *N. tabacum* and *N. benthamiana* (Fig. 3A and Fig. S8C). SH3P2 from *A. thaliana* was previously identified as a novel autophagy component that interacts with ATG8 and binds to phosphatidylinositol 3-phosphate (PI3P) to regulate autophagosome formation, having also a potential role in late events of autophagosome biogenesis (Zhuang & Jiang, 2014; Zhuang *et al*, 2013). SH3P2 was also found to play a role in the recognition of ubiquitinated membrane proteins, and in targeting them to the endosomal sorting complexes required for transport (ESCRT) machinery (Nagel *et al*, 2017). We next sought to determine whether the interaction between XopL and SH3P2 occurs *in planta*. Due to expression problems of tobacco SH3P2 and also due to their high identity, we conducted further interaction studies with AtSH3P2. Using bimolecular fluorescence complementation (BiFC) and in *vivo* co-immunoprecipitation (co-IP), we found that XopL and AtSH3P2 interact in the plant cell, in small punctate structures resembling autophagosomes and also in larger structures (Fig. 3B and C; Supp Video 1). Additional *in vitro* co-IP data, using *E. coli* produced recombinant MBP-XopL and GST-*At*SH3P2, suggests that XopL and SH3P2 might directly interact with each other (Fig. S8D). Given the fact that SH3P2 from *A. thaliana* interacts with ATG8 and XopL localizes in puncta within plant cells (Zhuang *et al*, 2013; Erickson *et al*, 2018), we assessed whether XopL co-localizes with autophagosomes *in planta*. We were able to identify that transient expression of GFP-XopL in *N. benthamiana* with the autophagosome markers RFP-ATG8e and SH3P2-RFP resulted in co-localization (Fig. 3D). SH3P2 also co-localized with ATG8e upon transient co-expression in *N. benthamiana* (Fig. Fig.3D). This further supports the idea that XopL is functioning in the autophagy pathway by associating with these components *in planta*. Since XopL possesses E3 ligase activity, we next sought to investigate whether XopL might destabilize SH3P2 via ubiquitination, and thereby block autophagic degradation. Indeed, *in planta* transient co-expression of GFP-XopL and AtSH3P2-GFP resulted in a reduction in the latter’s protein abundance in *N. benthamiana* (Fig. 3E).

**Figure 3:**
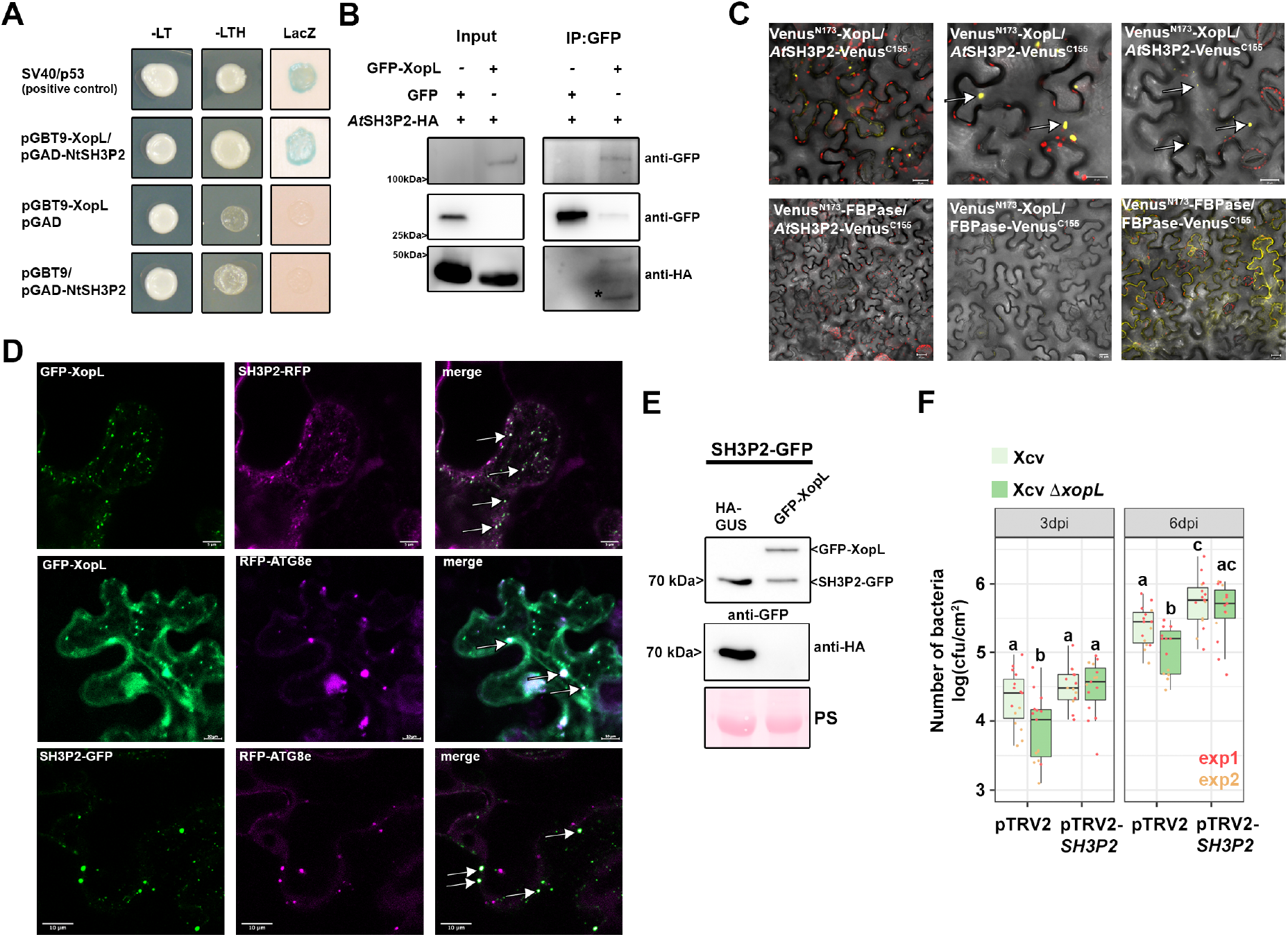
XopL interacts with and degrades SH3P2 and to boost *Xcv* virulence. **(A)** Interaction of XopL with SH3P2 in yeast two-hybrid assays. XopL fused to the GAL4 DNA-binding domain was expressed in combination with SH3P2 fused to the GAL4 activation domain (AD) in yeast strain Y190. Cells were grown on selective media before a LacZ filter assay was performed. pSV40/p53 served as positive control, while the empty AD or BD vector served as negative control. NtSH3P2 = *Nicotiana tabacum* SH3P2. –LT = yeast growth on medium without Leu and Trp, –HLT = yeast growth on medium lacking His, Leu, and Trp, indicating expression of the HIS3 reporter gene. LacZ, activity of the lacZ reporter gene. **(B)** Coimmunoprecipitation of GFP-XopL with AtSH3P2-HA. GFP-XopL or GFP were transiently coexpressed with AtSH3P2-HA in leaves of *N. benthamiana*. After 48 h, total proteins (Input) were subjected to immunoprecipitation (IP) with GFP-Trap beads, followed by immunoblot analysis using either anti-GFP or anti-HA antibodies. AtSH3P2 = *Arabidopsis thaliana* SH3P2. Two repetitions with similar results have been conducted. **(C)** Visualization of protein interactions *in planta* by the bimolecular fluorescence complementation assay. Yellow fluorescent protein (YFP) confocal microscopy images show *Nicotiana benthamiana* leaf epidermal cells transiently expressing Venus^N173^-XopL in combination with AtSH3P2-Venus^C155^. A positive control showing the dimerization of fructose-1,6-bisphosphatase (FBPase) within the cytosol. The red structures indicate autofluorescence of chloroplasts. The combination of Venus^N173^-XopL with FBPase-Venus^C155^ or Venus^N173^-FBPase with AtSH3P2-Venus^C155^ do not induce YFP fluorescence and serve as negative controls. Bars = 20 μm. **(D)** Colocalization analysis of GFP-XopL with SH3P2-RFP, RFP-ATG8e and RFP-ATG8g in *N. benthamiana* leaves. Imaging was performed 2 d after transient expression and images represent single confocal planes from abaxial epidermal cells (scale bars = 20 μm and 10 μm, lower panel). White arrows indicate colocalization of GFP and RFP signals. The experiment was repeated twice with similar results. **(E)** Total proteins were extracted 48 hpi with *A. tumefaciens* harboring the respective GFP-XopL, HA-XopL and SH3P2-GFP expression constructs. SH3P2-GFP protein levels (lower band) were detected using an anti-GFP antibody. Expression of the XopL was verified using an anti-HA or anti-GFP antibody. Expression of GUS-HA served as a control. Ponceau S staining serves as a loading control. The experiment was repeated three times with similar results. **(F)** Growth of Xcv *and* Xcv *ΔxopL* strains in *roq1 N. benthamiana* plants silenced for SH3P2 (pTRV2-*SH3P2*) compared to control plants (pTRV2). Leaves were dip-inoculated with a bacteria suspension at OD_600_ = 0.2 and bacteria were quantified at 3 and 6 dpi. Red and yellow data points indicate experimental repeats. Different letters indicate statistically different groups (*P* < 0.05) as determined by one way ANOVA.

Previously, downregulation of SH3P2 in *A. thaliana* has been shown to reduce autophagic activity (Zhuang *et al*, 2013). However, the role of SH3P2 is still controversial, as another study identified that SH3P2 functions in clathrin-mediated endocytosis without having any obvious effects on dark-induced autophagy (Nagel *et al*, 2017). To shed light on the enigmatic and versatile function of SH3P2, we used VIGS in *N. benthamiana* targeting both endogenous isoforms *NbSH3P2a* and *b*. Silencing had no obvious phenotypic effect on plants, and silencing efficiency was assessed by qPCR (Fig. S9A and B). Subsequent immunoblot analysis revealed that in comparison to the pTRV2-GFP control, SH3P2 VIGS plants displayed accumulation of ATG8 protein levels, similar results to that reported by Zhuang et al. 2013 (Fig. S9C). To corroborate this finding, we transiently expressed GFP-ATG8e in control and silenced plants and monitored autophagosome formation upon AZD8055 treatment, a TOR inhibitor and autophagy activator. The number of autophagosomes increased upon AZD8055 treatment in both plants but were significantly less in SH3P2 VIGS plants when treated with ConA (Fig. S9D). This indicates that downregulation of SH3P2 in *N. benthamiana* impairs maturation of autophagosomes and hence autophagic degradation. Indeed, using confocal microscopy and GFP-ATG8e labelling, we observed aberrant autophagosomal structures in VIGS SH3P2 plants, that might explain why autophagy is not entirely functional anymore. These data suggest that SH3P2 might be required during later steps of autophagosome formation, as autophagosomal structures seems to be normal during autophagy induction with AZD8055. VIGS in *N. benthamiana roq1* plants and subsequent bacterial growth measurements with *Xcv* and *Xcv ΔxopL* revealed that pTRV2-*SH3P2* plants are more susceptible towards *Xcv* (Fig. 3F). Essentially, partially reduced growth of *Xcv ΔxopL* was rescued in SH3P2-silenced plants strengthening our findings that XopL acts on *SH3P2* to suppress host autophagy and promote disease.

### XopL mediates proteasomal degradation of SH3P2 via its E3 ligase activity

Our results so far suggest that XopL might manipulate autophagy by interacting and degrading the autophagy component SH3P2. Previous research on SH3P2 revealed that RNAi-mediated downregulation of AtSH3P2 affects the autophagy pathway (Zhuang *et al*, 2013). To understand how SH3P2 is degraded by XopL we analyzed the degradation mechanisms in more detail. Firstly, degradation of AtSH3P2 by XopL was dependent on a functional proteasome, as chemical inhibition of the proteasome with MG132 partially restored AtSH3P2-GFP protein levels (Fig. 4A). Changes in SH3P2 protein levels were due to post-transcriptional events, as gene expression of SH3P2 is rather induced upon transient expression of XopL in *N. benthamiana,* (Fig S10A). To assess whether the proteasome-mediated degradation of SH3P2 was directly mediated by XopL and its E3 ligase activity, we performed an *in vitro* ubiquitination assay. In the presence of all required components of the E1-E2-E3 system, we observed that GST-XopL ubiquitinated MBP-AtSH3P2, which is indicated by a laddering pattern, leading to larger sized molecular species of MBP-AtSH3P2, when probed with the anti-MBP antibody (Fig. 4B). To address whether E3 ligase activity of XopL is crucial in the SH3P2-dependent modulation of host autophagy, we employed the triple point mutant of XopL_H584A L585A G586E_ (hereafter referred to as XopL ΔE3), lacking E3 ligase activity (Singer *et al*, 2013; Erickson *et al*, 2018). Transient co-expression revealed that XopL requires its E3 ligase activity to trigger the degradation of AtSH3P2 in *N. benthamiana* (Fig. 4C). This was not due to an altered localization of XopL ΔE3, as it still co-localizes with AtSH3P2 upon transient expression in *N. benthamiana* (Fig. S11A). In addition, XopL ΔE3 was also unable to ubiquitinate SH3P2 in an *in vitro* ubiquitination assay (Fig. S11B). Consistent with its inability to degrade SH3P2 *in planta*, XopL ΔE3 did not lead to a suppression of autophagy responses in the quantitative luciferase autophagy assay and increase in ATG8 protein levels (Fig. 4D and E). XopLΔE3 is also more unstable than XopL WT, suggesting that its E3 ligase activity is crucial to maintaining its stability, likely through its function in subverting autophagy (Fig. 4F). Taken together, our findings support the notion that the E3 ligase activity of XopL as well as its ability to directly ubiquitinate and degrade AtSH3P2 promote suppression of autophagy.

**Figure 4.**
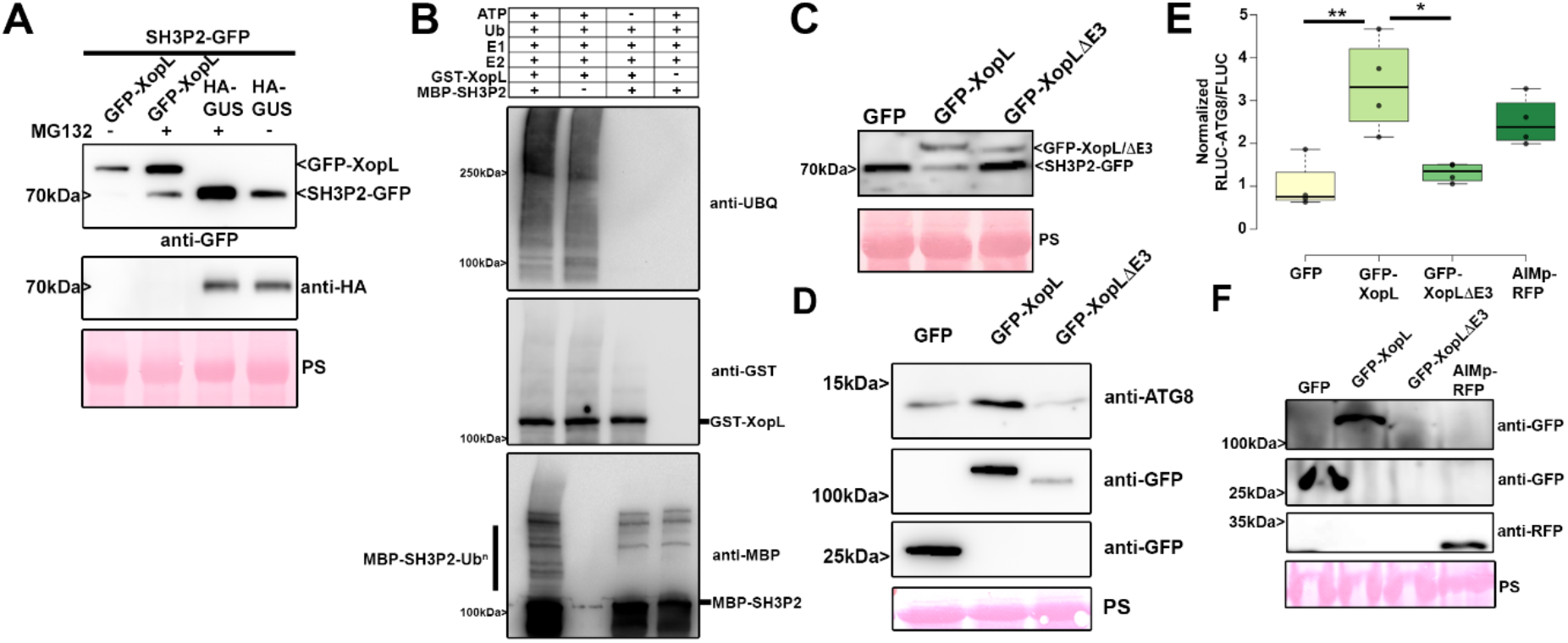
XopL mediates the proteasome degradation of SH3P2 via its E3 ligase activity. **(A)** SH3P2-GFP was transiently coexpressed together with GUS-HA and GFP-XopL in *N. benthamiana* using agroinfiltration. At 42 hpi, 200 μM MG132 was infiltrated into *A. tumefaciens*-inoculated leaves, and leaf material was collected 48 hpi. Expression of SH3P2-GFP (lower band) and GFP-XopL (upper band) was detected using an anti-GFP antibody. GUS-HA expression was confirmed with an anti-HA antibody. Ponceau S staining serves as a loading control. The experiment was repeated three times with similar results. **(B)** *In vitro* ubiquitination assay reveals ubiquitination of SH3P2 by XopL. GST-XopL, and MBP-SH3P2 were tested using the Arabidopsis His-AtUBA1 and His-AtUBC8. Lanes 2 to 4 are negative controls. Proteins were separated by SDS-PAGE and detected by immunoblotting using the indicated antibodies. The experiment was repeated twice with similar results. **(C)** SH3P2-GFP was transiently coexpressed together with GFP, GFP-XopL and GFP-XopL ΔE3 in *N. benthamiana* using agroinfiltration. GFP protein levels were detected with an anti-GFP antibody. Ponceau S staining serves as a loading control. The experiment was repeated three times with similar results. **(D)** Immunoblot analysis of ATG8 protein levels in *N. benthamiana* plants transiently expressing GFP-XopL, GFP-XopL ΔE3 or GFP control at 2dpi. Expression of binary constructs was verified with an anti-GFP antibody. Ponceau Staining (PS) served as a loading control. The experiment was repeated twice with similar results. **(E)** RLUC-ATG8a constructs were coexpressed with internal control FLUC in *N. benthamiana*. GFP-XopL, GFP-XopL ΔE3 or GFP control were co-infiltrated together with RLUC/FLUC mixture. *Renilla* and *Firefly* luciferase activities were simultaneously measured in leaf extracts at 48 hpi using the dual-luciferase system. Values represent the ratio of RLUC-ATG8a and FLUC activities (n=4). Statistical significance (* *P* < 0.5, ** *P* < 0.01) was revealed by Student’s *t*-test. The experiment was repeated 3 times. Expression of proteins was verified with indicated antibodies.

### NBR1/Joka2-mediated selective autophagy degrades ubiquitinated XopL

While we investigated the effect of *Xcv* and its T3E XopL on host autophagy, we noticed that NBR1/Joka2 responds at both transcript and protein levels during infection (Fig. 1B, 1D). We also observed that XopL protein accumulated under ConA treatment (Fig. 2C, Fig S5A), hinting that it was subject to autophagic degradation. Previous studies imply that NBR1/Joka2 mediates xenophagy by degrading viral particles, and that Joka2 is required for immunity against bacteria and *Phytophthora* (Üstün *et al*, 2018; Dagdas *et al*, 2016; Hafrén *et al*, 2018). However, the role of NBR1-mediated xenophagy in plant-*Phytophthora* and plant-bacteria interactions remains unknown. Plant NBR1 is a hybrid of the mammalian autophagy adaptors NBR1 and p62/SQSTM1 (Svenning *et al*, 2011). The latter was shown to mediate xenophagy of *Mycobacterium tuberculosis* (Mtb) by binding to the Mtb ubiquitin-binding protein Rv1468c and ubiquitin-coated *Salmonella enterica* in human cells (Chai *et al*, 2019; Zheng *et al*, 2009). To corroborate our previous finding that XopL accumulates when autophagy is blocked, we coexpressed GFP-XopL with autophagy inhibitor AIMp-RFP in *N. benthamiana leaves*. Immunoblot analysis revealed that indeed XopL also accumulates in the presence of autophagy inhibitor AIMp (Fig. 5A). As NBR1/Joka2 bodies were substantially induced during *Xcv* infection (Fig. S6), and block of autophagic degradation using ConA caused accumulation of GFP-XopL in vacuoles of *A. thaliana* (Fig. S12), we decided to examine whether XopL and NBR1/Joka2 associate *in planta*. Intriguingly, we discovered that XopL co-localizes with NBR1/Joka2 in puncta (Fig. 5B). Association of both proteins was determined using Förster resonance energy transfer by fluorescence lifetime imaging microscopy (FRET-FLIM). Only in the presence of RFP-XopL a significant reduction of 0.1 ns in the lifetime of the donor Joka2 was observed in comparison to co-expression of RFP or donor alone (Fig. 5C). This reduction of lifetime is similar to what was achieved with the positive control GFP-Joka2/RFP-ATG8e. These findings prompted us to investigate whether XopL might be targeted by NBR1/Joka2 for autophagic degradation, similar to insoluble ubiquitinated protein aggregates (Zhou *et al*, 2013). Performing pull-down experiments with GFP-XopL, we discovered that XopL associates with NBR1/Joka2 *in planta* (Fig. 5D), confirming the results of our FRET-FLIM and co-localization studies. To address whether E3 ligase activity of XopL is required for interaction, we employed XopL ΔE3, lacking E3 ligase activity. Co-IP experiments revealed that XopL ΔE3 was also able to pull-down NBR1/Joka2 after transient expression in *N. benthamiana* (Fig. 5E). This suggests that NBR1/Joka2 may not mediate the degradation of a complex containing XopL and its ubiquitinated target protein(s), but rather targets XopL for autophagic degradation. Given the fact that XopL is degraded by autophagy and associates with NBR1/Joka2, we next analyzed stability of XopL in Joka2-silenced *N. benthamiana* plants. Silencing of NBR1/Joka2 was confirmed by qPCR (Fig. S13A). Indeed, we could observe an increase in GFP-XopL protein abundance (Fig. 5F), but not for GFP (Fig. S13B), in pTRV2-Joka2 plants, arguing for a direct participation of NBR1/Joka2 in XopL turnover. To assess whether this might impact pathogenicity of *Xcv* we performed bacterial growth assays using the pTRV2-Joka2 plants. Increased growth at 3 dpi of *Xcv ΔxopQ* in *N. benthamiana* plants silenced for *Joka2* strengthened our finding that Joka2 is having anti-bacterial effects on *Xcv* early on during infection (Fig. 5G). As NBR1/Joka2 or p62 recognize their cargos by their ability to bind ubiquitinated substrates, we hypothesized that XopL might be ubiquitinated *in planta*. To test this, we transiently expressed GFP-XopL in *N. benthamiana* leaves and then immunoprecipitated GFP-XopL from leaf protein extracts. Ubiquitinated GFP-XopL was detected using immunoblotting. GFP-XopL, but not the GFP control, displayed polyubiquitination *in planta*, while GFP-XopL ΔE3 showed reduced polyubiquitination (Fig. 5H). To further confirm the ubiquitination of XopL, we purified total ubiquitinated proteins using the ubiquitin pan selector, which is based on a high-affinity single domain antibody that is covalently immobilized on cross-linked agarose beads. We detected a smear of high-molecular weight bands including the full-length XopL protein (Fig. 5I), which was strongly enhanced when we co-expressed with AIMp but absent in the GFP control (Fig. S14)

**Figure 5:**
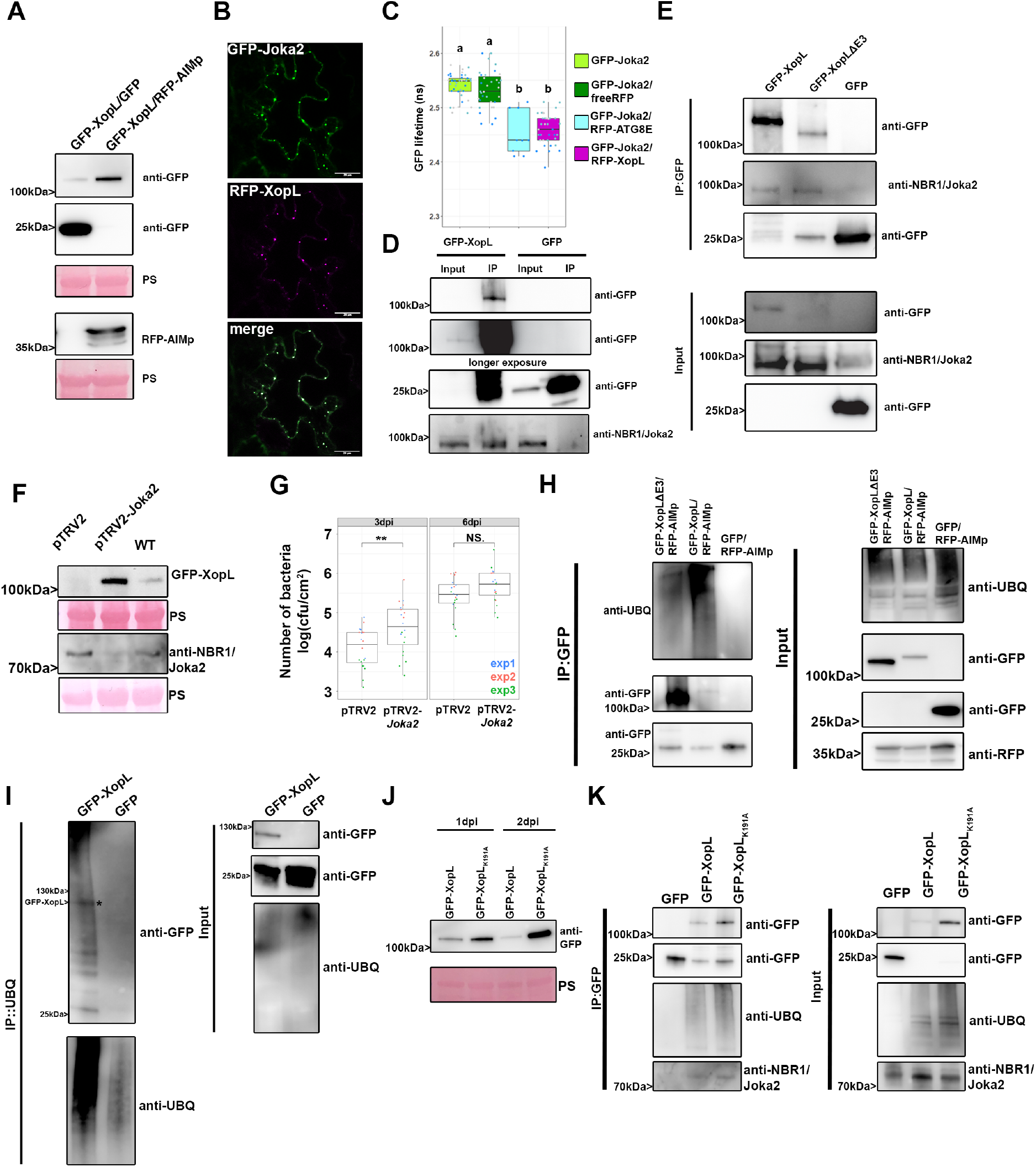
XopL is ubiquitinated in planta and degraded by NBR1-mediated selective autophagy. **(A)** GFP-XopL was coexpressed with GFP or AIMp-RFP. Proteins were separated by SDS-PAGE and detected by immunoblotting using the indicated antibodies. Ponceau Staining (PS) served as a loading control. The experiment was repeated three times with similar results. **(B)** Colocalization of GFP-XopL with RFP-Joka2 in *N. benthamiana* leaves. Imaging was performed 2 d after transient expression and images represent single confocal planes from abaxial epidermal cells (bars = 20 μm). The experiment was repeated twice with similar results. **(C)** FRET FLIM measurements of GFP-Joka2 and RFP-XopL in *N. benthamiana* leaves. The freeRFP construct served as a negative control and RFP-ATG8E (n = 9) as a positive control. Scattered points show individual data points, color indicates biological repeats. The lifetime (in ns) of GFP-Joka2 (donor, n = 41) was significantly reduced in the presence of RFP-XopL (n = 40) but not in the presence of freeRFP (n = 35). Significant differences were calculated using Wilcoxon rank sum test, with significantly different groups denoted by different letters. The experiment was repeated three times with similar results. **(D)** Immunoprecipitation (IP) of GFP-XopL reveals association with NBR1. Immunoblots of input and IP samples from *N. benthamiana* plants transiently expressing GFP or GFP-XopL were probed with anti-GFP and anti-NBR1 antibodies. **(E)** Immunoprecipitation (IP) of GFP-XopL and GFP-XopL ΔE3 reveals association with NBR1. Immunoblots of input and IP samples from *N. benthamiana* plants transiently expressing GFP, GFP-XopL and GFP-XopL ΔE3 were probed with anti-GFP and anti-NBR1 antibodies. **(F)** GFP-XopL was transiently expressed in pTRV2, pTRV2-Joka2 and *N. benthamiana* WT plants. Expression of binary constructs was verified with an anti-GFP antibody. Joka2 silencing was verified using an anti-NBR1 antibody. Ponceau Staining (PS) served as a loading control. The experiment was repeated twice with similar results. **(G)** Growth of Xcv *ΔxopQ* in *N. benthamiana* plants silenced for *Joka2* (pTRV2-Joka2) compared to control plants (pTRV2). Leaves were dip-inoculated with a bacteria suspension at OD_600_ = 0.2. and bacteria were quantified at 3 and 6 dpi. Red, blue, and green data points represent repeats of the experiments. Significant differences were calculated using Student’s *t-test* and are indicated by: **, P < 0.01. The experiment was repeated three times with similar trends. **(H)** GFP-XopL, GFP-XopL ΔE3 were transiently expressed in *N. benthamiana*. RFP-AIMp was co-infiltrated to stabilize both XopL variants. Samples were taken 48 hpi, and total proteins (Input) were subjected to immunoprecipitation (IP) with GFP-Trap beads, followed by immunoblot analysis of the precipitates using either anti-GFP or anti-ubiquitin antibodies. GFP served as a negative control. RFP-AIMp expression was verified by an anti-RFP antibody. The experiment was repeated three times with similar results. **(I)** GFP-XopL was transiently expressed in *N. benthamiana*. Samples were taken 48 hpi, and total proteins (Input) were subjected to immunoprecipitation (IP) with the ubiquitin pan selector, followed by immunoblot analysis of the precipitates using either anti-GFP or anti-ubiquitin antibodies. GFP served as a control. Asterisk indicates the GFP-XopL full-length protein. The experiment was repeated two times with similar results. **(J)** Immunoblot analysis GFP-XopL and GFP-XopL_K191A_ at 1 and 2 dpi using an anti-GFP antibody. Ponceau Staining (PS) served as a loading control. The experiment was repeated three times with similar results. **(K)** GFP-XopL and GFP-XopL_K191A_ were transiently expressed in *N. benthamiana*. Samples were taken 48 hpi, and total proteins (Input) were subjected to immunoprecipitation (IP) with GFP-Trap beads, followed by immunoblot analysis of the precipitates using either anti-GFP, anti-ubiquitin and anti-NBR1 antibodies. GFP served as a control. The experiment was repeated three times with similar results.

To identify ubiquitinated residues within the XopL protein, we immunoprecipitated GFP-XopL from *N. benthamiana* leaves transiently expressing GFP-XopL and performed mass spectrometry (MS) analysis. *In planta* MS analysis revealed one potential ubiquitination site at lysine 191 (K191) in the N-terminal part of XopL (Fig. S15A). For plant E3 ligases such as PUB22, it has been reported that its stability is dependent on its autoubiquitination activity (Furlan *et al*, 2017). Using MBP-XopL in an *in vitro* ubiquitination assay we confirmed self-ubiquitination (Fig. S15B), indicated by the presence of higher molecular weight bands probing with an anti-MBP antibody. Preforming LC-MS/MS we did detect the same ubiquitination (K191) site identified *in planta* when we analyzed *in vitro* ubiquitination samples containing GST-XopL (Fig. S15C). Additionally, an *in vitro* self-ubiquitination assay comparing XopL WT to its K191A mutant counterpart shows that K191A displays less intensity on its high molecular weight smear, suggesting that K191A is less ubiquitinated in this assay (Fig. S16A). This strongly argues for K191A being an autoubiquitination site of XopL. This is strengthened by our findings showing that the mutation of lysine 191 to alanine (K191A) rendered the XopL K191A more stable than WT XopL (Fig. 5J) without altering its subcellular localization (Fig. S16B). Changes in XopL vs. K191A protein levels were due to post-transcriptional events, as gene expression of XopL and XopL K191A are similar upon transient expression in *N. benthamiana* (Fig S16C) but did not abolish ubiquitination of XopL and association with autophagic machinery as shown by co-IP with NBR1/Joka2 (Fig. 5K, Fig. S16D). However, we cannot rule out trans-ubiquitination by plant E3 ligases, as XopL ΔE3 still interacts with NBR1 and is still ubiquitinated in *planta*. Immunoblot analysis also revealed that XopL ΔE3 is more unstable compared to XopL WT and degraded by autophagy, as it is not able to block autophagy (Fig S17). This might indicate the presence of additional ubiquitination sites in XopL. Taken together, our results suggest that XopL is ubiquitinated *in planta* and subjected to NBR1/Joka2-dependent selective autophagy.

## Discussion

Here, by studying *Xanthomonas*-host interactions, we revealed a complex multi-layered regulatory role of autophagy in plant immunity. We demonstrate that *Xanthomonas* can subvert plant autophagy which is achieved by the T3E XopL. Through its E3 ligase activity, XopL can ubiquitinate and degrade SH3P2, an autophagy component functioning in autophagosome biogenesis. This in turn dampens autophagy to boost the virulence of Xanthomonas. However, the same T3E is targeted by NBR1/Joka2-triggered selective autophagy/xenophagy which constitutes a novel form of plant defense mechanism (Fig. 6). The mutual targeting of pathogen effector XopL and plant protein SH3P2 unveils how different layers of regulation contribute to autophagy pathway specificity during host-bacteria interactions.

**Figure 6:**
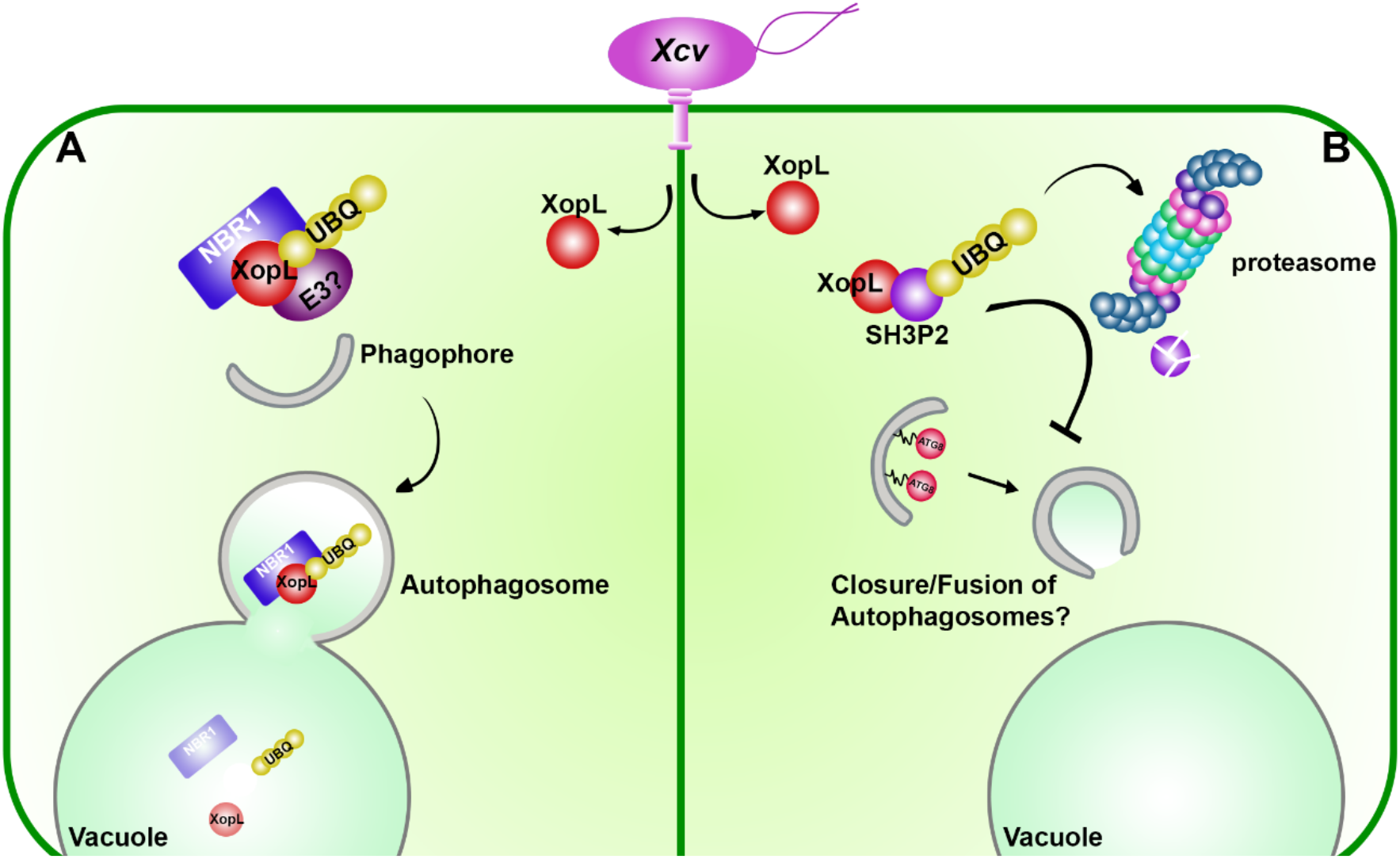
Model illustrating the function of XopL. **(A) Xenophagy of XopL:** Upon delivery of XopL in the plant XopL undergoes self-ubiquitination and possible ubiquitination by an unknown host E3 ligase. Joka2/NBR1 associates with XopL and triggers its degradation via the selective autophagy pathway in the vacuole. **(B) XopL blocks autophagy**: XopL interacts with autophagy component SH3P2 inside the cell and ubiquitinates it to degrade it via the 26S proteasome. Degradation of SH3P2 results in defects of autophagosome delivery into the vacuole and hence suppresses autophagy.

### Autophagy in host-immune interactions

In animals, several pathogenic bacteria have been identified to modulate the autophagy pathway to their own benefit (Huan & Brummel, 2014). Many intracellular animal pathogenic bacteria can be eliminated by autophagy, while others (such as *Shigella*, *Yersinia* and *Listeria*), are able to exploit autophagy to increase their pathogenicity (Huan & Brummel, 2014). In plants, several studies have highlighted that pathogens manipulate the autophagy pathway (Leary *et al*, 2018). More specifically, the plant pathogenic bacterium *Pseudomonas syringae* pv. *tomato* (*Pst*) has been shown to utilize effectors to modulate autophagic degradation (Üstün *et al*, 2018; Üstün *et al*, 2016). With *Xcv,* we have identified another plant pathogenic bacterium that modulates autophagy, similar to *Pst*, in an effector-dependent manner. Although, both pathogens have the same habitat and hemi-biotrophic lifestyle, they act in different ways on the autophagy pathway. For *Xcv* inhibition of the autophagy pathway is crucial to maintain pathogenicity while *Pst* activates it for its own benefit (Üstün *et al*, 2018). Previously, it has been reasoned that autophagy activation might be essential to maintain plant viability and lifespan (Hafrén *et al*, 2018). However, autophagy may not be required for this during *Xcv* infection, as it has been shown that T3Es XopD and XopJ are able to prolong the biotrophic phase by other mechanisms (Üstün et al, 2013; (Kim *et al*, 2008). As such, autophagy is dispensable for the virulence of *Xcv* and actively dampened to boost virulence and partially prevent xenophagy of XopL. Within the realm of xenophagy, the degradation of effectors by autophagy can be considered as a form of “effectorphagy”. Perturbation of general autophagy is achieved by degrading SH3P2 through the action of T3E XopL. SH3P2 was first identified as a novel autophagy component, stimulating autophagosome formation during nitrogen starvation and BTH-triggered immune responses (Zhuang *et al*, 2013). Later studies also showed that SH3P2 plays a key role in membrane tubulation during cell plate formation (Ahn *et al*, 2017) and clathrin-mediated endosomal sorting and degradation, with no effect in dark-induced autophagy (Nagel *et al*, 2017). Our findings that silencing of SH3P2 from *N. benthamiana* impairs autophagy and somewhat promotes pathogenicity sheds further light on its diverse functions. However, the effects of XopL on SH3P2 and increasing virulence of *Xcv* might not only be attributed to its function in autophagy. Because endocytic trafficking also plays a major role in plant immunity (Gu *et al*, 2017), it is likely that XopL has a function beyond autophagy to impair plant defense mechanisms. This is also supported by the fact that XopL can reduce PTI responses (Singer *et al*, 2013), which is not due its function in autophagy, as autophagy deficiency has no impact on PTI responses (Lenz *et al*, 2011). Previously, XopL was also shown to impact stromule formation by localizing to microtubules (Erickson *et al*, 2018), which was shown only for the XopL version lacking E3 ligase activity. Although, we do not see any microtubule localization of the XopL ΔE3 version, XopL might affect stromule formation as they are thought to be recognized by autophagic membranes prior autophagic degradation (Marshall & Vierstra, 2018).

### Role of xenophagy in immunity

One of the best studied selective autophagy receptors across kingdoms is NBR1/p62 that plays a central role in xenophagy and degradation of protein aggregates (Kirkin *et al*, 2009; Svenning *et al*., 2011). Its role in plant immune responses has been shown by the involvement of NBR1/Joka2-mediated selective autophagy in restricting pathogen growth or disease progression during *P. infestans* or *Pst* infection (Üstün *et al*, 2018, Dagdas *et al*, 2016). In animals, p62, the counterpart of plant NBR1, functions to mediate xenophagy, which has been also described for NBR1 in plants in a plant-virus infection context but not for other plant pathogens (Hafrén et al, 2017, Hafrén *et al*, 2018). Our analysis provides the first evidence that like viral proteins, bacterial effector XopL constitutes a target of NBR1-mediated selective autophagy. This sheds light on previous findings about the role of NBR1/Joka2 in plant immunity and makes it a central hub in plant-microbe interactions. Consequently, XopL developed the ability to suppress autophagy, to boost the virulence of Xcv. But does XopL suppress autophagy to an extent to escape its own degradation? Indeed, treating plants with ConA or expressing AIMp still stabilize XopL protein levels, indicating that the effect of XopL on the autophagy pathway is either very specific or not sufficient to shut down the pathway completely. It is also possible that XopL is degraded by other additional mechanisms such as endocytosis or proteasome-mediated degradation as it most likely ubiquitinated by plant E3 ligases. In line with our observations on the autophagy pathway is that the ability of XopL to suppress the autophagy is still less than the recently discovered autophagy inhibitor AIMp (Pandey *et al*, 2021). This is also in line with the fact that loss of SH3P2 is only partially suppressing autophagy formation (Zhuang *et al*, 2013) while silencing of ATG7 or expression of AIMp result in a complete block of autophagy. Currently, we also do not know whether SH3P2 might only affect a subset of ATG8 isoforms to facilitate autophagosome formation. Thus, it might be also possible that the NBR1/Joka2-selective autophagy pathway might involve ATG8 isoforms that do not require SH3P2.

To date, it has not been reported that bacterial effectors in animals and plants are removed by selective autophagy as an anti-microbial response. Interestingly, the *Salmonella* effector SseL, which inhibits selective autophagy to abolish p62-dependent degradation of *Salmonella*, was also found to interact with p62 (Mesquita *et al*, 2012). This might suggest the possibility that SseL might also have been an autophagy target before it acquired its function to block this pathway via its deubiquitinase activity (Mesquita *et al*, 2012). We hypothesize that NBR1/p62 might have evolved to have a function in anti-bacterial autophagy by triggering xenophagy of bacterial molecules as an alternative strategy to degrade entire intracellular bacteria. This may have happened for the function of NBR1 in plants, as fungal and oomycete pathogens as well as Gram-negative bacteria reside in the extracellular space. Animal pathogens also occupy the extracellular space, before entering the host cell. In the case of *Salmonella*, it first needs to inject bacterial effectors via its SPI-1 T3SS to establish internalization and its replication niche (Lou *et al*, 2019). It is therefore tempting to speculate that these effectors may be targeted by selective autophagy mechanisms as an early defense mechanism of the immune system. Similar to XopL, several of the SPI1 T3Es can mimic E3 ligases and/or are ubiquitinated in the host cell (Kubori & Galan, 2003), making them potential targets for NBR1/p62-mediated selective autophagy. Indeed, T3Es SopA and SopE have been reported to be degraded through the proteasome (Kubori & Galan, 2003; Zhang *et al*, 2005). A possible degradation by autophagy was not investigated in these studies. Our results also suggest that XopL is targeted by a host E3 ligase for degradation as the XopL variant lacking E3 ligase activity is still ubiquitinated *in planta* and degraded by autophagy. Several E3 ligases have been implicated in plant-microbe interactions, which opens up the possibility that they may target microbial proteins (Furlan *et al*, 2012).

Although plant pathogenic bacteria possess T3Es that are implicated in the host ubiquitin system, to date there is no evidence that they might be ubiquitinated in the host. In addition, we identified that XopL undergoes self-ubiquitination, which has not been reported for animal and plant pathogenic bacterial effectors.

In this scenario, it is tempting to speculate whether the self-ubiquitination activity of XopL attracts it to the autophagy pathway. Although the biological significance of K191 remains elusive, it might still have regulatory functions. To date, self-ubiquitination of E3 ligases has been assigned as a mechanism of self-regulation through which their activity is controlled (Furlan *et al*., 2017). In the case of bacterial T3Es that mimic E3 ligases, it might be a strategy to trick degradation systems. Other post-translational modifications of T3Es such as phosphorylation of AvrPtoB have been found to crucial for its virulence function (Lei *et al*, 2020), which might be also the case for the ubiquitination of XopL. Our results suggest that ubiquitinated XopL is targeted for autophagic degradation which might be indeed a strategy to recruit different autophagy components. Indeed, we can find a high conservation of K191 in several other XopL-like T3Es across different Xanthomonas species (Fig S18), which would be in favor of the proposed hijacking hypothesis. However, the fact that the K191 variant is still ubiquitinated *in planta* and associates with NBR1 and other autophagy components are not in favor of this hypothesis. The discovery of other alternative ubiquitination sites in XopL will help us to unravel why XopL undergoes self-ubiquitination and how this might contribute to its virulence.

Taken together, we provide a primary example where a bacterial effector subverts host autophagy to downregulate host immunity, and the autophagic machinery in turn targets the same bacterial effector for degradation. Thus, this reveals a complex multi-layered role of autophagy particularly in the context of immunity and disease. Additionally, XopL possesses self-ubiquitination activity, which some evidence we have uncovered here suggest that this functions to hijack the host autophagy system.

## Acknowledgements

We thank Ingrid Schießl for the initial Y2H screen with XopL. We also thank Tom Denyer for critical reading of the manuscript. We thank Johannes Stuttmann for the *roq1 N. benthamiana* seeds. We thank Silke Wahl and Irina Droste-Borel for technical support for the proteomics assay. This work was supported by an Emmy Noether Fellowship GZ: UE188/2-1 from the Deutsche Forschungsgemeinschaft (DFG; to S.Ü.), and through the collaborative research council 1101 (SFB1101; to G.L.). E.A.M. was supported by EU Horizon 2020 MSCA IF (799433). F.B. was supported by funds from the DFG (BO1961/5-2). D.H. was supported by grants from the Swedish research councils VR (2016-04562; 2020-05327) and FORMAS (2017-01596). T.O.B. was supported by BBSRC - Biotechnology and Biological Sciences Research Council, grant code BB/T006102/1. M.B.M. was supported by NSF IOS Grant 2026368. FORMAS (grant number 2016-01044) for A.H. We thank the confocal microscopy facility of ZMBP that is supported by funds from DFG (INST 37/819-1 FUGG and INST 37/965-1 FUGG), especially Sandra Richter and Natalie Krieger for their introduction into the Zeiss LSM880, Leica SP8 and FRET-FLIM.

## Material and Method

### Plant Material and Growth Conditions

Wild-type plants were *Arabidopsis thaliana* ecotype Columbia (Col-0) and *Nicotiana benthamiana. Arabidopsis* plants were grown on soil under short-day conditions (8/16-h light/dark cycles) in a growth chamber or for maintenance and crossings under long-day conditions (16/8-h light/ dark cycles) in a growth room with light intensity of 150 μE, 21°C, and 70% relative humidity, respectively. *N. benthamiana* plants were grown under long day conditions 16/8-h light/dark cycles, 21°C, and 70% relative humidity.

### Plasmid construction

For transient expression experiments, the coding region of XopL, AtSH3P2 or SlJoka2 were cloned into pENTR/D-TOPO and subsequently recombined into pUBN-DEST-GFP or RFP (Grefen *et al*, 2010), pGWB614/5 (Nakagawa *et al*, 2007), pMAlc2, pDEST15. The RFP-ATG8E/G, GFP-ATG8e, RFP-NBR1, RLUC-ATG8, RLUC-NBR1, XopD-GFP, XopJ-GFP, pTRV2-Joka2, pTRV2-ATG7 constructs were described previously (Üstün *et al,* 2018, Üstün *et al*, 2013). All binary plasmids were transformed into Agrobacterium tumefaciens strain C58C1 and infiltration of *N. benthamiana* was done at the four-to six-leaf stage. Stable Arabidopsis transformation was performed using the floral dip method (Clough & Bent, 1998).

### Transient Expression in *N. benthamiana* by Agrobacterium-Mediated Leaf Infiltration

Transient expression was performed as described previously (Üstün *et al*, 2018).

### Immunoprecipitation

GFP pull-down assays were performed as previously described (Üstün *et al*, 2018. Pulldown of ubiquitinated proteins was performed according to manufacturer’s instructions (NanoTag Biotechnologies).

### *In vitro* pull-down

In vitro pull-down was performed as previously described (Üstün *et al*, 2013).

### Dual Luciferase Assay

Dual luciferase assay was performed as described previously (Üstün *et al*, 2018).

### Virus-induced gene silencing

VIGS was performed as described previously (Üstün *et al*, 2013).

### Bacterial growth conditions

*Agrobacterium tumefaciens*, Agrobacteria strain C58C1 was grown in LB Hi-Salt (10g/L sodium chloride, 10g/L tryptone, 5g/L yeast extract) with 100 μg mL−1 rifampicin at 28 °C. The cultures supplemented with the appropriate antibiotics for those harboring plasmids. Xcv strain 85-10 was grown in NYG media (0.5% peptone, 0.3% yeast extract, 2% glycerol) with 100 μg mL−1 rifampicin at 28 °C.

### Construction of Xcv *ΔxopL* and *ΔxopQ* null mutants

To construct Xcv 85-10 *xopL* and *xopQ* deletion mutants, the 1.7-kb upstream and downstream regions of the xopL or xopQ gene were PCR amplified using Xcv 85-10 genomic DNA as template and cloned into pLVC18 linearized with EcoRI (New England Biolabs) using Gibson assembly. The plasmid was introduced into Xcv 85-10 by triparental mating. Xcv transconjugants were analysed by PCR to confirm that homologous recombination occurred at the xopL or XopQ locus.

### Constructs for Xcv *ΔxopL* complementation analysis

To construct xopL gene with 0.3-kb promoter region in broad host range vector, 0.3-kb promoter-xopL gene was PCR amplified using Xcv 85-10 genomic DNA as template and cloned into pBBR1MCS-2 (Kovach *et al*, 1995) linearized with EcoRV (New England Biolabs) using Gibson assembly. The plasmid was introduced into Xcv 85-10 Δ*xopL* by triparental mating.

### Bacterial infection

Xcv carrying a deletion mutation of XopQ (Xcv *ΔxopQ*), or of HrcN (Xcv Δ*hrcN*), or of both XopQ and XopL (Xcv *ΔxopQ ΔxopL*) were used to infect wild-type *N. benthamiana*. Xcv were grown overnight in NYG with appropriate antibiotics at 28°C with shaking. Bacteria were diluted to OD_600_ = 0.2 for dual-luciferase assays, immunoblot analysis of NBR1, ATG8 or confocal microscopy of autophagosomal structures. For in planta growth curves using syringe infiltration, Xcv strains were inoculated at OD_600_ = 0.0004, and for dip-inoculation OD_600_ = 0.2 was used.

### Tomato growth condition and bacterial growth assay

Tomato (*Solanum lycopersicum*) cv. VF36 was grown in greenhouse (22-28 °C, 50-70 % RH, 16-h light). For bacterial growth assays, leaflets were dipped in a 2 x 10^8^ CFU/mL suspension of Xcv 85-10 strains in 10mM MgCl2 with 0.025% (v/v) silwet L-77 (Helena Chemical Company) for 30 sec. Plants were then placed in plastic chambers at high humidity (>95%) for 24 h. For each strain analyzed, four leaf discs (0.5 cm^2^) per treatment per timepoint were ground in 10 mM MgCl_2_ and diluted and spotted onto NYGA plates in triplicate to determine bacterial load. Three biological replicates (i.e., three plants) were used, and the experiment was repeated at least three times.

### Confocal Microscopy

Live-cell images were acquired from abaxial leaf epidermal cells using a Zeiss LSM780 and LSM880 microscope. Excitation/emission parameters for GFP and RFP were 488 nm/490 to 552 nm and 561 nm/569 to 652 nm, respectively, and sequential scanning mode was used for colocalization of both fluorophores. Confocal images with ImageJ (version 2.00) software. Quantification of ATG8-labeled autophagosomal structures was done on z-stacks that were converted to eight-bit grayscale and then counted for ATG8 puncta either manually or by the Particle Analyzer function of ImageJ.

### FRET-FLIM measurement

FRET-FLIM was performed on SP8 confocal laser scanning microscope (CLSM) (Leica Microsystems GMBH) with LAS AF and SymPhoTime 64 software using a 63x/1.20 water immersion objective. FLIM measurements were performed with a 470 nm pulsed laser (LDH-P-C-470) with 40 MHz repetition rate and a reduced speed yielding, with an image resolution of 256×256, a pixel dwell time of ∼10 µs. Max count rate was set to ∼15000 cps. Measurements were stopped, when the brightest pixel had a photon count of 1000. The corresponding emission was detected with a Leica HyD SMD detector from 500 nm to 530 nm by time-correlated single-photon counting using a Timeharp260 module (PicoQuant, Berlin). The calculation of GFP lifetime was performed by iterative reconvolution, i.e. the instrument response function was convolved with exponential test functions to minimize the error with regard to the original TCSPC histograms in an iterative process. For measurements of GFP-JOKA2 protein aggregates, ROIs were drawn manually on SymphoTime64 software around the aggregates to analyze GFP lifetime in these structures.

### Immunoblot analysis

Proteins were extracted in 100 mM Tris (pH 7.5) containing 2% SDS, boiled for 10min in SDS loading buffer, and cleared by centrifugation. The protein extracts were then separated by SDS-PAGE, transferred to PVDF membranes (Biorad), blocked with5% skimmed milk in PBS, and incubated with primary antibodies anti-NBR1 (Agrisera), anti-ATG8 (Agrisera), anti-ubiquitin (Agrisera), anti-GFP (SantaCruz), anti-RFP (Chromotek), anti-HA (Sigma Aldrich) primary antibodies using 1:2000 dilutions in PBS containing 0.1% Tween 20. This was followed by incubation with horseradish peroxidase-conjugated secondary antibodies diluted 1:10,000 in PBS containing 0.1% Tween 20. The immunoreaction was developed using an ECL Prime Kit (GE Healthcare) and detected with Amersham Imager 680 blot and gel imager.

### *In vitro* ubiquitination assay

Recombinant proteins were expressed in *E. coli* BL21(DE3) and purified by affinity chromatography using amylose resin (New England Biolabs). Recombinant His-UBA1 and His-UBC8 were purified using Ni-Ted resin (Macherey-Nagel). Purified proteins were used for *in vitro* ubiquitination assays. Each reaction of 30 mL final volume contained 25 mM Tris-HCl, pH 7.5, 5 mM MgCl2, 50 mM KCl, 2 mM ATP, 0.6 mM DTT, 2 μg ubiquitin, 200 ng E1 His-*At*UBA1, 1.2 μg E2 His-*At*UBC8, 2 μg of E3s, and 0.3 µg of MBP-*At*SH3P2. Samples were incubated for 1h at 30°C and reaction was stopped by adding SDS loading buffer and incubated for 10 min at 68°C. Samples were separated by SDS-PAGE electrophoresis using 4–15% Mini-PROTEAN® TGX™ Precast Protein Gels (BioRad) followed by detection of ubiquitinated substrate by immunoblotting using anti-MBP (New England Biolabs), anti-GST and anti-ubiquitin (Santa Cruz Biotechnology) antibodies.

### NanoLC-MS/MS analysis and data processing

Proteins were purified on an NuPAGE 12% gel (Invitrogen) and Coomassie-stained gel pieces were digested in gel with trypsin as described previously (Borchert *et al*, 2010) with a small modification: chloroacetamide was used instead of iodoacetamide for carbamidomethylation of cysteine residues to prevent formation of lysine modifications isobaric to two glycine residues left on ubiquitinylated lysine after tryptic digestion. After desalting using C18 Stage tips peptide mixtures were run on an Easy-nLC 1200 system coupled to a Q Exactive HF-X mass spectrometer (both Thermo Fisher Scientific) as described elsewhere (Kliza *et al*, 2017) with slight modifications: the peptide mixtures were separated using a 87 minute segmented gradient from 10-33-50-90% of HPLC solvent B (80% acetonitrile in 0.1% formic acid) in HPLC solvent A (0.1% formic acid) at a flow rate of 200 nl/min. The seven most intense precursor ions were sequentially fragmented in each scan cycle using higher energy collisional dissociation (HCD) fragmentation. In all measurements, sequenced precursor masses were excluded from further selection for 30 s. The target values were 105 charges for MS/MS fragmentation and 3×106 charges for the MS scan.

Acquired MS spectra were processed with MaxQuant software package version 1.5.2.8 with integrated Andromeda search engine. Database search was performed against a *Nicotiana benthamiana* database containing 74,802 protein entries, the sequences of XopL from *Xanthomonas campestris* pv. *vesicatoria*, and 285 commonly observed contaminants. Endoprotease trypsin was defined as protease with a maximum of two missed cleavages. Oxidation of methionine, phosphorylation of serine, threonine and tyrosine, GlyGly dipetide on lysine residues, and N-terminal acetylation were specified as variable modifications. Carbamidomethylation on cysteine was set as fixed modification. Initial maximum allowed mass tolerance was set to 4.5 parts per million (ppm) for precursor ions and 20 ppm for fragment ions. Peptide, protein and modification site identifications were reported at a false discovery rate (FDR) of 0.01, estimated by the target-decoy approach (Elias and Gygi). The iBAQ (Intensity Based Absolute Quantification) and LFQ (Label-Free Quantification) algorithms were enabled, as was the “match between runs” option (Schwanhausser *et al*, 2011).

### RNA extraction and RT-qPCR

RNA was extracted from 4 leaf discs according to manufacturer instructions using the GeneMATRIX Universal RNA Purification Kit (Roboklon) with on-column DNase I digestion. RNA integrity was checked by loading on 1% agarose gel and separating by electrophoresis. RNA concentrations were measured using Nanodrop 2000 (Thermo Fisher), and equal amounts of RNA were used for cDNA synthesis. cDNA synthesis was performed using LunaScript™ RT SuperMix Kit (New England Biolabs) and in a standard thermocycler according to manufacturer instructions. Gene expression was measured by qPCR using MESA BLUE qPCR MasterMix Plus for SYBR® Assay No ROX (Eurogentec) and cycle quantification by Biorad CFX system.

### Drug treatments

For the analysis of protein stability 200 μM MG132 or 1% EtOH was infiltrated to plants transiently expressing binary constructs 2dpi. 6 hours later leaf material was harvested. were analysed under the CLSM. Concanamycin A treatment was performed by syringe infiltration of mature leaves with 0.5 μM ConA for 6-8 h prior confocal analysis or immunoblot analysis. AZD8055 (15 μM) was done for 6-8 hours prior confocal microscopy.

### Phylogenetic Analysis

An alignment between SH3P2 proteins was generated using ClustalW wand the tree was generated using the neighbor-joining method. Effector proteins related to XopL were identified performing a BLASTp (https://blast.ncbi.nlm.nih.gov/Blast.cgi) using XopL protein sequence (ID: CAJ24951.1). 18 protein sequences were extracted from the Top100 sequences from Blast results with only the top hit considered per species/pathovars. Related effectors from more distant bacteria were identified realizing a second BLASTp excluding Xanthomonas genus and 3 proteins from relevant species were extracted from the Top10 results. Protein from X*anthomonas campestris pv. Campestris* was included manually as an example of non-conserved protein from Xanthomonas genus. Multiple Sequence Alignment of the 24 extracted sequences was performed using COBALT (https://www.ncbi.nlm.nih.gov/tools/cobalt/cobalt.cgi) from NCBI.

### Data Analysis and Presentation

Data are presented as boxplots with visible datapoints, where middle horizontal bars of boxplots represent the median, the bottom and top represent the 25th and 75th percentiles, whiskers extend to at most 1.5 times the interquartile range. Statistical significance was analysed using appropriate statistical tests, either by Student’s t test, one way ANOVA or Kruskal-Wallis rank sum test (*P < 0.05, **P < 0.01, and P*** < 0.001). The number of biological replicates (n) is given in the figure legends. Statistical analyses and graphical presentation of data were made in R and RStudio (Version 1.2.5033). Boxplots were prepared using the ggplot2 package.

## Author contributions

J.X.L, S.Ü., M.R., D.S., G.L., A.R.G., J.-G.K., M.F.-W. performed the experiments. J.X.L-, M.R., G.L., E.A.M., M.F.W., B.M., A.H., T.O.B., M.B.M., F.B., D.H., and S.Ü. analysed the data. P.P. and T.O.B. provided novel material. S.Ü. planned the project and wrote the article together with J.X.L and input from all authors.

## Supporting information

**Fig. S1:**
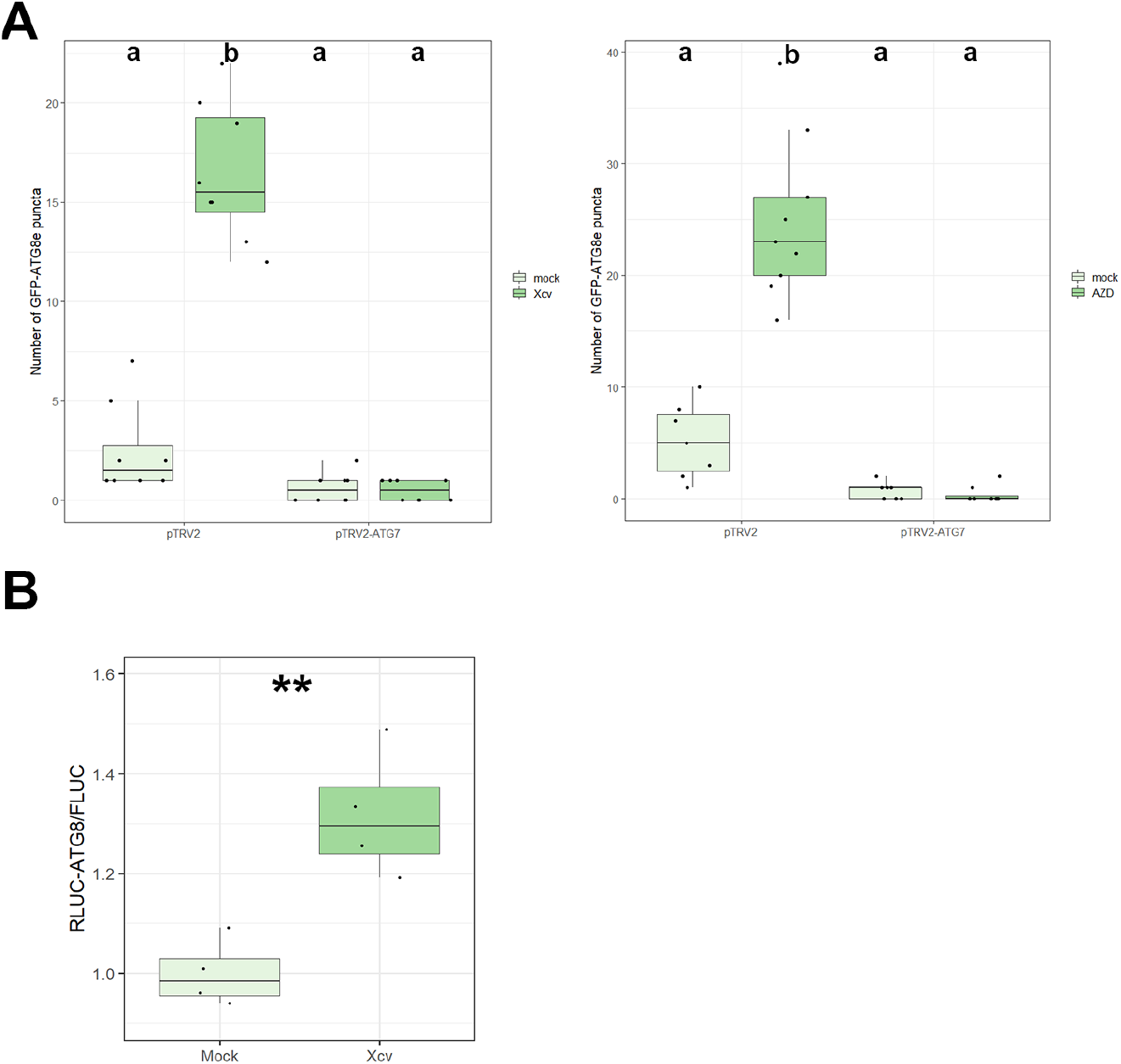
Silencing of *ATG7* in *N. benthamiana* plants abolishes autophagosome formation and Xcv blocks autophagy at 6hpi. GFP-ATG8e-labeled puncta were quantified from plants silenced for *ATG7* (pTRV2-ATG7) infected with mock or Xcv *ΔxopQ* at 6hpi in the presence or absence of ConA, and of AZD. Puncta were calculated from z-stacks (X) of *n*=12 individuals using ImageJ. Different letters indicate statistically significant different groups (*P* < 0.05) as determined by one way ANOVA. **(B)** RLUC-ATG8a or RLUC-NBR1 constructs were coexpressed with internal control FLUC in *N. benthamiana*. *Xcv ΔxopQ* or infiltration buffer (mock) was co-infiltrated with Agrobacteria containing the respective constructs. RLUC and FLUC activities were simultaneously measured in leaf extracts at 8 h post-infiltration using the dual-luciferase system. Values represent the ratio of RLUC-ATG8a and FLUC activities (n=4). Statistical significance (\*\*\**P* < 0.001) was revealed by Student’s *t*-test. The experiment was repeated 3 times with similar results.

**Fig. S2:**
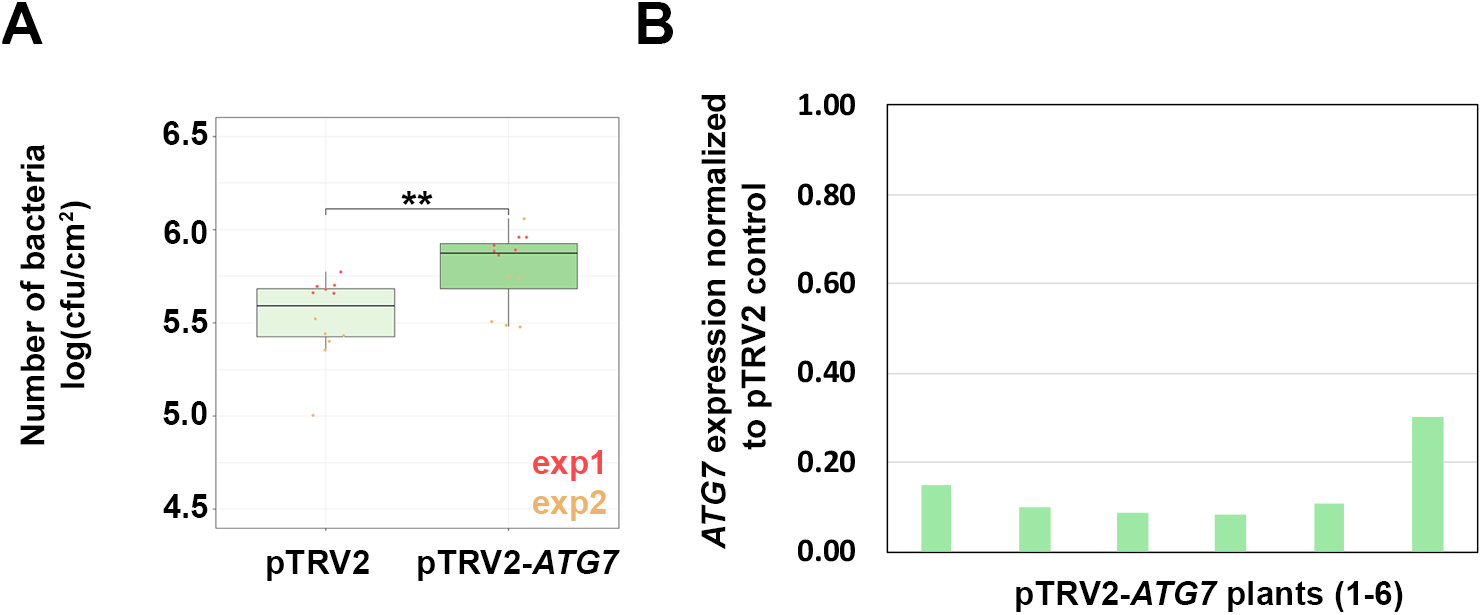
Virus induced gene silencing of *ATG7* in *N. benthamiana* is beneficial for Xcv. **(A)** Growth of Xcv *ΔxopQ* in *N. benthamiana* plants silenced for *ATG7* (pTRV2-*ATG7*) compared to control plants (pTRV2). Leaves were dip-inoculated with a bacteria suspension at OD_600_ = 0.2 and bacteria were quantified at 6 dpi. Data represent the mean SD (n = 6). Significant differences were calculated using Student’s *t-test* and are indicated by **, P < 0.01. The experiment was repeated twice with similar trends. Red and yellow data points represent repeats of the experiment. **(B)** qRT-PCR analysis of *ATG7* mRNA levels in silenced *N. benthamiana* plants. *Actin* expression was used to normalize the expression value in each sample, and relative expression values were determined against the mean expression in pTRV2 (control) plants.

**Fig. S3:**
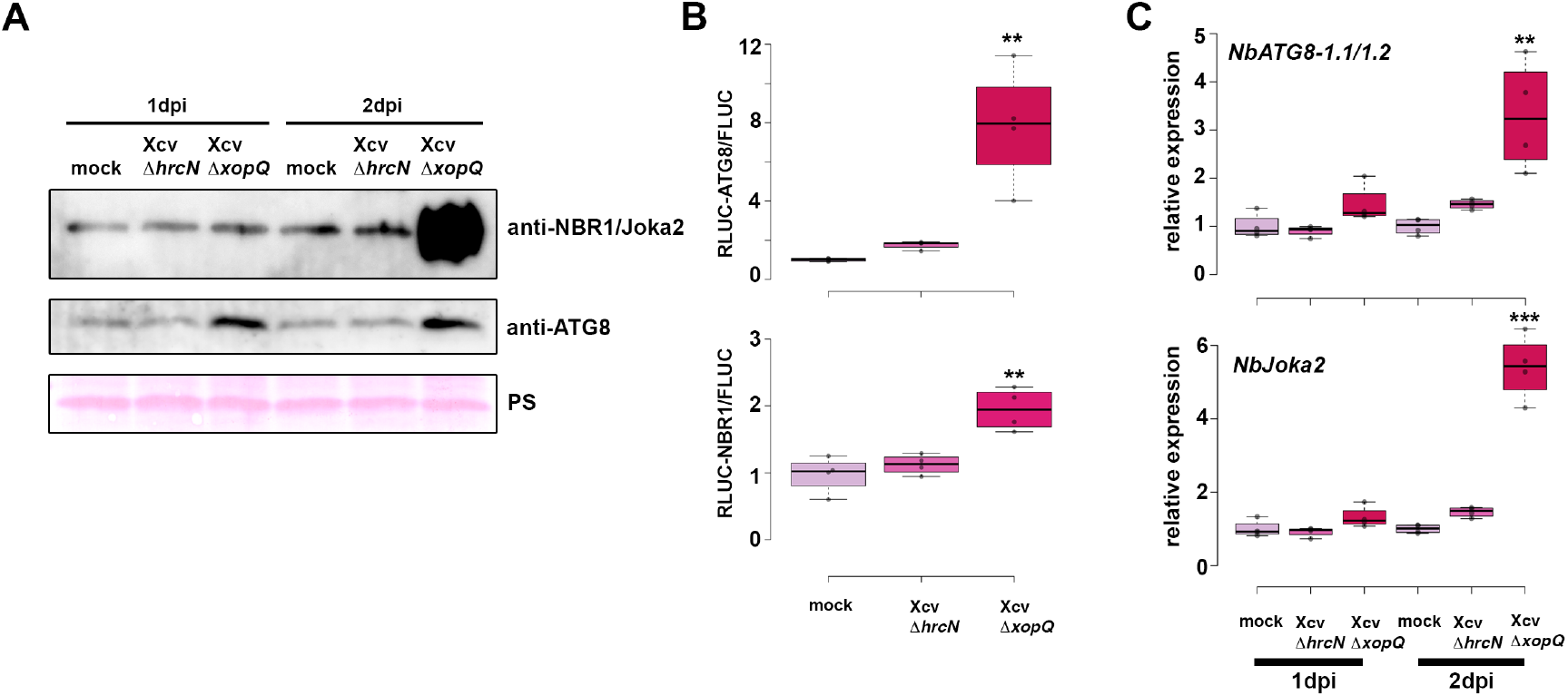
Suppression of autophagy is enhanced by T3Es. **(A)** Immunoblot analysis of NBR1 and ATG8 protein levels in Xcv *ΔxopQ, ΔhrcN* or mock infected *N. benthamiana* plants at 1 and 2dpi. Ponceau Staining (PS) served as a loading control. The experiment was repeated twice with similar results. **(B)** Autophagic flux determined by quantitative dual-luciferase assay. RLUC-ATG8a or RLUC-NBR1 constructs were coexpressed with internal control FLUC in *N. benthamiana*. Xcv *ΔxopQ* and *ΔhrcN* were respectively co-infiltrated with Agrobacteria containing the luciferase constructs. *Renilla* and *Firefly* luciferase activities were simultaneously measured in leaf extracts at 48 h post-infiltration using the dual-luciferase system. Values represent the ratio of RLUC-ATG8a and FLUC activities normalized to mock (n=4). Statistical significance (\*\**P*<0.01) was revealed by Student’s *t*-test. The experiment was repeated 2 times with similar results. **(C)** RT-qPCR analysis of *NbATG8-1.1/2* and *NbJoka2* transcript levels upon challenge of *N. benthamiana* plants with Xcv *ΔxopQ* and *ΔhrcN* for 1 and 2 dpi compared to mock infected plants. Values represent expression relative to mock control of respective time point and were normalized to *actin*. Statistical significance (*** *P* < 0.001) was revealed by Student’s *t*-test.

**Fig. S4:**
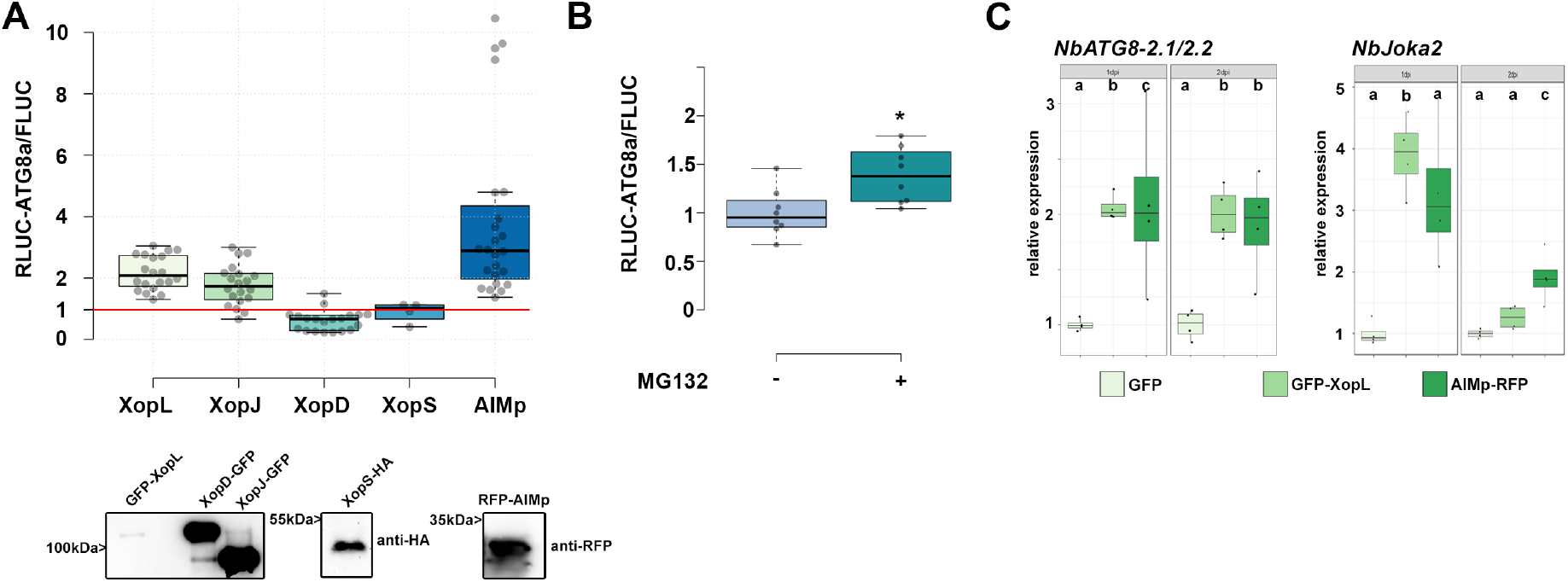
Screening for Xanthomonas T3Es with altered autophagic flux. **(A)** RLUC-ATG8a constructs were coexpressed with internal control FLUC in *N. benthamiana*. GFP-XopL, XopJ-GFP, XopD-GFP and XopS-HA were co-infiltrated with Agrobacteria carrying the RLUC-ATG8a and FLUC constructs. Renilla and Firefly luciferase activities were simultaneously measured in leaf extracts at 48 h post-infiltration using the dual-luciferase system. Values represent the ratio of RLUC-ATG8a to FLUC activity normalized to GFP control (XopL, XopJ, XopD, AIMp; n=20; XopS n=4). Expression of T3Es and RFP-AIMp were verified with the indicated antibodies. **(B)** RLUC-ATG8a constructs were coexpressed with internal control FLUC in *N. benthamiana*. Plants were treated with MG132 for 6 hours prior measurement. Values represent the ratio of RLUC-ATG8a to FLUC activity normalized to vector control (n=8). Statistical significance (* *P* < 0.5) was revealed by Student’s *t*-test. **(C)** RT-qPCR analysis of *NbATG8-2.1/2* and *NbJoka2* transcript levels upon Agrobacteria-mediated transient expression of GFP, GFP-XopL or AIMp for 1 and 2 dpi. Values represent expression relative to GFP control of respective time point and were normalized to *actin*. Statistical significantly different groups are denoted by different letters, as calculated using Kruskal-Wallis rank sum test (*P*<0.05).

**Fig. S5:**
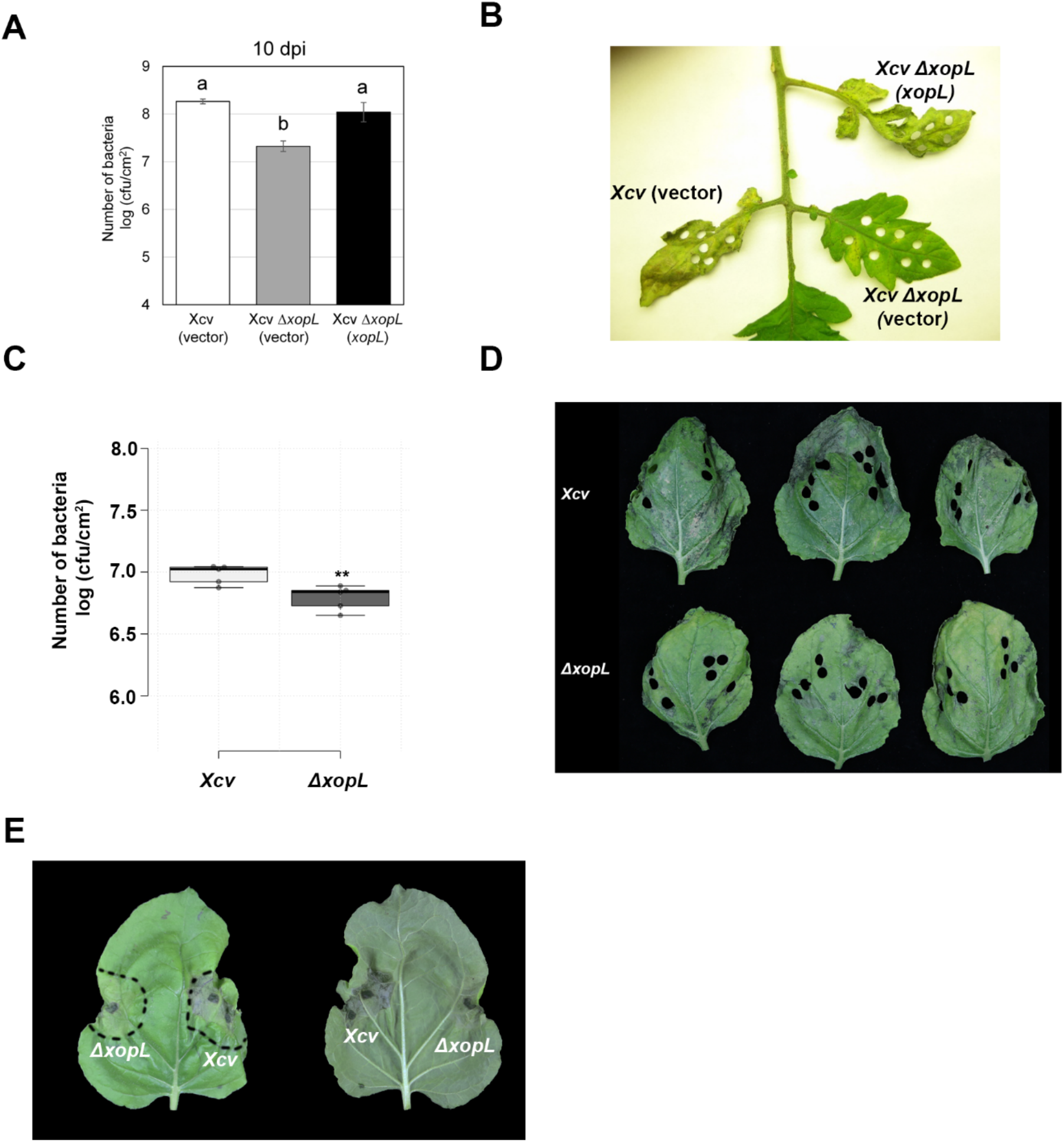
XopL contributes to Xcv virulence. **(A)** Growth of Xcv 85-10 (vector) (white bar), Xcv 85-10 Δ*xopL* (vector) (grey bar), and Xcv 85-10 Δ*xopL* (*xopL*) (black bar) strains in tomato VF36 leaves. Leaves were dipped in a 2 x 10^8^ CFU/mL suspension of bacteria. The number of bacteria in each leaf was quantified at 10 dpi. Data points represent mean log_10_ colony-forming units per cm^2^ ± SD of three plants. Different letters above bars indicate statistically significant (Tukey’s honestly significant difference (HSD) test, P < 0.05) differences between samples. Vector = pBBR1MCS-2. **(B)** Delayed disease symptom development in tomato leaves inoculated with *Xcv* or *Xcv* Δ*xopL*. Tomato leaves inoculated with strains described in (A) were photographed at 14 dpi. **(C)** Growth of Xcv 85-10, Xcv 85-10 Δ*xopL* strains in *roq1 N. benthamiana* leaves. Leaves were dipped in a 2 x 10^8^ CFU/mL suspension of bacteria. The number of bacteria in each leaf was quantified at 10 dpi (n = 5). Significant differences were calculated using Student’s *t-test* (**, P < 0.01). The experiment was repeated twice with similar trends. **(D)** Delayed disease symptom development in *roq1 N. benthamiana* leaves dip-inoculated with *Xcv* or *Xcv* Δ*xopL*. *N. benthamiana roq1* leaves inoculated with strains described in (C) were photographed at 10 dpi. **(E)** Delayed symptom development in *roq1 N. benthamiana* leaves inoculated with *Xcv* or *Xcv* Δ*xopL*. Leaves were syringe-inoculated with OD_600_=0.2 and photographed at 3 dpi.

**Fig. S6:**
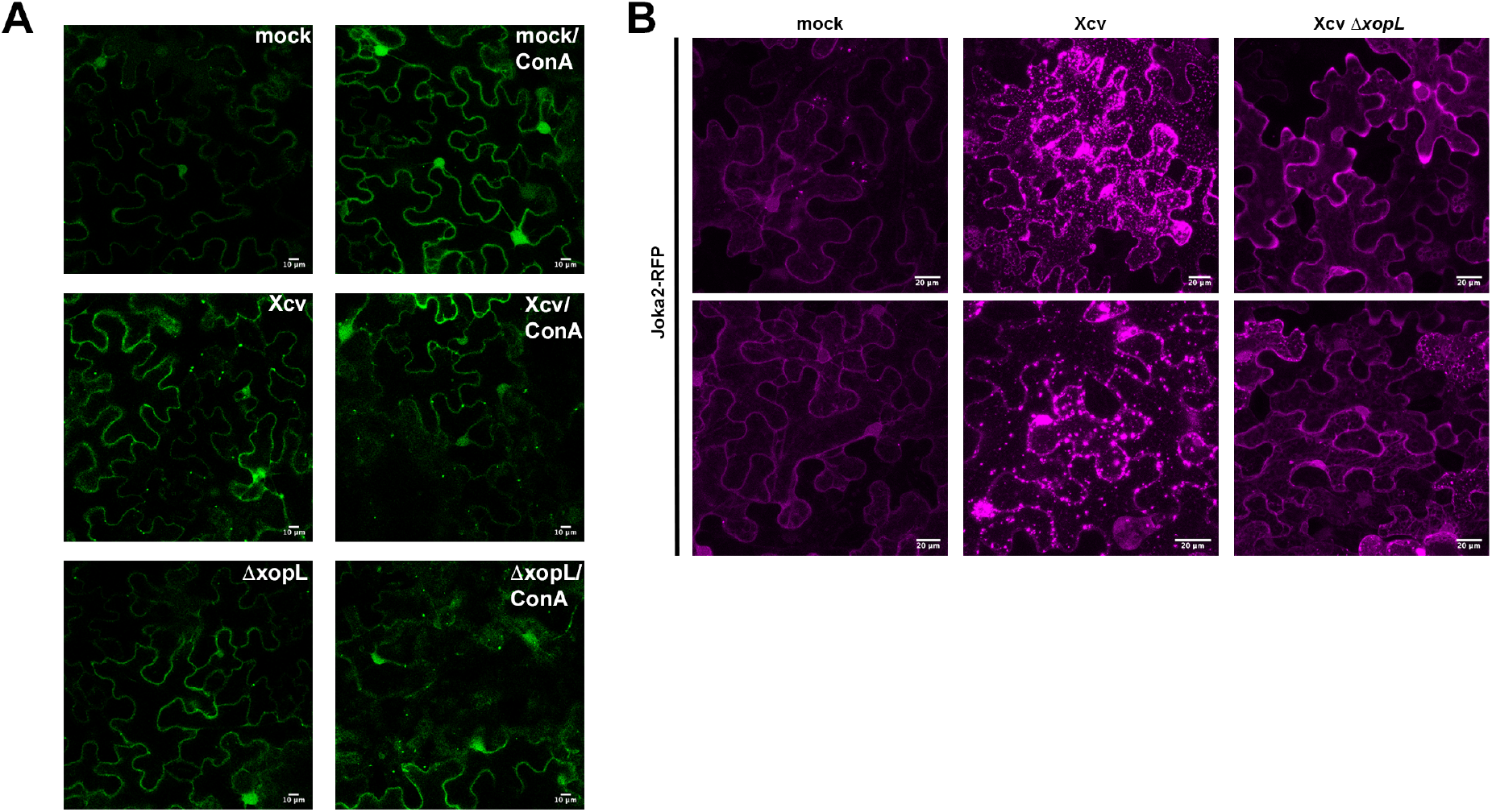
Joka2 bodies are induced during Xcv infection in a XopL-dependent manner. **(A)** GFP-ATG8e-labeled autophagosomes imaged from *N. benthamiana* plants infected with mock, Xcv or *Xcv* Δ*xopL* at 2 dpi in the presence or absence of ConA (bars = 10 μm). **(B)** RFP-Joka2 labelled puncta or aggregates upon challenge of *N. benthamiana* leaves with mock, *Xcv* or *Xcv* Δ*xopL* infection at 1dpi.

**Fig. S7:**
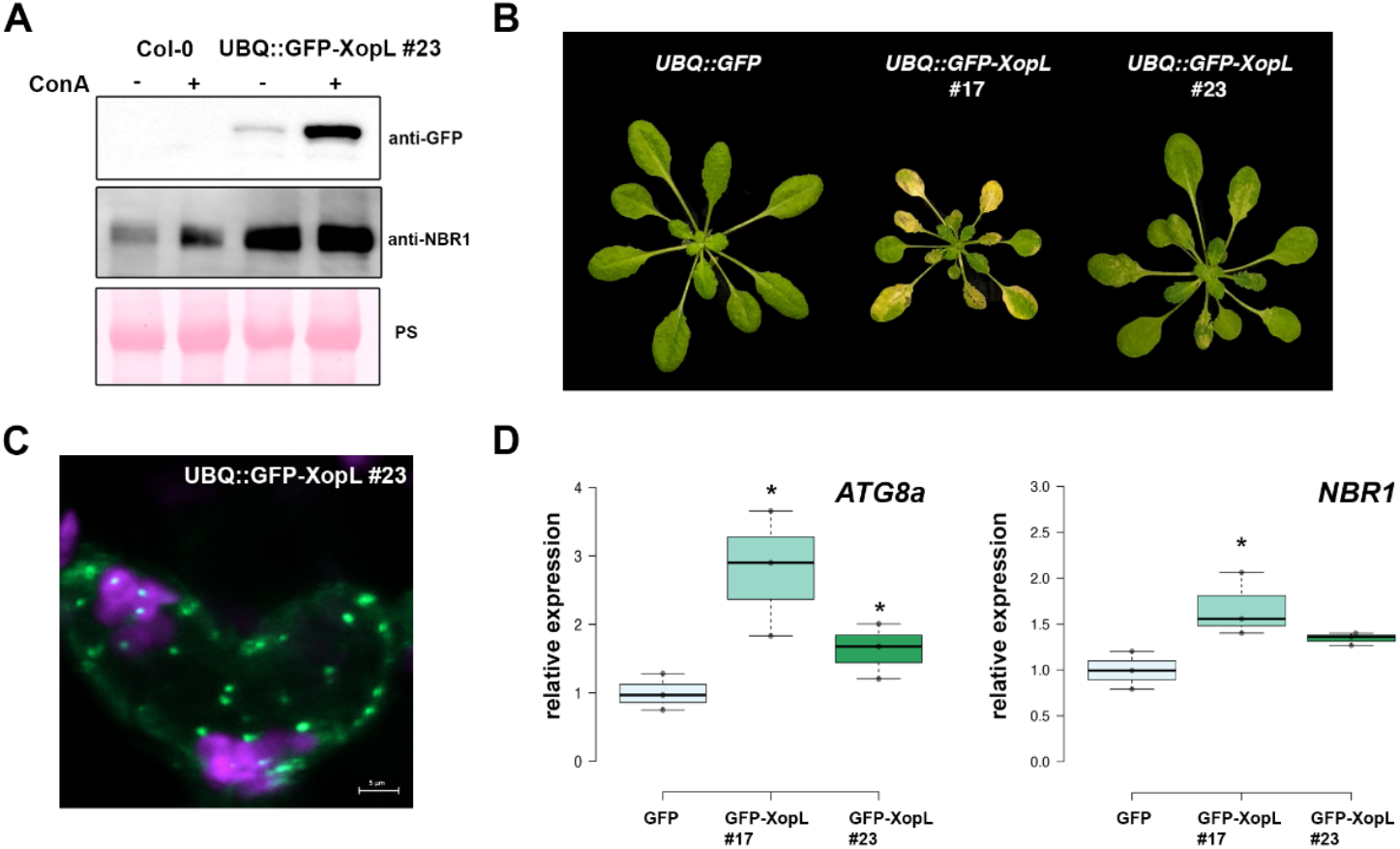
Transgenic *A. thaliana* GFP-XopL plants display defects in autophagic degradation. **(A)** Immunoblot analysis of NBR1 protein levels in transgenic UBQ::GFP-XopL plants or Col-0. Plants were treated with concanamycin A (ConA) for 6 hours. Expression of GFP-XopL was verified with an anti-GFP antibody. Ponceau S staining serves as a loading control. **(B)** 5 weeks old *A. thaliana* plants expressing UBQ::GFP-XopL develop an early senescence phenotype reminiscent of autophagy deficient mutants. **(C)** Localization analysis of GFP-XopL of transgenic *A. thaliana* UBQ::GFP-XopL #23 line. Image represents single confocal planes from abaxial epidermal cells (bars = 5 μm). **(D)** RT-qPCR analysis of *ATG8a* and *NBR1* transcript levels in *Arabidopsis thaliana* GFP or GFP-XopL plants. Values represent expression (n=3) relative to GFP control and were normalized to *PP2A*. Statistical significance (\**P*<0.05) was revealed by Student’s *t*-test.

**Fig. S8:**
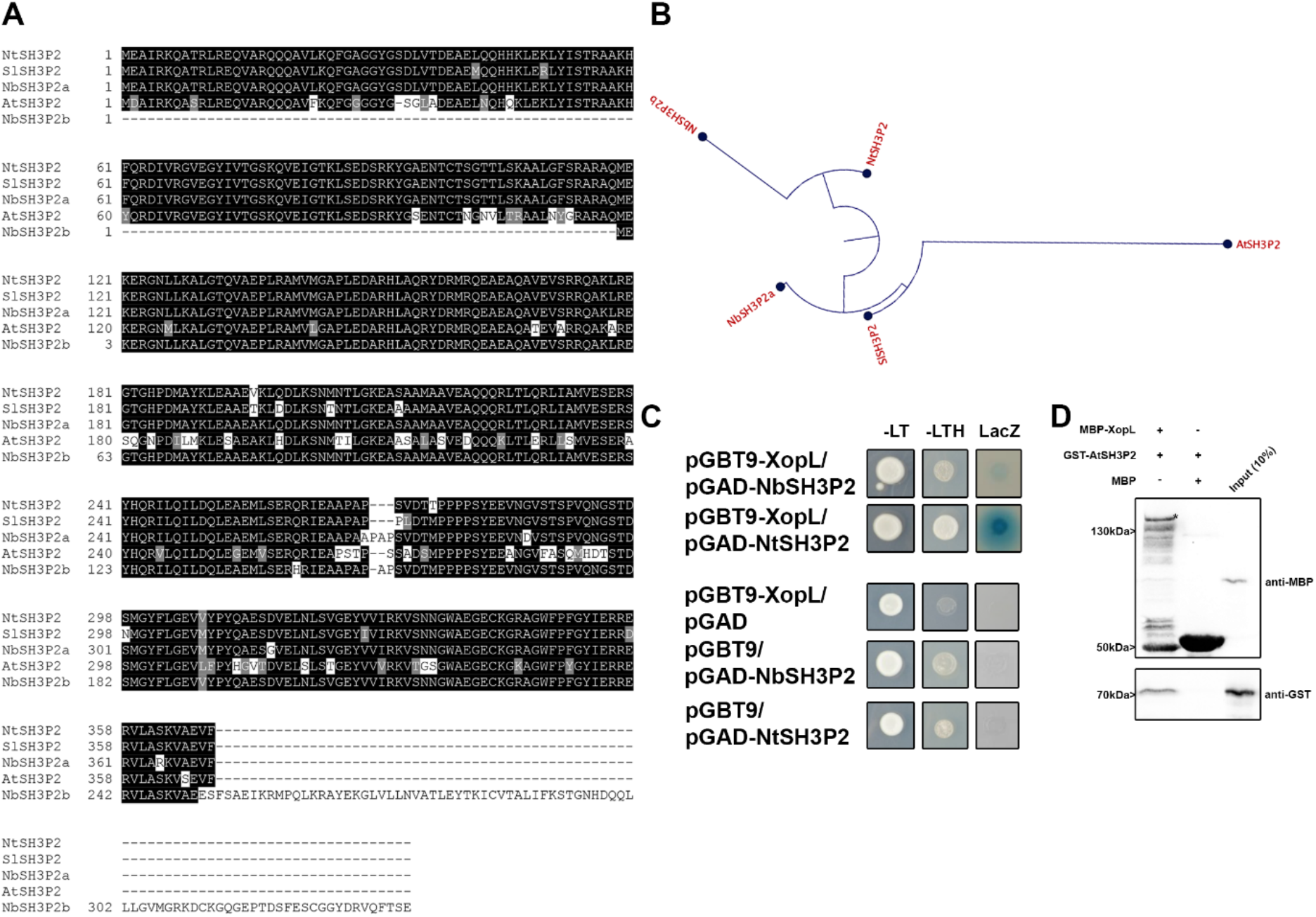
SH3P2 is conserved in different plant species. **(A)** Protein sequence alignment of SH3P2 from different species. The alignment was generated using CLUSTALW2 with default parameters and BoxShade 3.21. Positions of identical and similar sequences are boxed in black and grey, respectively. The following sequences were used to build the alignment: *Arabidopsis thaliana*, *Nicotiana tabacum, Nicotiana benthamiana*, *Solanum lycopersicum*. **(B)** Phylogentic analysis of the SH3P2 from different plant species. **(C)** XopL interacts with SH3P2 from *Nicotiana tabacum* and *Nicotiana benthamiana*. Interaction of XopL with SH3P2 in yeast two-hybrid assays. XopL fused to the GAL4 DNA-binding domain was expressed in combination with SH3P2 fused to the GAL4 activation domain (AD) in yeast strain Y190. Cells were grown on selective media before a LacZ filter assay was performed. The empty AD or BD vector served as negative control. NtSH3P2 = *Nicotiana tabacum* SH3P2, NbSH3P2 = *Nicotiana benthamiana* –LT = yeast growth on medium without Leu and Trp, –HLT = yeast growth on medium lacking His, Leu, and Trp, indicating expression of the HIS3 reporter gene. LacZ, activity of the lacZ reporter gene. **(D)** *In vitro* co-IP assay showing direct interaction of XopL with AtSH3P2. MBP-XopL and GST-AtSH3P2 were expressed in E. coli. Pull down was performed using amylose resin. Proteins were detected in an immunoblot using antibodies as indicated.

**Supp Video 1: XopL/SH3P2 puncta are mobile**

*Nicotiana benthamiana* leaf epidermal cells transiently expressing Venus^N173^-XopL in combination with AtSH3P2-Venus^C155^.

**Fig. S9:**
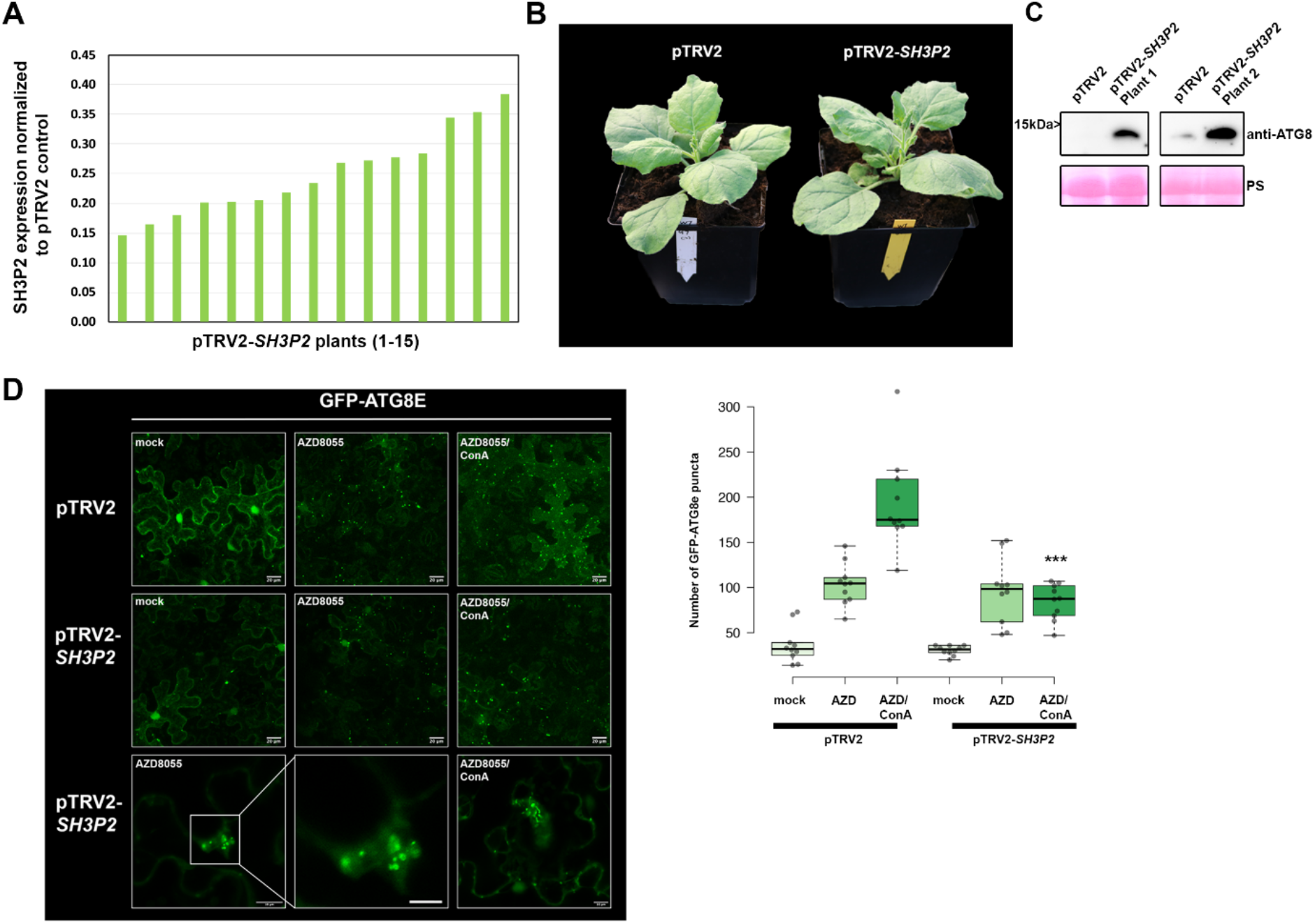
Silencing of SH3P2 in *N. benthamiana* perturbs autophagy. **(A)** qRT-PCR analysis of SH3P2 mRNA levels in silenced plants. *Actin* expression was used to normalize the expression value in each sample, and relative expression values were determined against pTRV2 control plants (set to 1). **(B)** Phenotype of SH3P2-VIGS plants in comparison to the pTRV2 control. Picture was taken 14 dpi. **(C)** Immunoblot analysis of ATG8 protein levels in *N. benthamiana* pTRV2 (control) and pTRV2-*SH3P2* (*SH3P2* silencing) 2 weeks after VIGS. Ponceau Staining (PS) served as a loading control. The experiment was repeated twice with similar results. **(D)** GFP-ATG8e-labeled autophagosomes were quantified from pTRV2 or pTRV2-SH3P2 plants transiently expressing GFP-ATG8e at 2dpi in the presence of autophagy inducer AZD = AZD8055 and AZD/ConA. Puncta were calculated from z-stacks of *n*=10 individuals using ImageJ. Statistical significance (*** *P* < 0.5) was revealed by Student’s *t*-test comparing number of autophagosomes in AZD/ConA treatments in pTRV2 and pTRV2-*SH3P2* plants. The experiment was repeated twice with similar results.

**Fig. S10:**
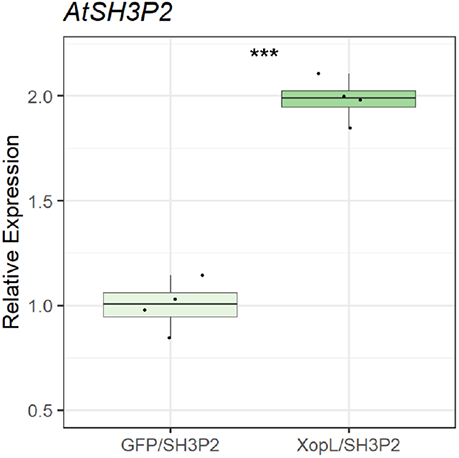
Gene expression of SH3P2 is induced by XopL and XopL-mediated degradation is due to post-transcriptional degradation events. **(A)** qRT-PCR analysis of AtSH3P2-specific mRNA levels in *N. benthamiana* plants transiently expressing AtSH3P2-HA, upon coexpression with GFP or GFP-XopL. *Actin* expression was used to normalize the expression value in each sample. Values represent *AtSH3P2* transcript level normalized to control (n=4).

**Fig. S11:**
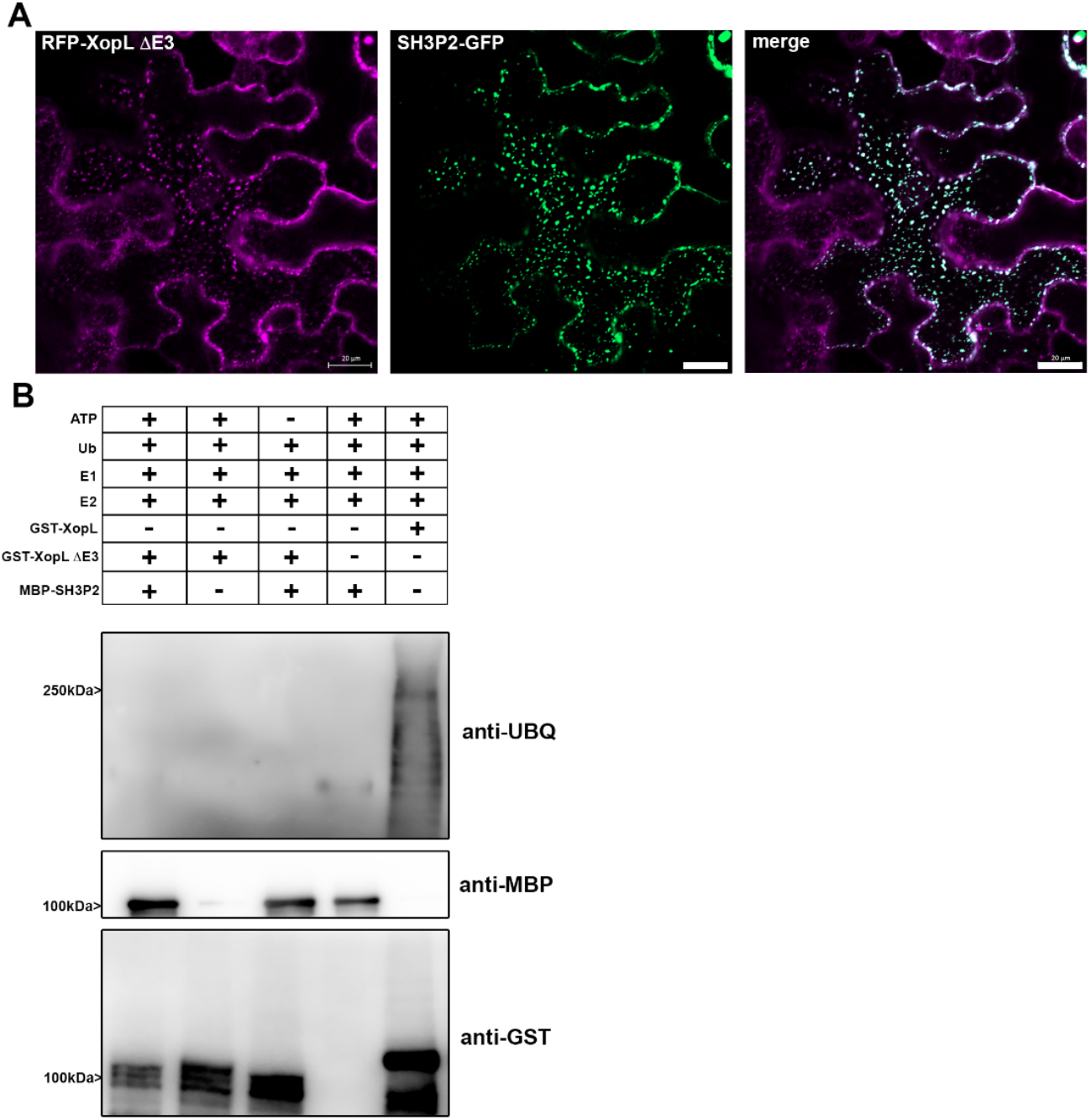
RFP-XopL ΔE3 co-localizes with and is unable to ubiquitinate SH3P2-GFP. **(A)** Colocalization analysis of RFP-XopL ΔE3 with SH3P2-GFP in *N. benthamiana* leaves. Imaging was performed 2 d after transient expression and images represent single confocal planes from abaxial epidermal cells (bars = 20 μm). **(B)** *In vitro* ubiquitination assay reveals GST-XopL ΔE3 is unable to ubiquitinate MBP-SH3P2. GST-XopL, ΔE3 and MBP-SH3P2 were tested using the Arabidopsis His-AtUBA1 and His-AtUBC8. Lanes 2 to 4 are negative controls, while Lane 5 is the positive control with GST-XopL. Proteins were separated by SDS-PAGE and detected by immunoblotting using the indicated antibodies. The experiment was repeated three times with similar results.

**Fig. S12:**
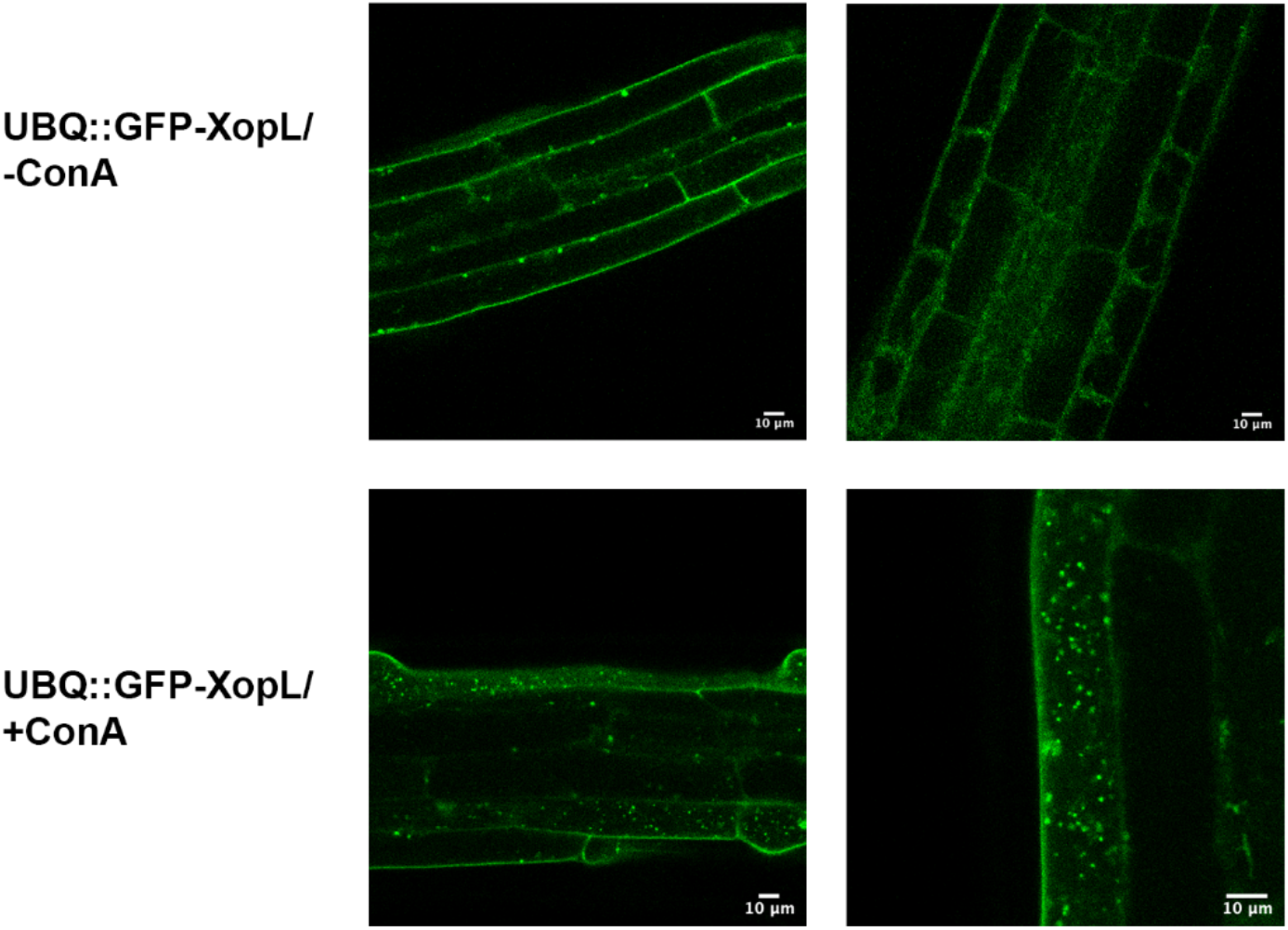
XopL is degraded in the vacuole. Localization of GFP-XopL in the presence or absence of ConA in transgenic GFP-XopL. DMSO or 0.5 μM ConA was used to treat seedlings, followed by confocal imaging of the roots. GFP-labeled puncta detectable upon ConA treatment indicate XopL accumulation in the vacuole (bars = 20 μm).

**Fig. S13:**
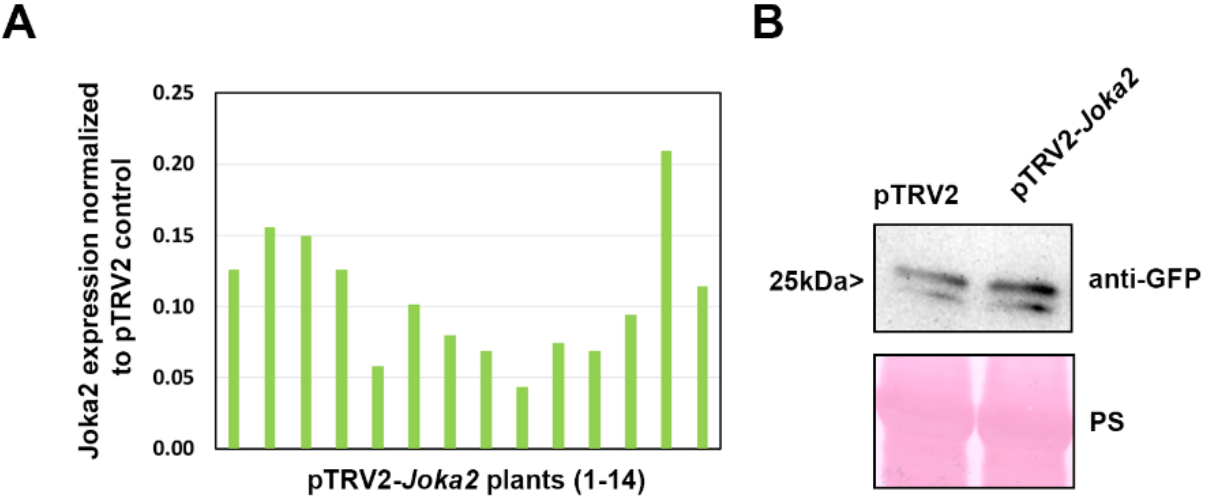
Virus-induced gene silencing of *Joka2* in *N. benthamiana* plants. **(A)** qRT-PCR analysis of Joka2 mRNA levels in *Joka2* silenced pepper plants. *Actin* expression was used to normalize the expression value in each sample, and relative expression values were determined against pTRV2 control plants (set to 1). **(B)** Immunoblot analysis of GFP and GFP-XopL protein levels in *N. benthamiana* plants silenced for *Joka2* (pTRV2-Joka2) compared against control (pTRV2). Ponceau staining (PS) served as a loading control.

**Fig. S14:**
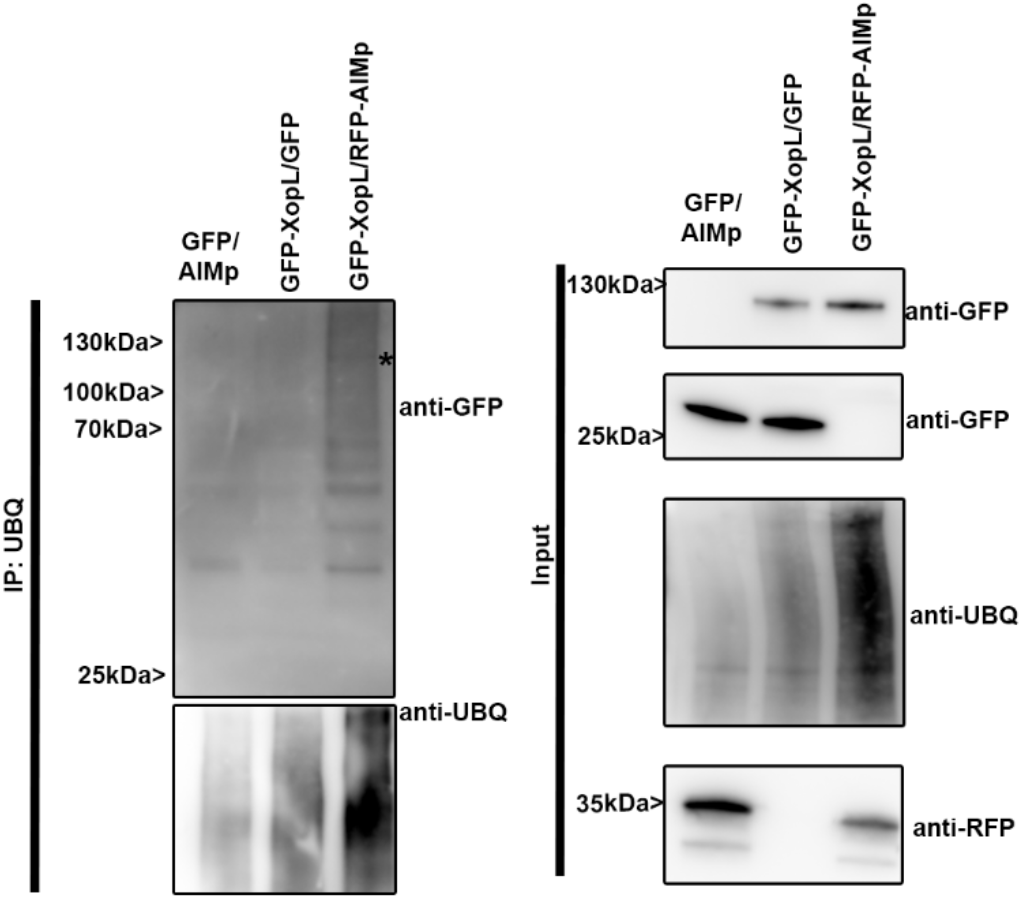
XopL in planta ubiquitination is enhanced by the presence of AIMp. GFP-XopL or GFP were transiently expressed in *N. benthamiana*. RFP-AIMp was co-infiltrated. Samples were taken 48 hpi, and total proteins (Input) were subjected to immunoprecipitation (IP) with the ubiquitin pan selector, followed by immunoblot analysis of the precipitates using either anti-GFP or anti-ubiquitin antibodies. RFP-AIMp expression was verified by an anti-RFP antibody. Asterisk indicates the GFP-XopL full-length protein. The experiment was repeated twice with similar results.

**Fig. S15:**
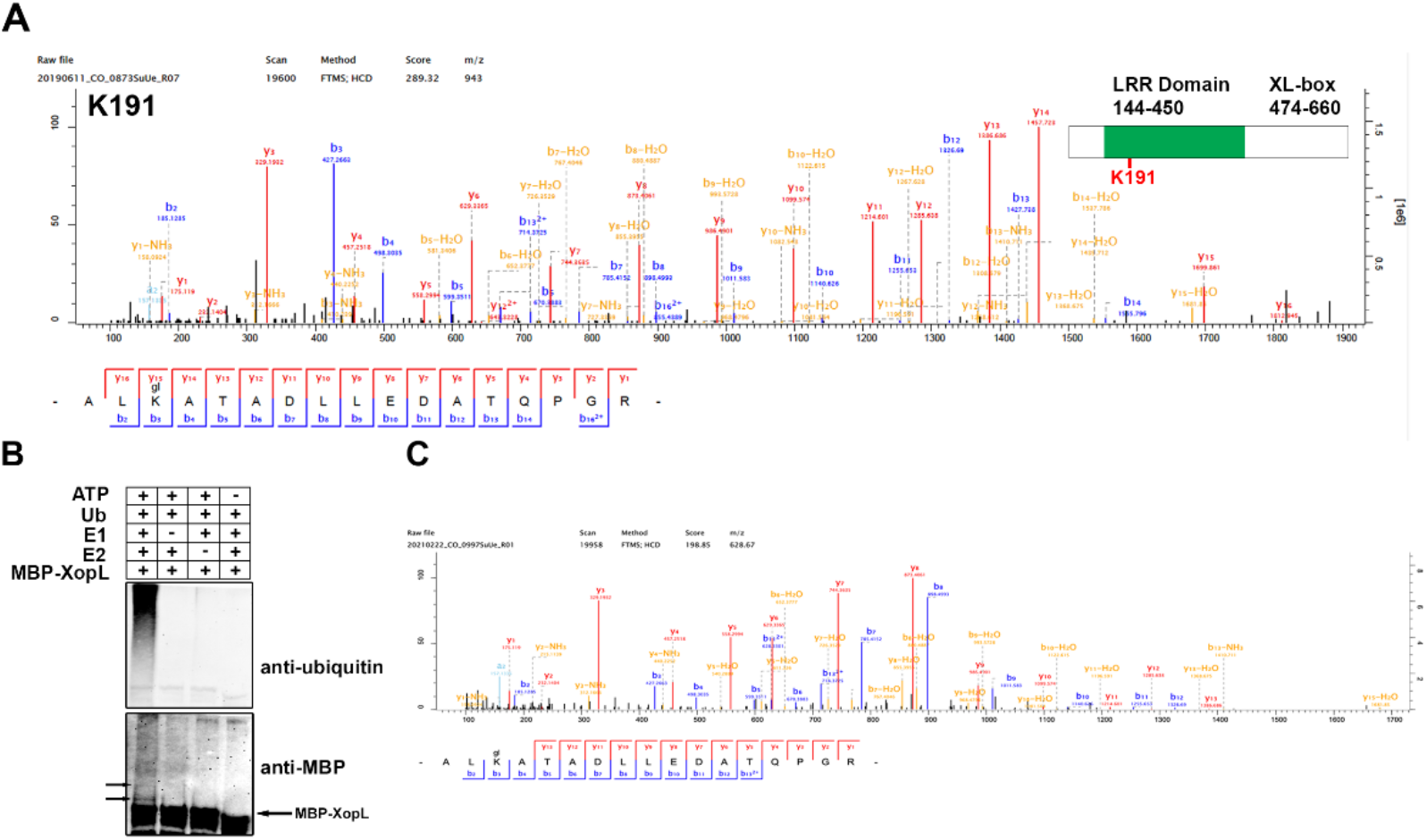
XopL is ubiquitinated *in planta* and undergoes self-ubiquitination. **(A)** XopL ubiquitination site at lysine 191 was identified in vivo by LC-MS/MS. GFP-XopL was transiently expressed in *N. benthamiana* and total proteins were subjected to anti-GFP IP followed by trypsin digestion. Ubiquitinated peptides were detected by LC-MS/MS. The spectrum shows the fragmentation pattern of the GlyGly modified peptide ALglKATADLLEDATQPGR corresponding to amino acids 189-205. **(B)** *In vitro* ubiquitination assay reveals autoubiquitination of XopL. Ubiquitination of MBP-XopL was tested using the Arabidopsis His-AtUBA1 and His-AtUBC8. Lanes 2 to 4 are negative controls. Proteins were separated by SDS-PAGE and detected by immunoblotting using the indicated antibodies. Arrows in the MBP blot indicate higher molecular weight bands of MBP-XopL and autoubiquitination events. The experiment was repeated twice with similar results. **(C)** XopL ubiquitination site at lysine 191 was identified in vitro by LC-MS/MS. GST-XopL was used in an in vitro ubiquitination assay and samples were subjected to trypsin digestion. Ubiquitinated peptides were detected by LC-MS/MS. The spectrum shows the fragmentation pattern of the GlyGly modified peptide ALglKATADLLEDATQPGR corresponding to amino acids 189-205.

**Fig. S16:**
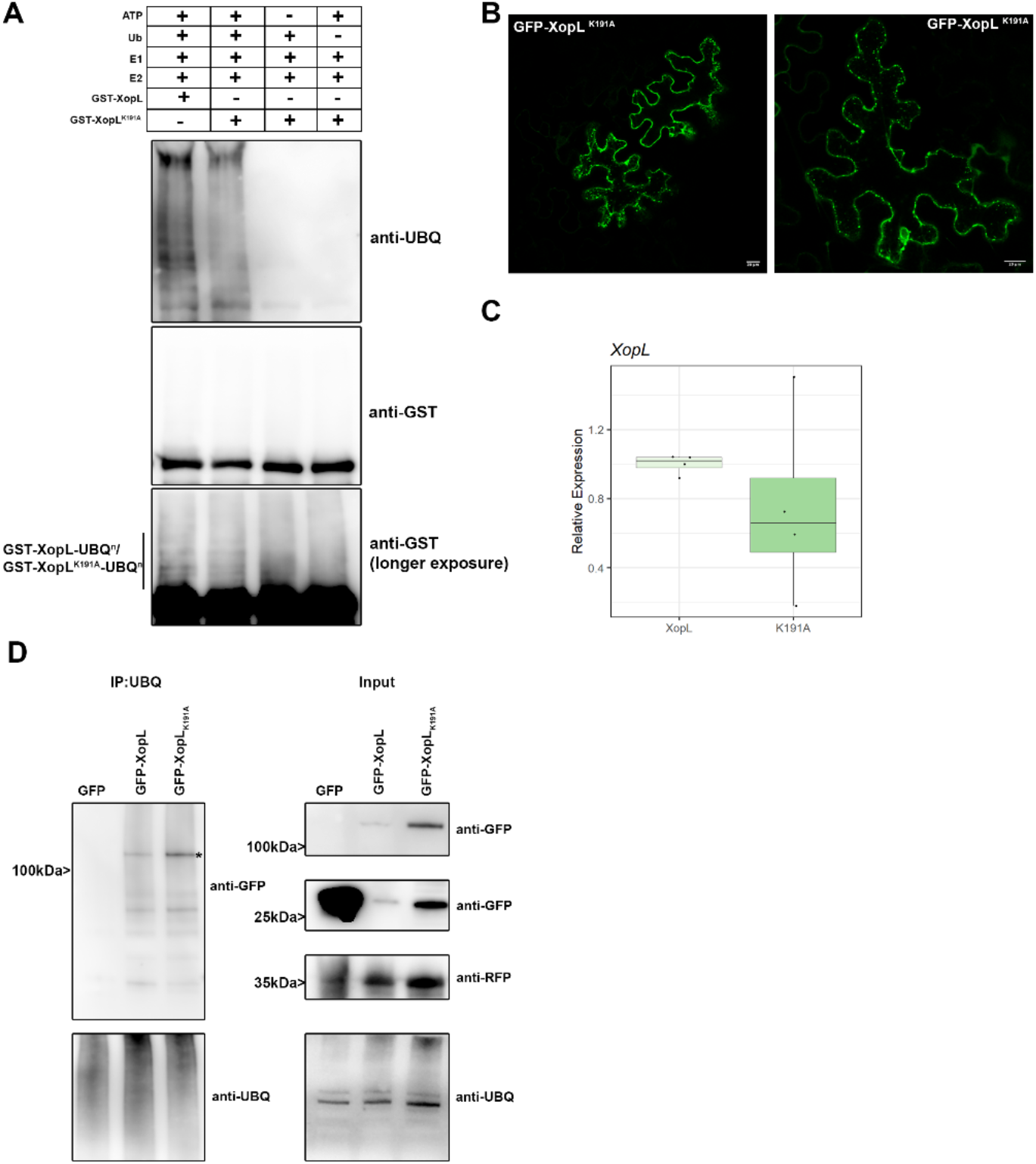
Characterization of XopL K191A variant in vitro and in planta. **(A)** *In vitro* ubiquitination assay reveals less autoubiquitination of XopL K191A compared to XopL WT. Ubiquitination of GST-XopL was tested using the Arabidopsis His-AtUBA1 and His-AtUBC8. Lanes 3 to 5 are negative controls. Proteins were separated by SDS-PAGE and detected by immunoblotting using the indicated antibodies. The experiment was repeated twice with similar results. **(B)** Localization analysis of GFP-XopL_K191A_ in *N. benthamiana* leaves. Imaging was performed 2 d after transient expression and images represent single confocal planes from abaxial epidermal cells (bars = 20 μm). **(C)** qRT-PCR analysis of GFP-XopL and GFP-XopL K191A mRNA levels in *N. benthamiana* plants during transient expression. *Actin* expression was used to normalize the expression value in each sample. **(D)** GFP-XopL and GFP-XopL_K191A_ were transiently expressed in *N. benthamiana*. RFP-AIMp was co-infiltrated. Samples were taken 48 hpi, and total proteins (Input) were subjected to immunoprecipitation (IP) with the ubiquitin pan selector, followed by immunoblot analysis of the precipitates using either anti-GFP or anti-ubiquitin antibodies. GFP served as a control. RFP-AIMp expression was verified by an anti-RFP antibody. Asterisk indicates the GFP-XopL full-length protein. The experiment was repeated twice with similar results.

**Fig. S17:**
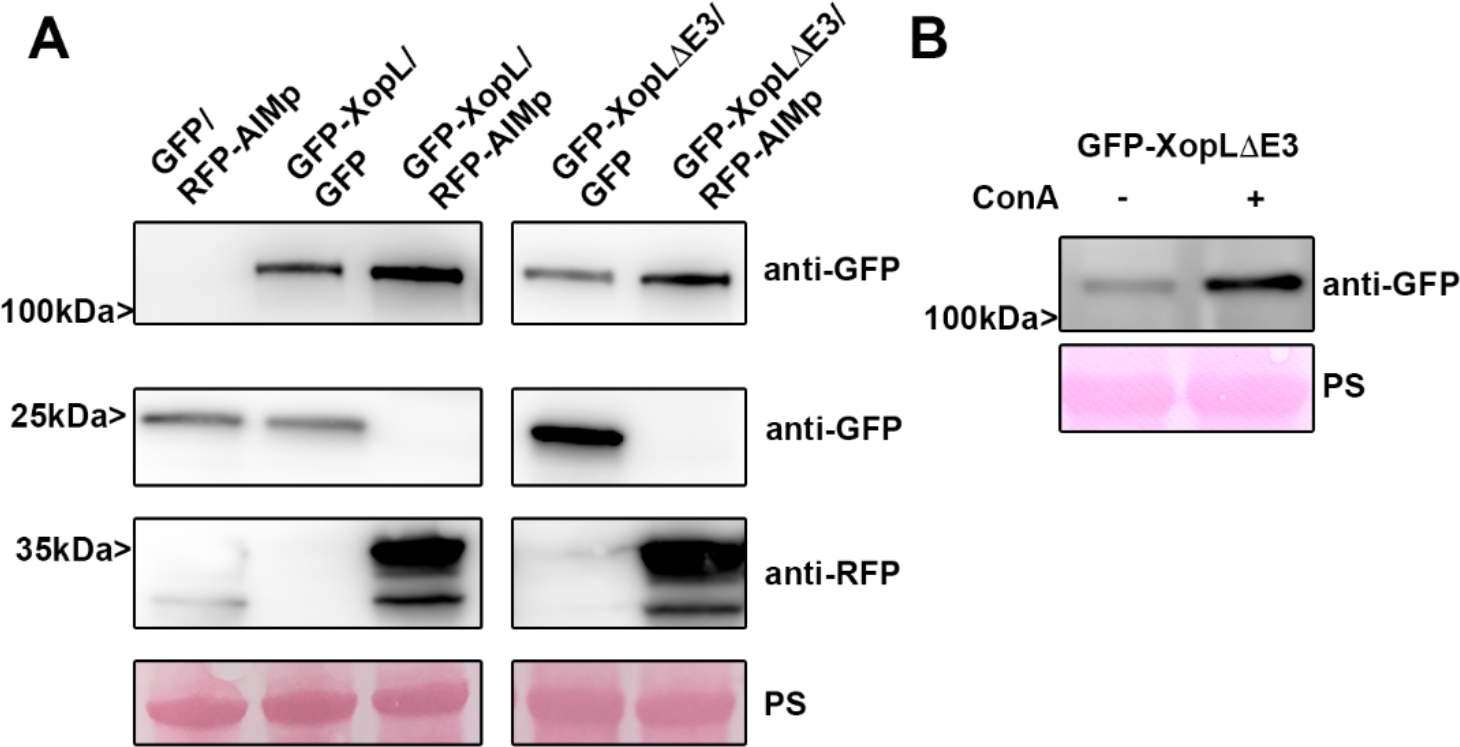
XopL ΔE3 is degraded by autophagy. **(A)** GFP, GFP-XopL, or GFP-XopL ΔE3 was coexpressed with RFP-AIMp or RFP control in *N. benthamiana*. Samples were taken at 2 dpi, total proteins extracted and immunoblotted using the indicated antibodies. Ponceau Staining (PS) served as a loading control. **(B)** Immunoblot of transiently expressed GFP-XopL ΔE3 in *N. benthamiana* after treatment of ConA or DMSO carrier. Ponceau Staining (PS) served as a loading control.

**Fig. S18:**
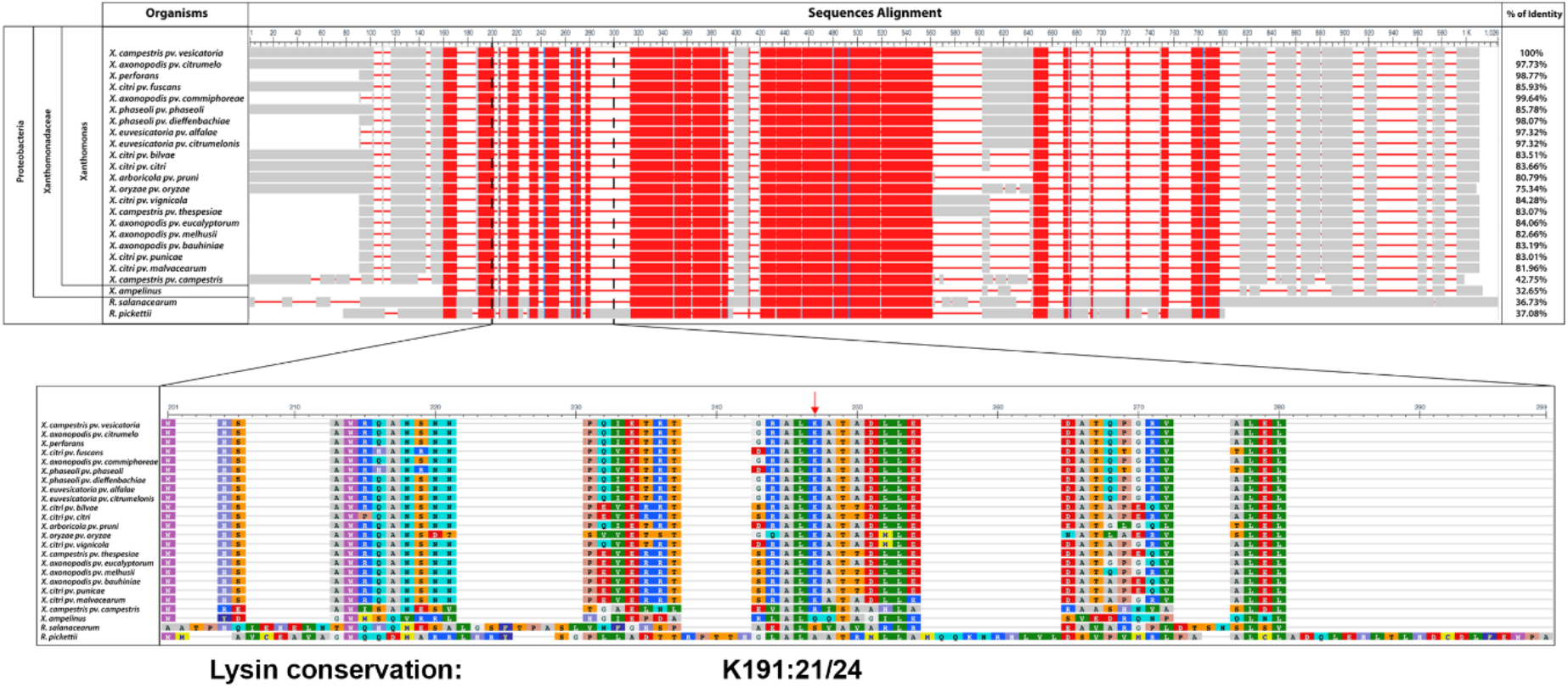
XopL K191 residue is highly conserved through the Xanthomonas genus. **(A)** Sequence alignment of XopL protein from *Xcv* with related effectors from *Xanthomonas* genus or more distantly related plant pathogen bacteria. Colors represent amino acid conservation through the alignment, with red for highly conserved residues, blue for lower conservation and grey for no conservation. Identical amino acid percentage to XopL*^Xcv^* is displayed to the right of the alignment. **(B)** Closer view of the region 201-299 of the alignment with amino acids colored according to the Rasmol coloration. Lysine K191 is indicated with a red arrow.

**Supplemental Table 1**: Primers used in manuscript.

