## Supplemental Table 1 for "A bacterial effector counteracts host autophagy by promoting degradation of an autophagy component"

| **Method/Name** | **Sequence** |
|---|---|
| **qPCR** | |
| *NbActin* Fwd | 5'-AAAGACCAGCTCATCCGTGGAGAA-3' |
| *NbActin* Rev | 5'-TGTGGTTTCATGAATGCCAGCAGC-3' |
| *NbATG8-2.1/2* Fwd | 5'-CACCCACTTGAAAGGCGACAGGC-3' |
| *NbATG8-2.1/2* Rev | 5'-GCCTTCTCAGCACTAAGCTTTATTCTC-3' |
| *NbATG8-1.1/2* Fwd | 5'-CTTGAGAAGAGGCGTGCTGAAGC-3' |
| *NbATG8-1.1/2* Rev | 5'-ATCTGTTGGTGGTAGGACATTATCAAC-3' |
| *NbJoka2* Fwd | 5'-CGTTGTGATGGTTGTGGTGT-3' |
| *NbJoka2* Rev | 5'-AGGGACGCCGGTAAGTTAAA-3' |
| *NbSH3P2* Fwd | 5'-TACTATGCCTCCACCTCCCT-3' |
| *NbSH3P2* Rev | 5'-TTTGCATTCACCTTCAGCCC-3' |
| *NbJoka2* Fwd | 5'-CGCAGCTGGATCTGACATTA-3' |
| *NbJoka2* Rev | 5'-CGGGAGCAGAAGAACTTGAC-3' |
| *NbATG7* Fwd | 5'-ATGGCGGATAGTGGAAGAGG-3' |
| *NbATG7* Rev | 5'-CCTGACTAGCTAGCCGAGAC-3' |
| *NBR1* Fwd | 5'-GAGGACCCAGACCGGAAGG-3' |
| *NBR1* Rev | 5'-GACAAACACGACGAGGATGC-3' |
| *ATG8a* Fwd | 5'-CAAGCTTGGAGCTGAGAAAG-3' |
| *ATG8a* Rev | 5'-GCAACGGTAAGAGATCCAAA-3' |
| **Cloning** | |
| *NbSH3P2*-VIGS_cloning_Fwd | 5'-CTAGCTCAAAGATATGATAGAATGCGA-3' |
| *NbSH3P2*-VIGS_cloning_Rev | 5'-GTGGTAGCTGATCAAGAATCTGAAGGAT-3' |
| *XopL* Fw | 5'-CACCATGCGACGCGTCGATCAAC-3' |
| *XopL* Rev | 5'-CTACTGATGGCCTGAAGGTTCCGG-3' |
| *AtSH3P2* Fwd (with STOP) | 5'-CACCATGGATGCAATTAGAAAACAAGC-3' |
| *AtSH3P2* Rev | 5'-TCAGAAAACTTCGGACACTTTGCTAGC-3' |
| *AtSH3P2* Fwd (wihout STOP) | 5'-CACCAACAATGGATGCAATTAGAAAACAAGC-3' |
| *AtSH3P2* Rev | 5'-GAAAACTTCGGACACTTTGCTAGCAAG-3' |
| *NbSH3P2* Fwd (with STOP) | 5'-GAATTCATGGAAGCAATCAGAAAGCAAGC-3' |
| *NbSH3P2* Rev | 5'-GTCGACTCAGAAAACCTCGGCAACTTTCC-3' |
| *SlJoka2* Fwd | 5'-GGGGACAAGTTTGTACAAAAAAGCAGGCTTC ATGTGTGAGTTGGGGCTAT-3' |
| *SlJoka2* Rev | 5'-GGGGACCACTTTGTACAAGAAAGCTGGGTTCTACTGCTCTCCAGCAATAA-3' |
| **Site-directed Mutagenesis** | |
| XopL K191A sense | 5'-CGCACAGGCCGGGCGCTGGCGGCGACAGCCGACCTGCTGGA-3' |
| XopL K191A antisense | 5'-CTCCAGCAGGTCGGCTGTCGCCGCCAGCGCCCGGCCTGTGCG-3' |
| XopLH584A L585A G586E sense | 5'-CAGCACATGACGATCGACGCGGCAGGGTTGATGGAGCTCCCGGA-3' |
| XopLH584A L585A G586E antisense | 5'-TCCGGGAGCTCCATCAACCCTGCCGCGTCGATCGTCATGTGCTG-3' |
| **Construction of null mutants and complementation in Xcv** | |
| XopQ up Fwd | 5'-GCCCTTTCGTCTTCAAGGCCGCCTGCGCCTGCTCA-3' |
| XopQ up Rev | 5'-CGGAGCGCGGGAGGACCTCTGGCAGTGAAAG-3' |
| XopQ down Fwd | 5'-TCCTCCCGCGCTCCGCAGCTTTCGC-3' |
| XopQ down Rev | 5'-AAGCTGTCAAACATGAGGCTGTATCCGGCCGGTTG-3' |
| XopL up Fwd | 5'-GCCCTTTCGTCTTCAAGGCGGATTGACTGCGGTTG-3' |
| XopL up Rev | 5'-TGGCTCTCGCATTCCTCGTCTTGGC-3' |
| XopL down Fwd | 5'-GGAATGCGAGAGCCAACGCGACAGGC-3' |
| XopQ down Rev | 5'-AAGCTGTCAAACATGAGTGCCGCCAATCAGGAAGC-3' |
| XopL comp Fwd | 5'-GGCACGACAGGTTTGATGGCAACGCGTGCTGACGC-3' |
| XopL comp Rev | 5'-TGGTGATGATGGTGGATCTGATGGCCTGAAGGTTCCG-3' |
