## Supporting information for "A bacterial effector counteracts host autophagy by promoting degradation of an autophagy component"

**Figure S1: S**ilencing of *ATG7* in *N. benthamiana* plants abolishes autophagosome formation and Xcv blocks autophagy at 6hpi.
**Figure S2:** Virus induced gene silencing of *ATG7* in *N. benthamiana* is beneficial for Xcv.
**Figure S3:** Suppression of autophagy is enhanced by T3Es**.**
**Figure S4:** Screening for Xanthomonas T3Es with altered autophagic flux.
**Figure S5:** XopL contributes to Xcv virulence.
**Figure S6:** Joka2 bodies are induced during Xcv infection in a XopL-dependent manner.
**Figure S7:** Transgenic *A. thaliana* GFP-XopL plants display defects in autophagic degradation.
**Figure S8**: SH3P2 is conserved in different plant species.
**Figure S8:** XopL is ubiquitinated *in planta*.
**Supplemental Video 1:** XopL/SH3P2 puncta are mobile.
**Figure S9:** Silencing of SH3P2 in *N. benthamiana* perturbs autophagy.
**Figure S10:** Gene expression of SH3P2 is induced by XopL and XopL-mediated degradation is due to post-transcriptional degradation events.
**Figure S11:** RFP-XopL ΔE3 co-localizes with and is unable to ubiquitinate SH3P2-GFP.
**Figure S12:** XopL is degraded in the vacuole.
**Figure S13:** Virus-induced gene silencing of *Joka2* in *N. benthamiana* plants.
**Figure S14:** XopL in planta ubiquitination is enhanced by the presence of AIMp.
**Figure S15:** XopL is ubiquitinated *in planta* and undergoes self-ubiquitination.
**Figure S16**: Characterization of $XopL_{K191A}$ variant i*n vitro* and *in planta*.
**Figure S17:** XopL ΔE3 is degraded by autophagy.
**Figure S18:** XopL K191 residue is highly conserved through the Xanthomonas genus
**Supplemental Table 1**: Primers used in manuscript.

**Supplemental Figure 1**

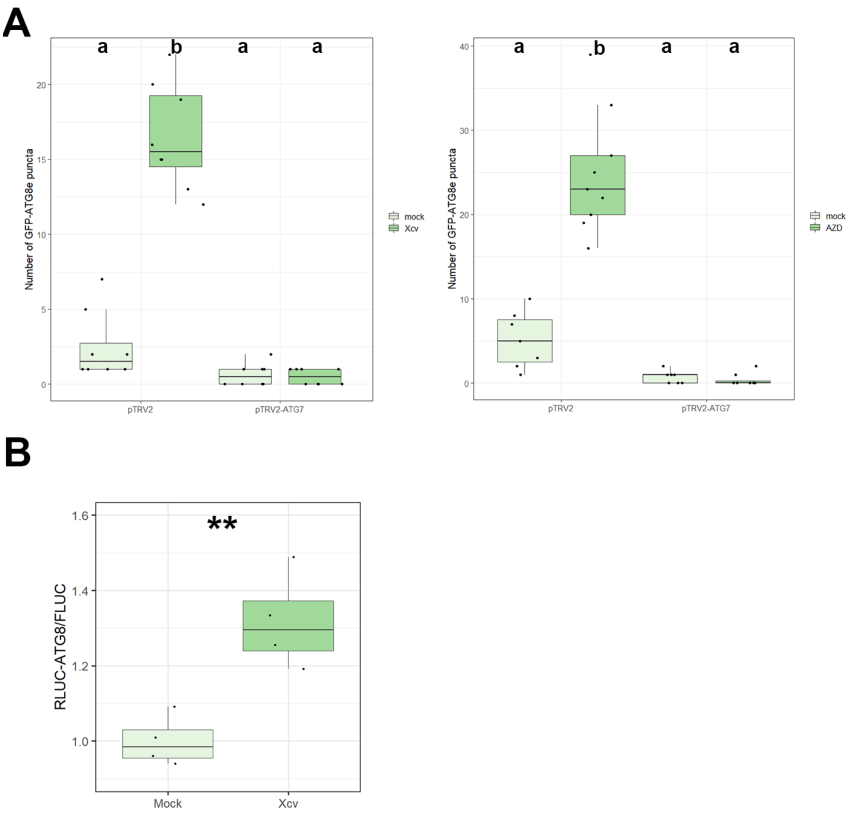


**Fig. S1: Silencing of *ATG7* in *N. benthamiana* plants abolishes autophagosome formation and Xcv blocks autophagy at 6hpi.** GFP-ATG8e-labeled puncta were quantified from plants silenced for *ATG7* (pTRV2-ATG7) infected with mock or Xcv *ΔxopQ* at 6hpi in the presence or absence of ConA, and of AZD. Puncta were calculated from z-stacks (X) of $n$=12 individuals using ImageJ. Different letters indicate statistically significant different groups ($P$ < 0.05) as determined by one way ANOVA.
**(B)** RLUC-ATG8a or RLUC-NBR1 constructs were coexpressed with internal control FLUC in *N. benthamiana*. *Xcv ΔxopQ* or infiltration buffer (mock) was co-infiltrated with Agrobacteria containing the respective constructs. RLUC and FLUC activities were simultaneously measured in leaf extracts at 8 h post- infiltration using the dual-luciferase system. Values represent the ratio of RLUC-ATG8a and FLUC activities (n=4). Statistical significance (***$P$ < 0.001) was revealed by Student's $t$-test. The experiment was repeated 3 times with similar results.

**Supplemental Figure 2**

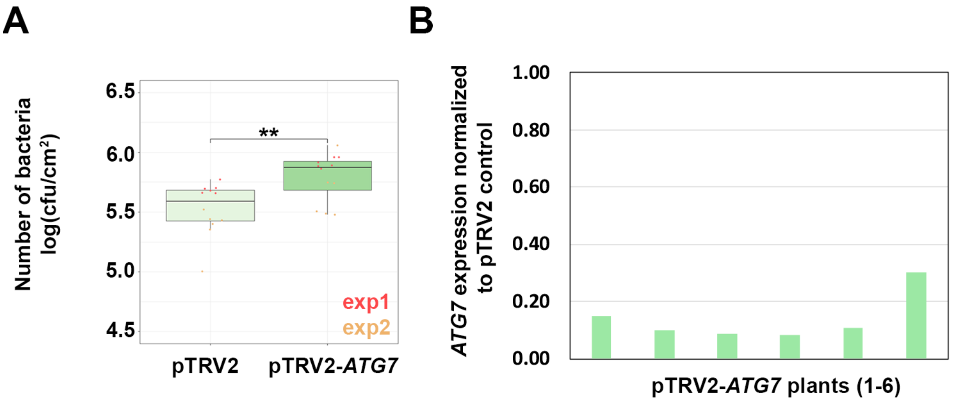


**Fig. S2: Virus induced gene silencing of *ATG7* in *N. benthamiana* is beneficial for Xcv.**

**(A)** Growth of Xcv *ΔxopQ* in *N. benthamiana* plants silenced for *ATG7* (pTRV2-*ATG7*) compared to control plants (pTRV2). Leaves were dip-inoculated with a bacteria suspension at $OD_{600}$ = 0.2 and bacteria were quantified at 6 dpi. Data represent the mean SD (n = 6). Significant differences were calculated using Student's *t-test* and are indicated by **, P < 0.01. The experiment was repeated twice with similar trends. Red and yellow data points represent repeats of the experiment.
**(B)** qRT-PCR analysis of *ATG7* mRNA levels in silenced *N. benthamiana* plants. *Actin* expression was used to normalize the expression value in each sample, and relative expression values were determined against the mean expression in pTRV2 (control) plants.

**Supplemental Figure 3**

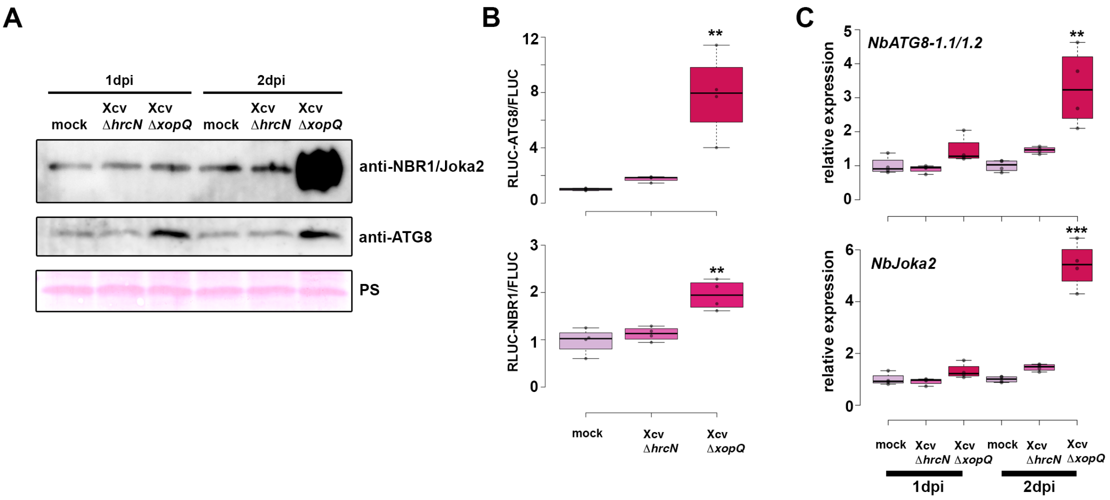


**Fig. S3: Suppression of autophagy is enhanced by T3Es.**
**(A)** Immunoblot analysis of NBR1 and ATG8 protein levels in Xcv *ΔxopQ, ΔhrcN* or mock infected *N. benthamiana* plants at 1 and 2dpi. Ponceau Staining (PS) served as a loading control. The experiment was repeated twice with similar results.
**(B)** Autophagic flux determined by quantitative dual-luciferase assay. RLUC-ATG8a or RLUC-NBR1 constructs were coexpressed with internal control FLUC in *N. benthamiana*. Xcv *ΔxopQ* and *ΔhrcN* were respectively co-infiltrated with Agrobacteria containing the luciferase constructs. *Renilla* and *Firefly* luciferase activities were simultaneously measured in leaf extracts at 48 h post- infiltration using the dual-luciferase system. Values represent the ratio of RLUC-ATG8a and FLUC activities normalized to mock (n=4). Statistical significance (***P*<0.01) was revealed by Student's *t*-test. The experiment was repeated 2 times with similar results.
**(C)** RT-qPCR analysis of *NbATG8-1.1/2* and *NbJoka2* transcript levels upon challenge of *N. benthamiana* plants with Xcv *ΔxopQ* and *ΔhrcN* for 1 and 2 dpi compared to mock infected plants. Values represent expression relative to mock control of respective time point and were normalized to *actin*. Statistical significance (*** *P* < 0.001) was revealed by Student's *t*-test.

**Supplemental Figure 4**

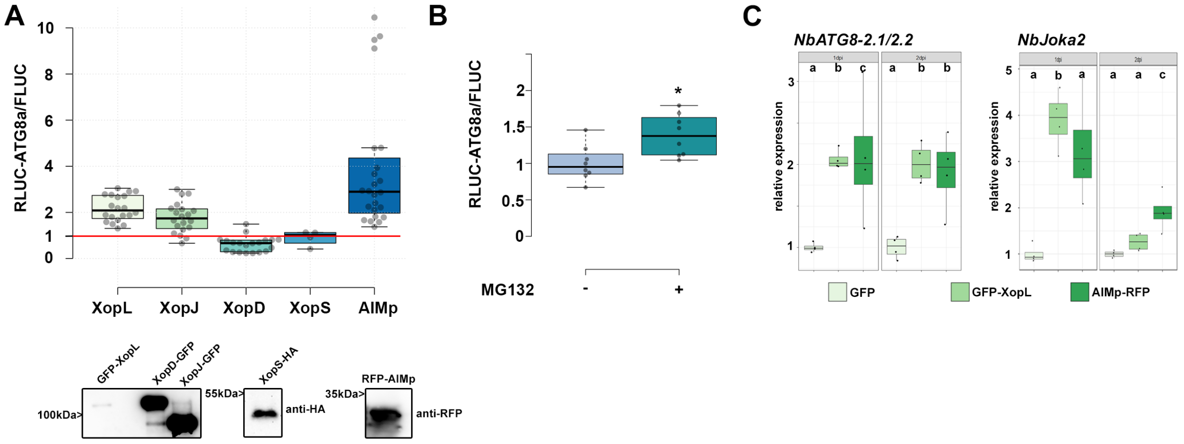


**Fig. S4: Screening for Xanthomonas T3Es with altered autophagic flux.**

**(A)** RLUC-ATG8a constructs were coexpressed with internal control FLUC in *N. benthamiana*. GFP-XopL, XopJ-GFP, XopD-GFP and XopS-HA were co-infiltrated with Agrobacteria carrying the RLUC-ATG8a and FLUC constructs. Renilla and Firefly luciferase activities were simultaneously measured in leaf extracts at 48 h post- infiltration using the dual-luciferase system. Values represent the ratio of RLUC-ATG8a to FLUC activity normalized to GFP control (XopL, XopJ, XopD, AIMp; n=20; XopS n=4). Expression of T3Es and RFP-AIMp were verified with the indicated antibodies.

**(B)** RLUC-ATG8a constructs were coexpressed with internal control FLUC in *N. benthamiana*. Plants were treated with MG132 for 6 hours prior measurement. Values represent the ratio of RLUC-ATG8a to FLUC activity normalized to vector control (n=8). Statistical significance (* $P < 0.5$) was revealed by Student's *t*-test.

**(C)** RT-qPCR analysis of *NbATG8-2.1/2* and *NbJoka2* transcript levels upon Agrobacteria-mediated transient expression of GFP, GFP-XopL or AIMp for 1 and 2 dpi. Values represent expression relative to GFP control of respective time point and were normalized to *actin*. Statistical significantly different groups are denoted by different letters, as calculated using Kruskal-Wallis rank sum test ($P$<0.05).

**Supplemental Figure 5**

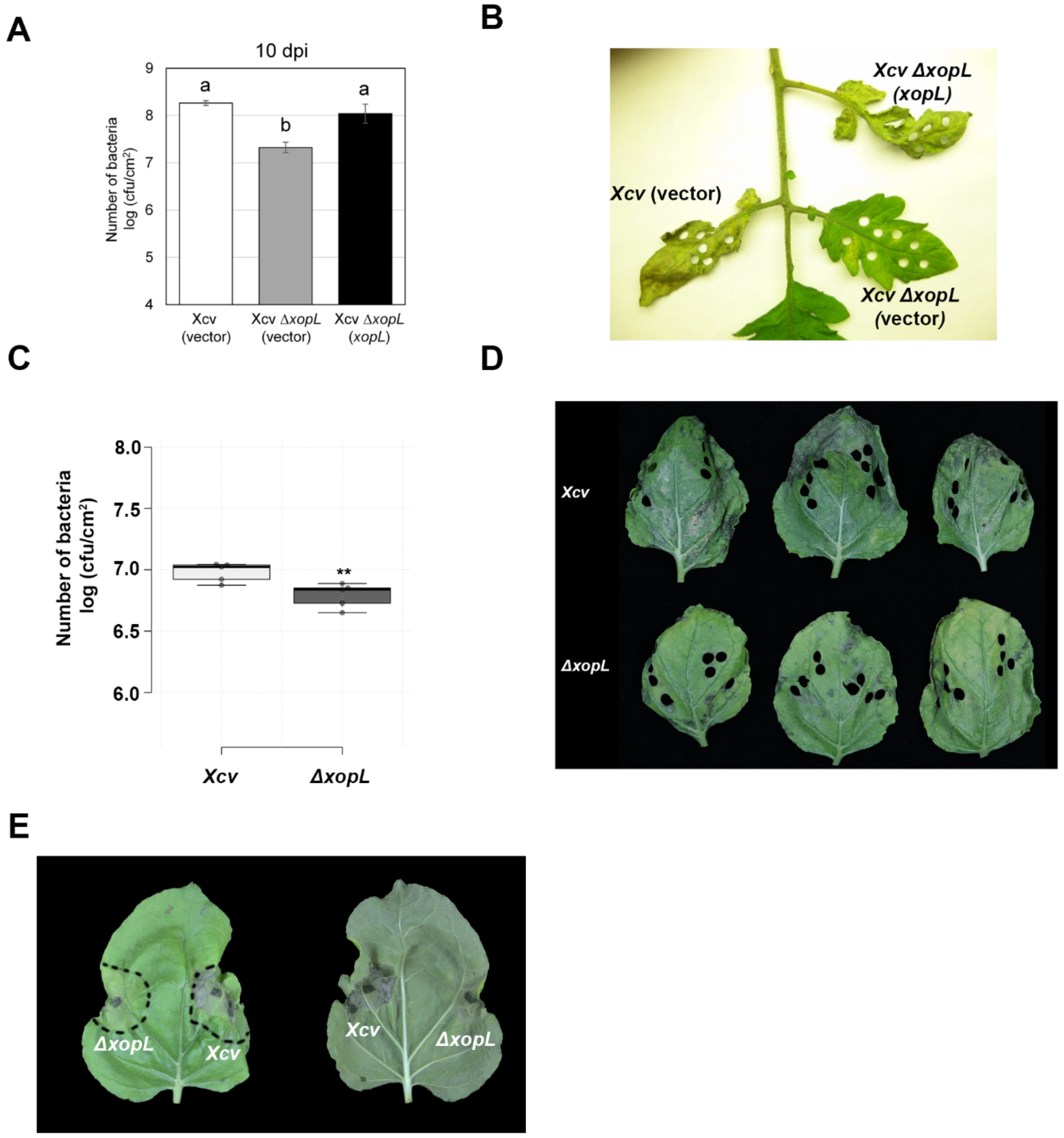


**Fig. S5: XopL contributes to Xcv virulence.**
**(A)** Growth of Xcv 85-10 (vector) (white bar), Xcv 85-10 Δ*xopL* (vector) (grey bar), and Xcv 85-10 Δ*xopL* (*xopL*) (black bar) strains in tomato VF36 leaves. Leaves were dipped in a 2 x $10^8$ CFU/mL suspension of bacteria. The number of bacteria in each leaf was quantified at 10 dpi. Data points represent mean $\log_{10}$ colony-forming units per $cm^2 \pm$ SD of three plants. Different letters above bars indicate statistically significant (Tukey's honestly significant difference (HSD) test, $P < 0.05$) differences between samples. Vector = pBBR1MCS-2.
**(B)** Delayed disease symptom development in tomato leaves inoculated with *Xcv* or *Xcv* Δ*xopL*. Tomato leaves inoculated with strains described in (A) were photographed at 14 dpi.
**(C)** Growth of Xcv 85-10, Xcv 85-10 Δ*xopL* strains in *roq1 N. benthamiana* leaves. Leaves were dipped in a 2 x $10^8$ CFU/mL suspension of bacteria. The number of bacteria in each leaf was quantified at 10 dpi (n = 5). Significant differences were calculated using Student's *t-test* (**, $P < 0.01$). The experiment was repeated twice with similar trends.

**(D)** Delayed disease symptom development in *roq1 N. benthamiana* leaves dip-inoculated with *Xcv* or *Xcv ΔxopL*. *N. benthamiana roq1* leaves inoculated with strains described in (C) were photographed at 10 dpi.
**(E)** Delayed symptom development in *roq1 N. benthamiana* leaves inoculated with *Xcv* or *Xcv ΔxopL*. Leaves were syringe-inoculated with $OD_{600}$=0.2 and photographed at 3 dpi.

**Supplemental Figure 6**

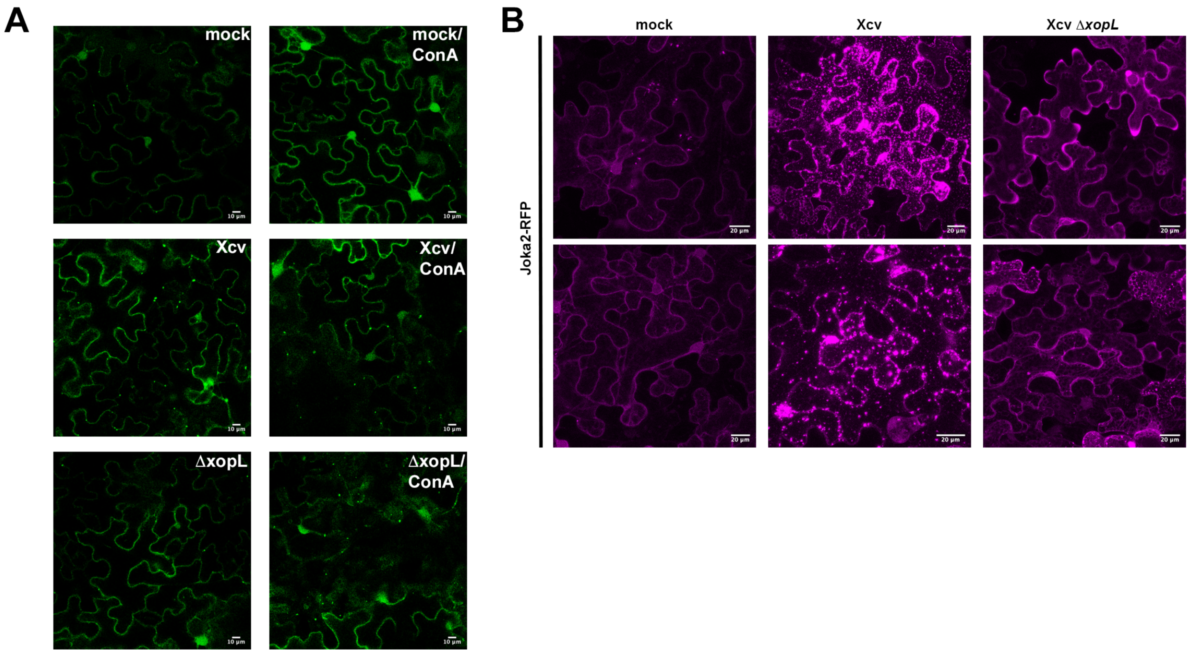


**Fig. S6: Joka2 bodies are induced during Xcv infection in a XopL-dependent manner.**
**(A)** GFP-ATG8e-labeled autophagosomes imaged from *N. benthamiana* plants infected with mock, Xcv or *Xcv ΔxopL* at 2 dpi in the presence or absence of ConA (bars = 10 µm).
**(B)** RFP-Joka2 labelled puncta or aggregates upon challenge of *N. benthamiana* leaves with mock, *Xcv* or *Xcv ΔxopL* infection at 1dpi.

**Supplemental Figure 7**

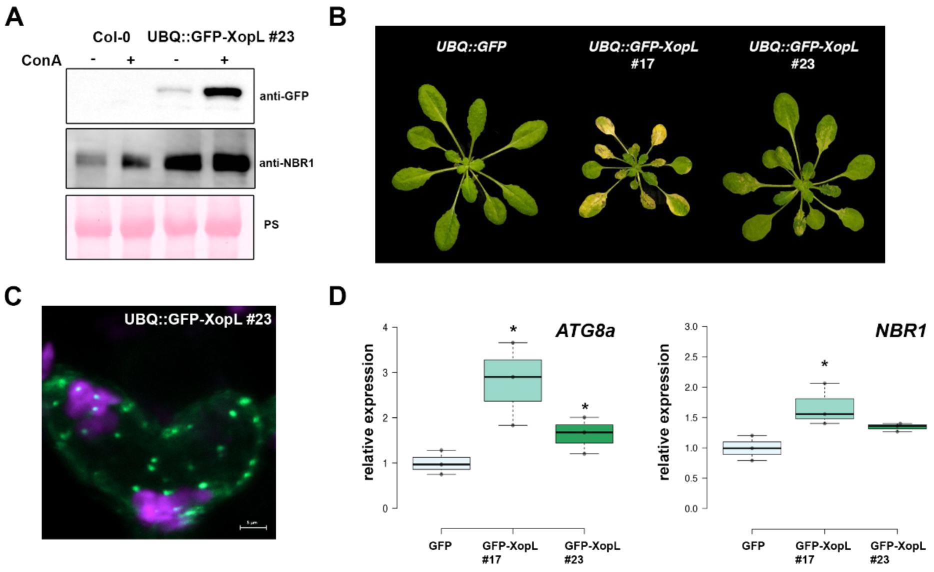

**Fig. S7: Transgenic *A. thaliana* GFP-XopL plants display defects in autophagic degradation**

**(A)** Immunoblot analysis of NBR1 protein levels in transgenic UBQ::GFP-XopL plants or Col-0. Plants were treated with concanamycin A (ConA) for 6 hours. Expression of GFP-XopL was verified with an anti-GFP antibody. Ponceau S staining serves as a loading control.

**(B)** 5 weeks old *A. thaliana* plants expressing UBQ::GFP-XopL develop an early senescence phenotype reminiscent of autophagy deficient mutants.

**(C)** Localization analysis of GFP-XopL of transgenic *A. thaliana* UBQ::GFP-XopL #23 line. Image represents single confocal planes from abaxial epidermal cells (bars = 5 µm).

**(D)** RT-qPCR analysis of *ATG8a* and *NBR1* transcript levels in *Arabidopsis thaliana* GFP or GFP-XopL plants. Values represent expression (n=3) relative to GFP control and were normalized to *PP2A*. Statistical significance (**P*<0.05) was revealed by Student's *t*-test.

**Supplemental Figure 8**

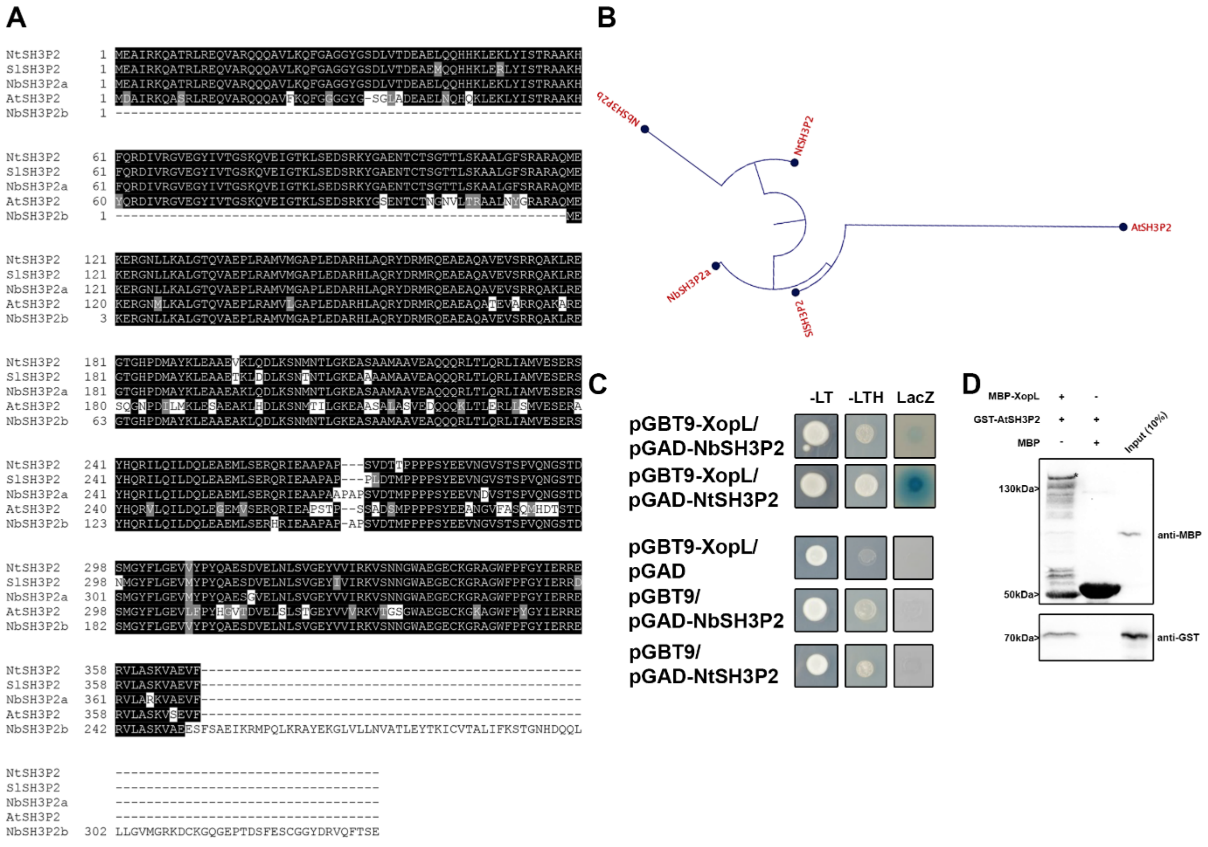


**Fig. S8: SH3P2 is conserved in different plant species.**

**(A)** Protein sequence alignment of SH3P2 from different species. The alignment was generated using CLUSTALW2 with default parameters and BoxShade 3.21. Positions of identical and similar sequences are boxed in black and grey, respectively. The following sequences were used to build the alignment: *Arabidopsis thaliana*, *Nicotiana tabacum, Nicotiana benthamiana*, *Solanum lycopersicum*.

**(B)** Phylogentic analysis of the SH3P2 from different plant species.

**(C)** XopL interacts with SH3P2 from *Nicotiana tabacum* and *Nicotiana benthamiana*. Interaction of XopL with SH3P2 in yeast two-hybrid assays. XopL fused to the GAL4 DNA-binding domain was expressed in combination with SH3P2 fused to the GAL4 activation domain (AD) in yeast strain Y190. Cells were grown on selective media before a LacZ filter

assay was performed. The empty AD or BD vector served as negative control. NtSH3P2 = *Nicotiana tabacum* SH3P2, NbSH3P2 = *Nicotiana benthamiana* –LT = yeast growth on medium without Leu and Trp, –HLT = yeast growth on medium lacking His, Leu, and Trp, indicating expression of the HIS3 reporter gene. LacZ, activity of the lacZ reporter gene.
**(D)** *In vitro* co-IP assay showing direct interaction of XopL with AtSH3P2. MBP-XopL and GST-AtSH3P2 were expressed in E. coli. Pull down was performed using amylose resin. Proteins were detected in an immunoblot using antibodies as indicated.

**Supp Video 1: XopL/SH3P2 puncta are mobile**
*Nicotiana benthamiana* leaf epidermal cells transiently expressing Venus$^{N173}$-XopL in combination with AtSH3P2-Venus$^{C155}$.

**Supplemental Figure 9**

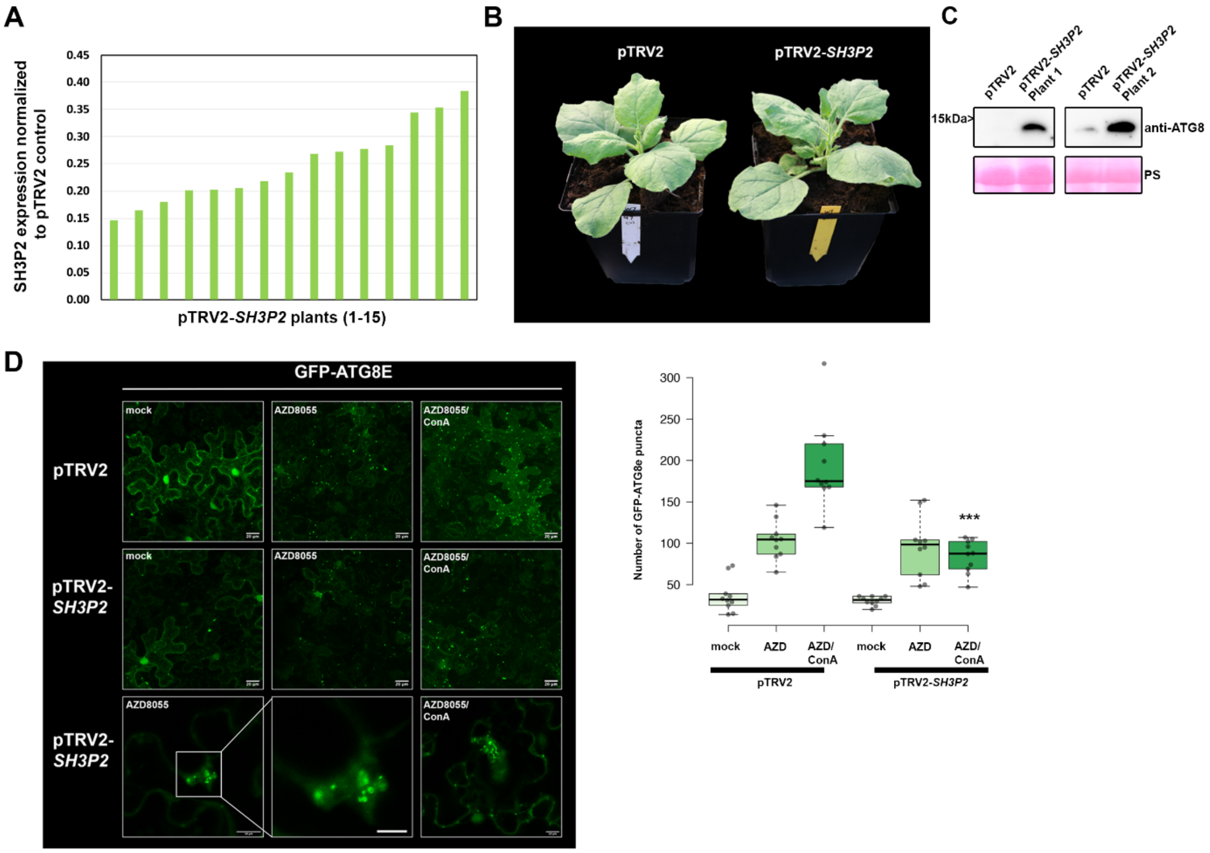


**Fig. S9: Silencing of SH3P2 in *N. benthamiana* perturbs autophagy.**
**(A)** qRT-PCR analysis of SH3P2 mRNA levels in silenced plants. *Actin* expression was used to normalize the expression value in each sample, and relative expression values were determined against pTRV2 control plants (set to 1).
**(B)** Phenotype of SH3P2-VIGS plants in comparison to the pTRV2 control. Picture was taken 14 dpi.
**(C)** Immunoblot analysis of ATG8 protein levels in *N. benthamiana* pTRV2 (control) and pTRV2-*SH3P2* (*SH3P2* silencing) 2 weeks after VIGS. Ponceau Staining (PS) served as a loading control. The experiment was repeated twice with similar results.
**(D)** GFP-ATG8e-labeled autophagosomes were quantified from pTRV2 or pTRV2-SH3P2 plants transiently expressing GFP-ATG8e at 2dpi in the presence of autophagy inducer AZD = AZD8055 and AZD/ConA. Puncta were calculated from z-stacks of *n*=10 individuals using

ImageJ. Statistical significance (*** $P < 0.5$) was revealed by Student's *t*-test comparing number of autophagosomes in AZD/ConA treatments in pTRV2 and pTRV2-*SH3P2* plants. The experiment was repeated twice with similar results.

### Supplemental Figure 10

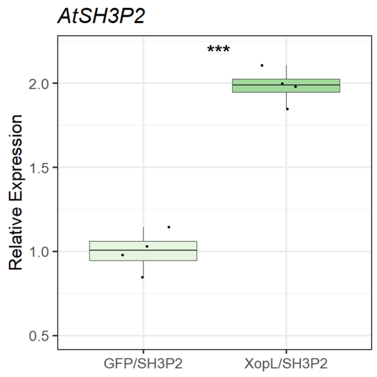


**Fig. S10: Gene expression of SH3P2 is induced by XopL and XopL-mediated degradation is due to post-transcriptional degradation events.**
**(A)** qRT-PCR analysis of AtSH3P2-specific mRNA levels in *N. benthamiana* plants transiently expressing AtSH3P2-HA, upon coexpression with GFP or GFP-XopL. *Actin* expression was used to normalize the expression value in each sample. Values represent *AtSH3P2* transcript level normalized to control (n=4).

### Supplemental Figure 11

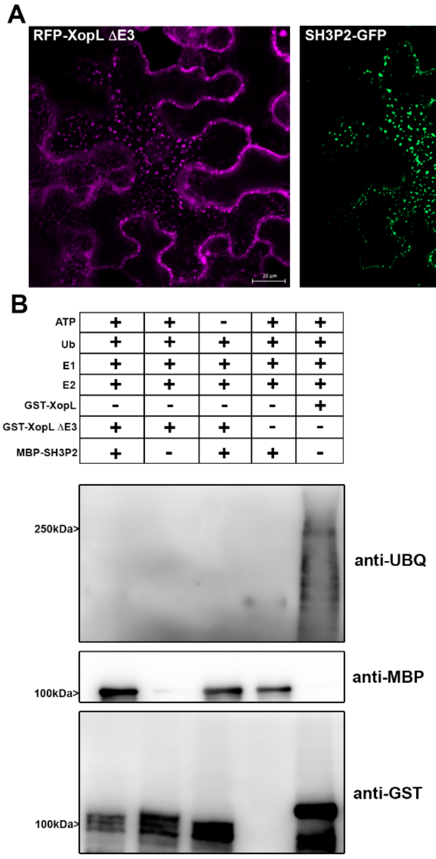


**Fig. S11: RFP-XopL ΔE3 co-localizes with and is unable to ubiquitinate SH3P2-GFP.**

**(A)** Colocalization analysis of RFP-XopL ΔE3 with SH3P2-GFP in *N. benthamiana* leaves. Imaging was performed 2 d after transient expression and images represent single confocal planes from abaxial epidermal cells (bars = 20 µm).
**(B)** *In vitro* ubiquitination assay reveals GST-XopL ΔE3 is unable to ubiquitinate MBP-SH3P2. GST-XopL, ΔE3 and MBP-SH3P2 were tested using the Arabidopsis His-AtUBA1 and His-AtUBC8. Lanes 2 to 4 are negative controls, while Lane 5 is the positive control with GST-XopL. Proteins were separated by SDS-PAGE and detected by immunoblotting using the indicated antibodies. The experiment was repeated three times with similar results.

**Supplemental Figure 12**

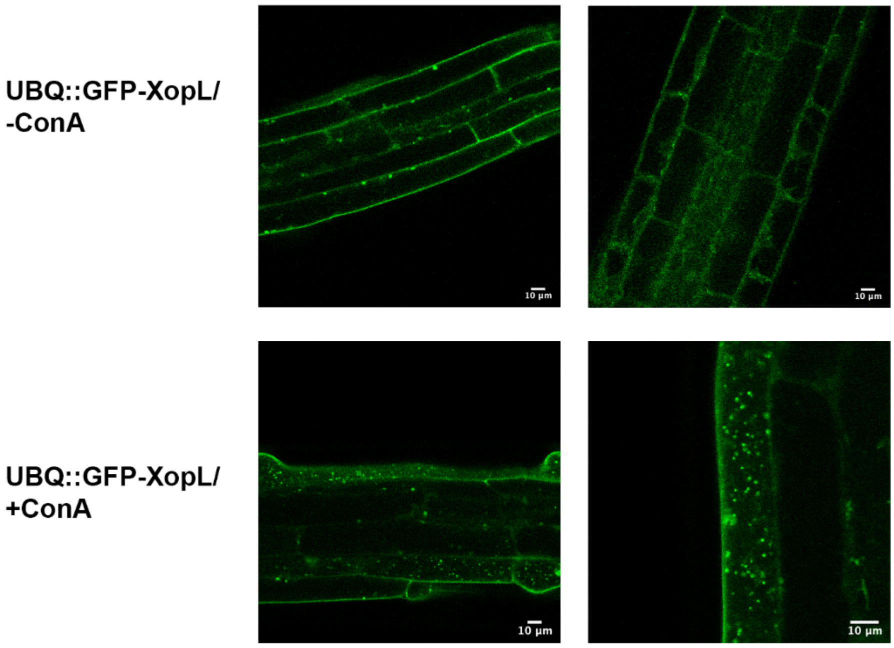


**Fig. S12: XopL is degraded in the vacuole**. Localization of GFP-XopL in the presence or absence of ConA in transgenic GFP-XopL. DMSO or 0.5 µM ConA was used to treat seedlings, followed by confocal imaging of the roots. GFP-labeled puncta detectable upon ConA treatment indicate XopL accumulation in the vacuole (bars = 20 µm).

**Supplemental Figure 13**

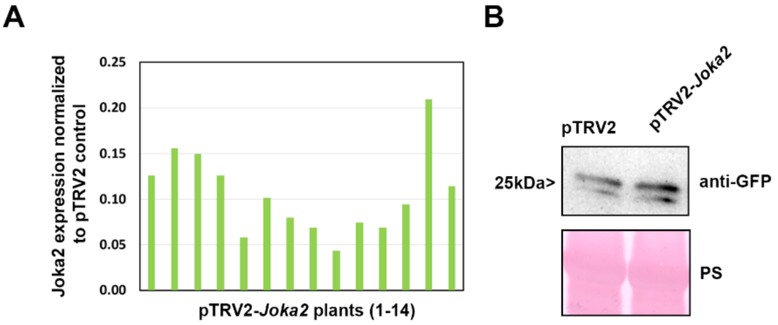


**Fig. S13: Virus-induced gene silencing of *Joka2* in *N. benthamiana* plants.**
**(A)** qRT-PCR analysis of Joka2 mRNA levels in *Joka2* silenced pepper plants. *Actin* expression was used to normalize the expression value in each sample, and relative expression values were determined against pTRV2 control plants (set to 1)**.**

**(B)** Immunoblot analysis of GFP and GFP-XopL protein levels in *N. benthamiana* plants silenced for *Joka2* (pTRV2-Joka2) compared against control (pTRV2). Ponceau staining (PS) served as a loading control.

**Supplemental Figure 14**

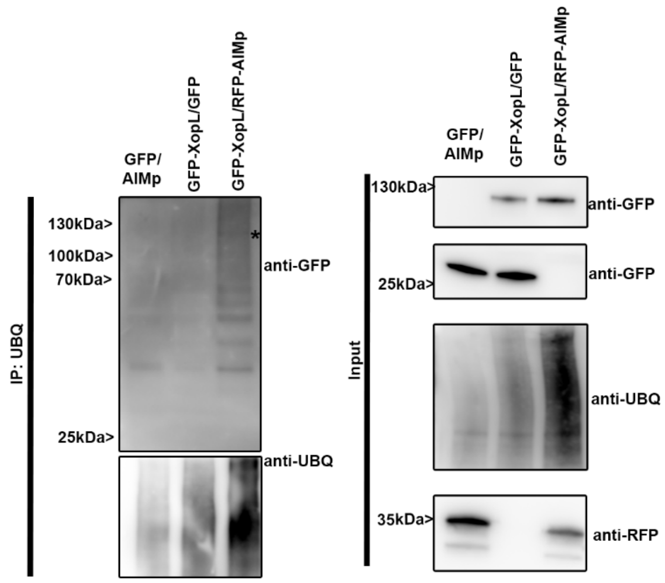


**Fig. S14: XopL in planta ubiquitination is enhanced by the presence of AIMp** GFP-XopL or GFP were transiently expressed in *N. benthamiana*. RFP-AIMp was co-infiltrated. Samples were taken 48 hpi, and total proteins (Input) were subjected to immunoprecipitation (IP) with the ubiquitin pan selector, followed by immunoblot analysis of the precipitates using either anti-GFP or anti-ubiquitin antibodies. RFP-AIMp expression was verified by an anti-RFP antibody. Asterisk indicates the GFP-XopL full-length protein. The experiment was repeated twice with similar results.

**Supplemental Figure 15**

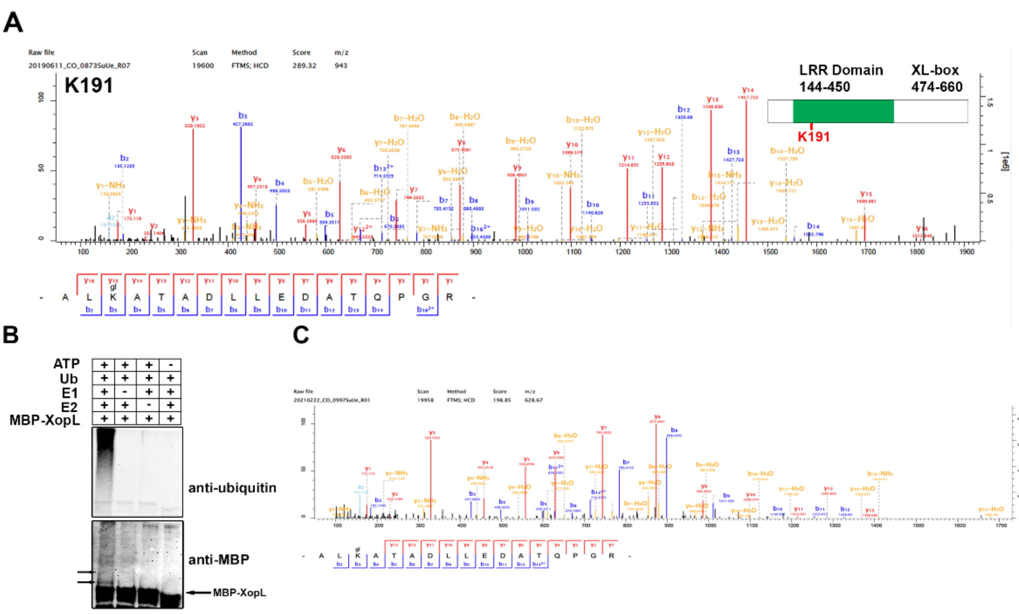


**Fig. S15: XopL is ubiquitinated *in planta* and undergoes self-ubiquitination**

**(A)** XopL ubiquitination site at lysine 191 was identified in vivo by LC-MS/MS. GFP-XopL was transiently expressed in *N. benthamiana* and total proteins were subjected to anti-GFP IP followed by trypsin digestion. Ubiquitinated peptides were detected by LC-MS/MS. The spectrum shows the fragmentation pattern of the GlyGly modified peptide ALglKATADLLEDATQPGR corresponding to amino acids 189-205.
**(B)** *In vitro* ubiquitination assay reveals autoubiquitination of XopL. Ubiquitination of MBP-XopL was tested using the Arabidopsis His-AtUBA1 and His-AtUBC8. Lanes 2 to 4 are negative controls. Proteins were separated by SDS-PAGE and detected by immunoblotting using the indicated antibodies. Arrows in the MBP blot indicate higher molecular weight bands of MBP-XopL and autoubiquitination events. The experiment was repeated twice with similar results.
**(C)** XopL ubiquitination site at lysine 191 was identified in vitro by LC-MS/MS. GST-XopL was used in an in vitro ubiquitination assay and samples were subjected to trypsin digestion. Ubiquitinated peptides were detected by LC-MS/MS. The spectrum shows the fragmentation pattern of the GlyGly modified peptide ALglKATADLLEDATQPGR corresponding to amino acids 189-205.

### Supplemental Figure 16

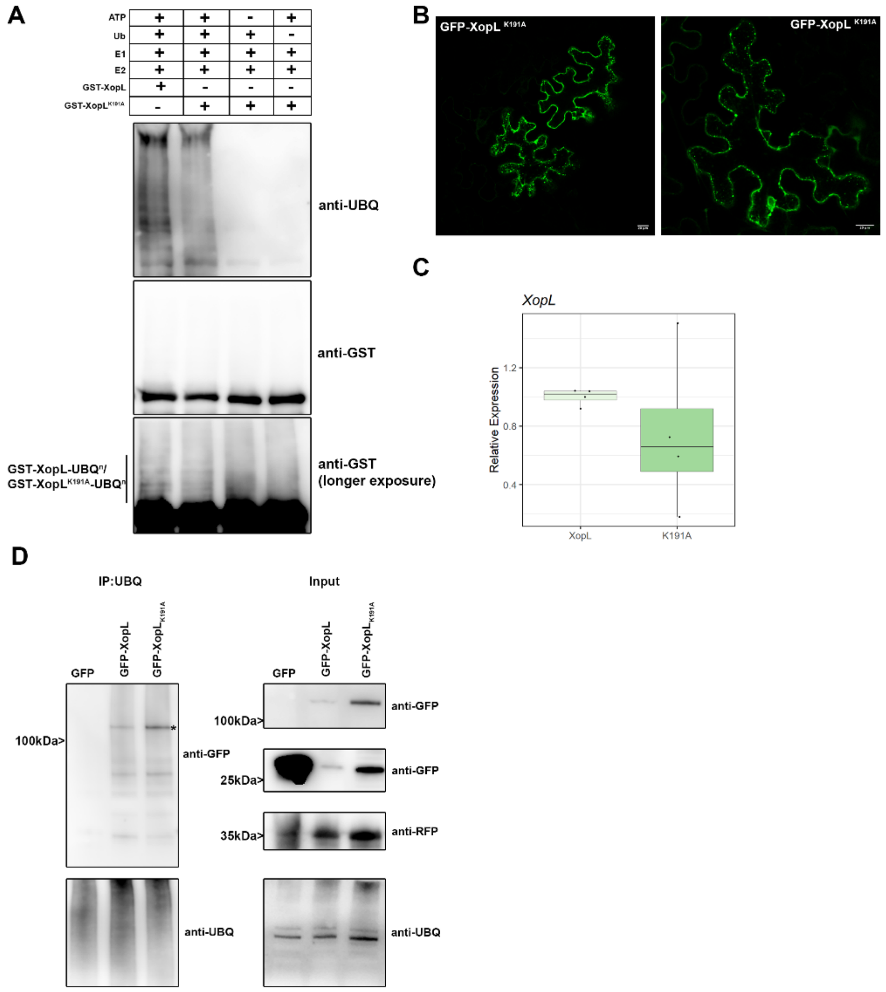


**Fig. S16: Characterization of XopL K191A variant in vitro and in planta.**
**(A)** *In vitro* ubiquitination assay reveals less autoubiquitination of XopL K191A compared to XopL WT. Ubiquitination of GST-XopL was tested using the Arabidopsis His-AtUBA1 and His-AtUBC8. Lanes 3 to 5 are negative controls. Proteins were separated by SDS-PAGE and

detected by immunoblotting using the indicated antibodies. The experiment was repeated twice with similar results.

**(B)** Localization analysis of GFP-XopL$_{K191A}$ in *N. benthamiana* leaves. Imaging was performed 2 d after transient expression and images represent single confocal planes from abaxial epidermal cells (bars = 20 µm).

**(C)** qRT-PCR analysis of GFP-XopL and GFP-XopL K191A mRNA levels in *N. benthamiana* plants during transient expression. *Actin* expression was used to normalize the expression value in each sample.

**(D)** GFP-XopL and GFP-XopL$_{K191A}$ were transiently expressed in *N. benthamiana*. RFP-AIMp was co-infiltrated. Samples were taken 48 hpi, and total proteins (Input) were subjected to immunoprecipitation (IP) with the ubiquitin pan selector, followed by immunoblot analysis of the precipitates using either anti-GFP or anti-ubiquitin antibodies. GFP served as a control. RFP-AIMp expression was verified by an anti-RFP antibody. Asterisk indicates the GFP-XopL full-length protein. The experiment was repeated twice with similar results.

**Supplemental Figure 17**

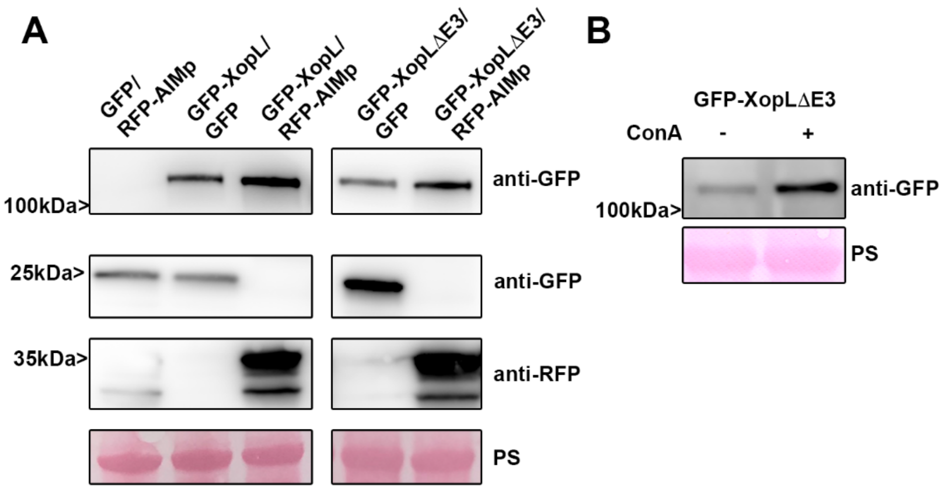


**Fig. S17: XopL ΔE3 is degraded by autophagy.**

**(A)** GFP, GFP-XopL, or GFP-XopL ΔE3 was coexpressed with RFP-AIMp or RFP control in *N. benthamiana*. Samples were taken at 2 dpi, total proteins extracted and immunoblotted using the indicated antibodies. Ponceau Staining (PS) served as a loading control.

**(B)** Immunoblot of transiently expressed GFP-XopL ΔE3 in *N. benthamiana* after treatment of ConA or DMSO carrier. Ponceau Staining (PS) served as a loading control.

**Supplemental Figure 18**

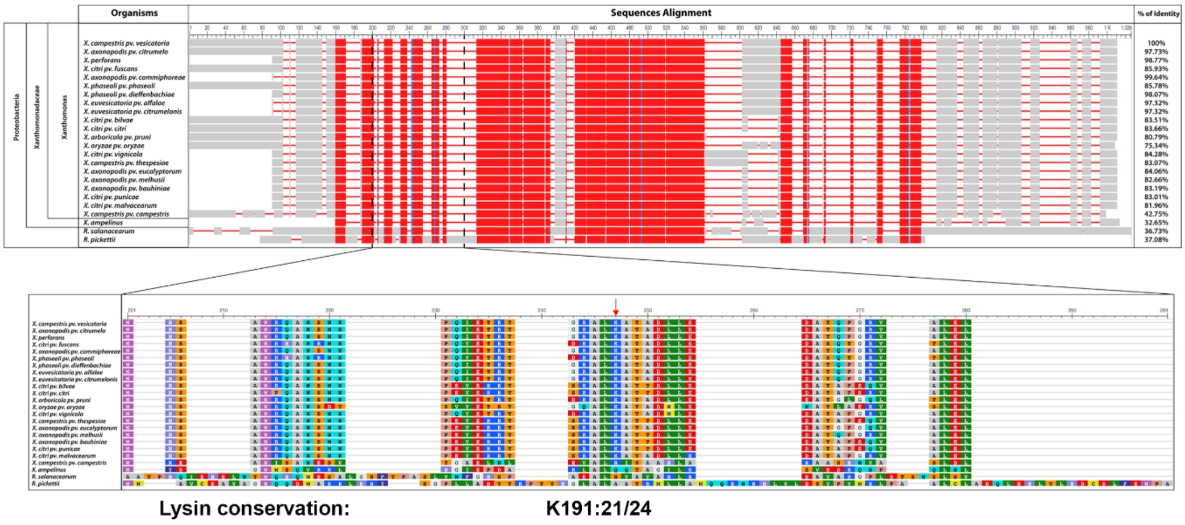


**Fig. S18: XopL K191 residue is highly conserved through the Xanthomonas genus**

**(A)** Sequence alignment of XopL protein from *Xcv* with related effectors from *Xanthomonas* genus or more distantly related plant pathogen bacteria. Colors represent amino acid conservation through the alignment, with red for highly conserved residues, blue for lower conservation and grey for no conservation. Identical amino acid percentage to XopL$^{Xcv}$ is displayed to the right of the alignment.

**(B)** Closer view of the region 201-299 of the alignment with amino acids colored according to the Rasmol coloration. Lysine K191 is indicated with a red arrow.
